# Site-Specific Introduction of Non-Canonical Amino Acids into natural and engineered Non-Ribosomal Peptides

**DOI:** 10.64898/2026.07.12.738027

**Authors:** Max Schreiber, Maryam Dehghan, Shadrack Kibet, Marie Tvilum, Carsten Kegler, Kevin Hoffmann, Peter Grün, Sven Balluff, Karsten Siems, Helge B. Bode

## Abstract

The incorporation of non-canonical amino acids (ncAAs) into proteins, developed in the past 20 years, has opened new avenues with respect to protein structure, protein modification, protein-protein interaction or enzyme catalysis beyond what is possible with the 20 proteinogenic AAs. Although >300 unusual building blocks including several ncAAs have been described in nonribosomal peptides (NRPs) naturally, we aimed to further expand the scope of the underlying nonribosomal peptide synthetases (NRPS) to incorporate ncAAs beyond the naturally available ones. We have therefore systematically screened for ncAA accepting NRPS systems, applied NRPS engineering to transfer the respective ncAA-accepting parts into other NRPSs and thereby created novel peptides that were further derivatized in post-enzymatic chemical synthesis reactions directly in bacterial culture extracts.

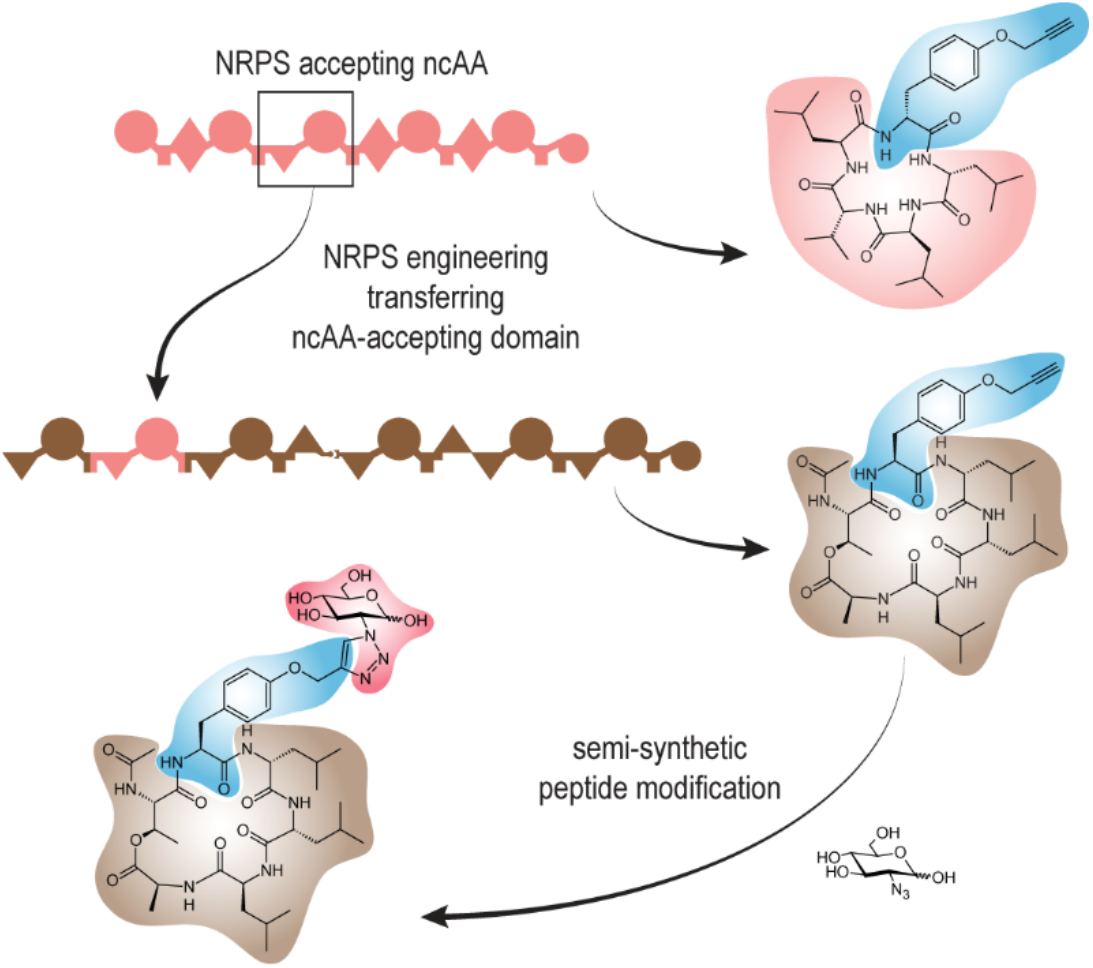

## Introduction

Many small molecule drugs in use today are natural product-related compounds^1^. For pharmaceutical use, natural products are often further derivatized resulting in semi-synthetic drugs^2,3^ like antibiotics of the penicillin family^4^. Such derivatizations are often needed to improve bioactivity, to adjust the pharmacokinetics of a compound or to overcome resistance against existing drugs^5–8^.

A variety of clinically used drugs are peptides produced by so-called non-ribosomal peptide synthetases (NRPS)^1,3^ e.g., the immune-suppressive drug cyclosporin^9^, the anti-cancer drug romidepsin^10^, or antibiotics like vancomycin^11^ and daptomycin^12^ and therefore the modification of these NRPS is a major goal of synthetic biology in the natural product biosynthesis field. During the last years, numerous methods for NRPS engineering have been developed enabling their targeted or untargeted modification and thus leading to the production of backbone-modified peptide or even hybrids of peptides and polyketides.

What has not been used in NRPS engineering extensively is the natural ability of the NRPS systems to incorporate amino acids (AAs) far beyond the 20 proteinogenic ones, despite >300 different AA and fatty acid building blocks have been found in natural non-ribosomal peptides (NRP)^13,14^. Although there is increasing knowledge about unusual building blocks generated by post-translational modifications of the 20 proteinogenic AAs in the field of ribosomally generated and post-translationally modified peptides (RiPPs)^15^, ribosomally and non-ribosomally peptides greatly differ in the overall number of already known building blocks.

To overcome this limitation of ribosomally derived peptides and proteins, genetic code expansion (GCE) methods have been developed as a powerful approach to evolve suitable amino acyl tRNA synthetases (aaRS) that allow efficient incorporation of even AAs very unrelated to the 20 proteinogenic ones^16^, opening completely new avenues for shaping structures and properties of such modified peptides and proteins. The starting point of this field was the identification of natural aaRS systems that naturally incorporate AA-derivatives like pyrrolysine into proteins^17,18^.

Although in natural product research, incorporation of non-canonical building blocks have been used in precursor-directed biosynthesis (PDB) or mutasynthesis campaigns to obtain desired derivatives^19,20^, such approaches have not been systematically evaluated to incorporate ncAAs into NRPS-derived peptides and to understand the natural substrate promiscuity of NRPS systems towards non-natural ncAAs. However, it has been shown previously that NRPS systems can indeed be evolved to incorporate such ncAAs preferentially over the natural AA^21–23^.

Encouraged by these results, we systematically analyzed the ability of natural NRPS systems to accept such ncAAs naturally. We used the parallel cultivation of strains producing selected NRPs, followed by molecular network analysis of these cultures containing different ncAAs in the growth medium, in order to identify those NRPS parts able to incorporate the desired ncAAs. The identified natural NRPS parts were then incorporated into other NRPS systems showing their transferability and enabling the targeted modification of the produced peptides at the peptide backbone as well as further derivatization of the incorporated reactive moieties.

## Results

### Workflow establishment

To identify promiscuous A-domains that can be used for later NRPS engineering, precursor-directed biosynthesis (PDB) of different non-ribosomal peptides (NRP) was performed by supplementing desired building blocks to the production medium. Upon their incorporation, NRP derivatives can be generated without prior knowledge about the biosynthesis and the structure of the A-domains^24^. Figure 1A gives a schematic overview of the workflow for rapid identification of promiscuous A-domains using PDB and feature-based molecular network analysis, an untargeted data analysis workflow that organizes MS^2^ spectra into a map based on their spectral similarities^25,26^. A given NRP-producing strain was cultured and supplemented with one specific ncAA, including a non-producing negative control and a positive control without any supplementation. All cultures were extracted individually and measured by high-performance liquid chromatography high-resolution mass spectrometry (HPLC-HR-MS) and a combined feature-based molecular network was generated using GNPS^26^. Ten NRPS from seven different strains from the genera *Xenorhabdus* and *Photorhabdus* (Table 1) were subjected to PDB by supplementing a selection of 19 ncAA (Figure 1B) to analyze them for promiscuous A-domains. These entomopathogenic Gram-negative proteobacteria^27–29^ were selected because they harbor many NRPS, which often produce more than one variant due to natural A-domain promiscuity^30–38^. To decrease the complexity of the extracts to be analyzed, the NRPS gene cluster of interest is artificially activated by promotor exchange^39–41^. Additionally, the *hfq* gene, encoding for a small RNA chaperone, is deleted (Δ*hfq*) in those strains, which causes the loss of most other secondary metabolites known as easyPACId approach^40,42^. This workflow was validated using the known promiscuity of the third A-domain of the GameXPeptide synthetase (GxpS) from *P. laumondii* known to incorporate *para*-substituted phenylalanine derivatives (Figure S2, Table S12)^43^.

**Table 1.** Overview of all NRPS and the strains from which they originate that have been subjected to precursor-directed biosynthesis (PDB) in this work.

| NRPS | Strain |
| --- | --- |
| GameXPeptide synthetase (GxpS)<sup>32,37</sup> | *Photorhabdus laumondii* TTO1 |
| szentiamide synthetase (SzeS)<sup>63–65</sup> | *Xenorhabdus szentirmaii* DSM 26338 |
| taxlllaid synthetase (TxIAB)<sup>35</sup> | *X. indica* DSM 17382 |
| fitayylide synthetase (FitAB)<sup>48</sup> | *X. innexi* DSM 16336 |
| xeneprotide synthetase (XnpAB)<sup>38</sup> | *X. stockiae* KJ12.1 |
| xenortide synthetase (XndAB)<sup>31,63,66,67</sup> | *X. nematophila* ATCC 19061 |
| GameXPeptide synthetase (GxpS)<sup>38</sup> | *X. doucetiae* DSM 17909 |
| protegomycin synthetase (PrtAB)<sup>68</sup> |  |
| xenoamicin synthetase (XabABC)<sup>34</sup> |  |
| peptide-antimicrobial-*Xenorhabdus* synthetase (PaxABC)<sup>30,38,44,45</sup> |  |

**Figure 1.**
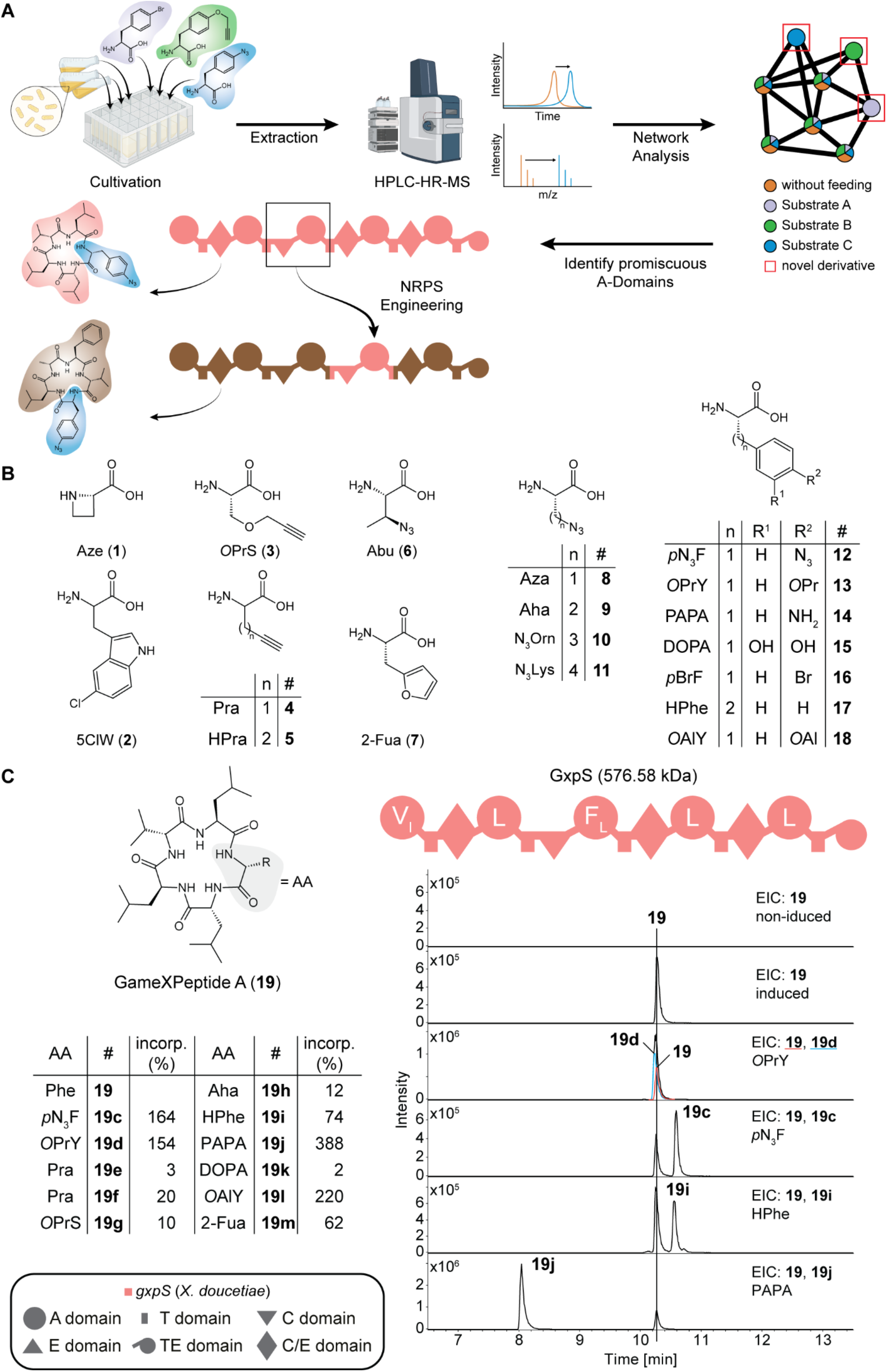
Identification of promiscuous A-domains. **A.** Workflow: The combination of plate-based precursor-directed biosynthesis with feature-based molecular network analysis allows rapid identification of NRP derivatives and identification of promiscuous A-domains. **B**. Overview of selected ncAA for precursor-directed biosynthesis. *O*Pr: propargyl-; *O*Al: allyl-(also see Figure S1). **C**. Schematic depiction of the NRPS GxpS from *X. doucetiae* DSM 17909 and produced GameXPeptide derivatives **19c**-**19m** identified by supplementing the production medium with indicated non-cognate amino acids. Extracted ion chromatograms (EIC) of compounds **19, 19d, 19c, 19j**, and **19**. A-domain specificity is indicated within the A domains by the amino acid one-letter code.

### Generation of NRPs incorporating ncAAs

The GameXPeptide (GXP) synthetase GxpS from *X. doucetiae* was subjected to precursor-directed biosynthesis by supplementing the production medium with 13 different substrates (**3**-**9, 7, 12**-**15, 17, 18**; Figure 1B) of which 11 were incorporated leading to novel GXP derivatives (Figure S3-S6, Table S13). Figure 1C shows the domain architecture of GxpS together with its product **19** and a selection of well-produced derivatives (**19d, 19c, 19i**, and **19j)**. Their production compared to **19** can be seen in the extracted ion chromatograms (EICs) in Figure 1C. All exchanges took place at module three, rendering the A3-domain of GxpS an A-domain with extreme relaxed substrate specificity. The A3-domain was able to incorporate various azide and alkyne building blocks. It accepted *O*PrS (**3**), Pra (**4**), Hpra (**5**), Aha (**9**), *p*N_3_F (**12**), and *O*prY (**13**) (Figure 1C, Figure S1, S3-S6, Table S13). Comparing their production relative to **19**, with 164% (**19c**) and 154% (**19d**), the aromatic **12** and **13** worked more efficient than aliphatic azides and alkynes (Table S13). The azides Aza (**8**) and Abu (**6**) were not incorporated (Figure 1C, Figure S1). In addition to azides and alkynes, also the diene 2-Fua (**7**) and the dienophile *O*alY (**18**), which can be used in Diels-Alder reactions, were incorporated with relative incorporation rates of 62 (**19m**) and 220% (**19l**), respectively (Figure S6, Table S13). Compared to the incorporation rate of *para*-substituted phenylalanine derivatives, the *meta*-substituted DOPA (**15**) was incorporated poorly with only 2% (**19k**; Table S13). Relative production rates are based on peak areas of the derivatives compared to the area of natural **19**.

Similarly, incorporation of ncAAs into NRPS systems producing the depsipeptides szentiamide, xenoamicin, xeneprotide, taxlllaid and fitayylide, as well as the linear peptide xenortide (Table 1) was investigated (Figure 2) and in all cases, incorporation of some ncAAs was observed (Figure S7-S22). In a few cases much better as the canonical AA (see the 12-fold better incorporation of **1** into xeneprotide (**27**) compared to natural Pro (Figure S13)). While ncAA incorporation could also be observed for protegomycin (Figure S24) carrying multiple Tyr residues, the promiscuous position(s) could not be fully confirmed (Figure S25) and therefore it was excluded from further work. Further details about the incorporation results for the different peptide classes are described in the supplementary information.

**Figure 2.**
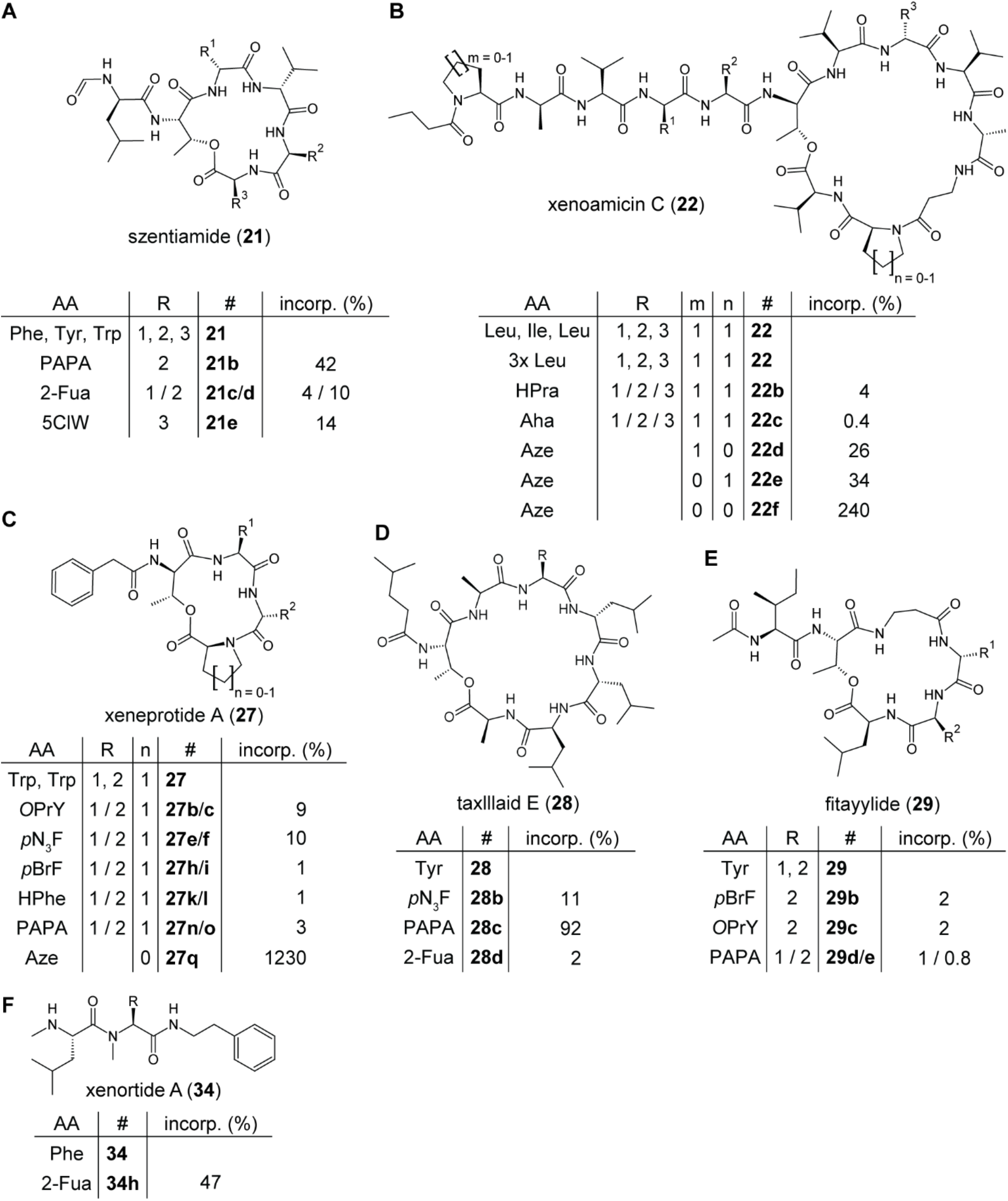
Overview of NRP derivatives generated by precursor-directed biosynthesis. Shown are szentiamide derivatives **21b**-**21e** (**A**) produced by the NRPS SzeS, xenoamicin derivatives **22b**-**22f** (**B**) produced by XabABC, xeneprotide derivatives **27b**-**27q** (**C**) produced by XnpAB, taxlllaid derivatives **28b**-**28d** (**D**) produced by TxlAB, fitayylide derivatives **29b**-**29e** (**E**) produced by FitAB, and the xenortide derivative **34h** (**F**) produced by XndAB. Modular organization of all NRPS is shown in Figure S7. For references of all analyzed peptides, see Table 1.

The PAX synthetase PaxABC from *X. doucetiae* DSM 17909 consisting of three proteins and producing the highly cationic, cyclic lipopeptide PAX (peptide-antimicrobial-*Xenorhabdus*, **35**, Figure S23)^30,44,45^ was the only NRPS system not accepting any of the ncAAs tested (**6, 8**-**11**; Figure 1B). However, it is almost exclusively composed of Arg and Lys residues of which no ncAA analogs have been used in the PDB.

In summary, nine out of ten NRPS harbored at least one promiscuous A-domain generating a total of 42 unique NRP derivatives (Table S21), excluding double- or multiple incorporations of ncAA. Thus, the presented microtiter plate-based workflow, combining precursor-directed biosynthesis and feature-based molecular network, is a powerful tool for the *in vivo* generation of NRP derivatives and the identification of promiscuous A-domains.

### Transfer of ncAA-incorporating A domains into model NRPS

To utilize the identified, promiscuous A-domains for the generation of synthetic hybrid NRPS that produce new peptides harboring ncAA, a recently developed Golden Gate-based NRPS engineering method was used^46,47^. It combines Golden Gate cloning with the XUT^IV^ (e**X**change **U**nits inside **T**-domains) concept^48^ to allow scarless assembly and reusability of the part collection. This allows the rapid generation of hybrid NRPS^46,47^. Nine of the 15 identified, promiscuous A-domains were used to generate synthetic NRPS (Figure 3A). For this, the respective A-domains were cloned into Golden Gate donor vectors as XUT^IV^ modules^48^ and introduced into an acceptor generating nine trimodular, hybrid NRPS (NRPS-3 to NRPS-11). Except for NRPS-4, all were functional and heterologously produced the lipopeptides **37**-**40** (Figure S26-S29). From seven of those hybrid NRPS subjected to PDB with a focus on azides and alkynes, four were able to incorporate ncAAS into their respective lipopeptide (Table S22; Figure S30-S34). NRPS-3, harboring the phenylalanine-specific A3-domain of GxpS from *X. doucetiae*, produced the lipopeptides **37b** and **37c** at relative yields of 103 and 199% compared to **37** when supplemented with *O*prY (**13**) and *p*N_3_F (**12**), respectively (Figure 3B, C; Figure S30, S31).

**Figure 3.**
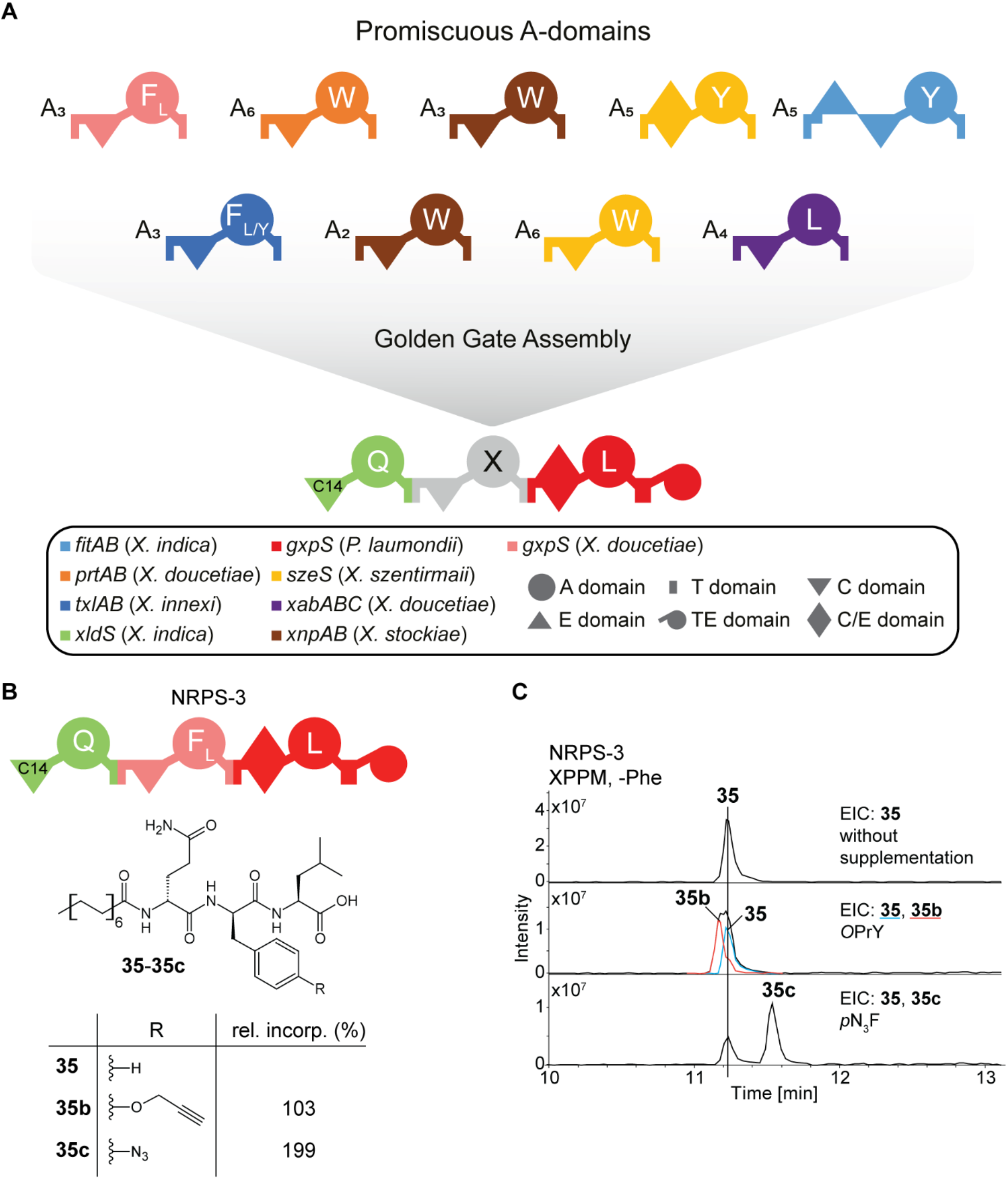
Transferability of promiscuous A domains using Golden Gate-based NRPS engineering^46,47^. **A.** Selection of ten identified promiscuous A-domains from eight different NRPS used as XUT^IV^ donor modules^48^. Golden Gate assembly of the donors with the acceptor results in the synthetic hybrids NRPS-11 to NRPS-20. Natural A-domain specificity indicated by the amino acid one-letter code. **B**. Schematic depiction of synthetic hybrid NRPS-11 and structures of produced lipopeptides **35**-**35c** together with their relative production compared to **35** produced by precursor-directed biosynthesis with *E. coli* DH10B::*mtaA*^69^ heterologously expressing *NRPS-11* in XPP media without phenylalanine (-Phe). Extracted ion chromatograms (EIC) of **35**-**35c** produced in XPPM, -Phe supplemented with **12** (*p*N_3_F) or **13** (*O*PrY).

### Targeted ncAA-incorporation into natural peptides

PDB of fitayylide derivatives showed that the tyrosine-specific A5-domain has moderate promiscuity (Figure 2E, Table S18; Figure S18) and was able to incorporate *O*prY (**13**) yielding 2% of **30b** (Table S18; Figure S18-S21, S35), while the azide *p*N_3_F (**12**) was not accepted at all (Figure 4A, S35, Table S18). In order to generate fitayylide derivatives that harbor **12**, a synthetic hybrid NRPS was constructed using the promiscuous A3-domain of GxpS from *X. doucetiae*, which accepts **12** in its natural NRPS (Figure 1C) as well embedded in the model NRPS-3 (Figure 3B, C). Indeed, fitayylide derivatives **41** with much higher incorporations rates of 7 (**41c**) and 22% (**41b**) for **12** and **13** could be achieved, respectively. However, since the inserted XUT fragment also changed the E-^D^C_L_ didomain into an ^L^C_L_ domain, the produced fitayylide variant showed an *S* configuration for the first Tyr. This change in configuration of the overall peptide chain enabled the TE domain to form a larger ring between the acyl-OH and the C-terminal Leu, leading to a cycle made of seven instead of five building blocks (**42**), but with the expected incorporation of the two ncAAs (**42b** and **42c**) in very good yields of 40 and 105% (Figure 4B, C, S36).

**Figure 4.**
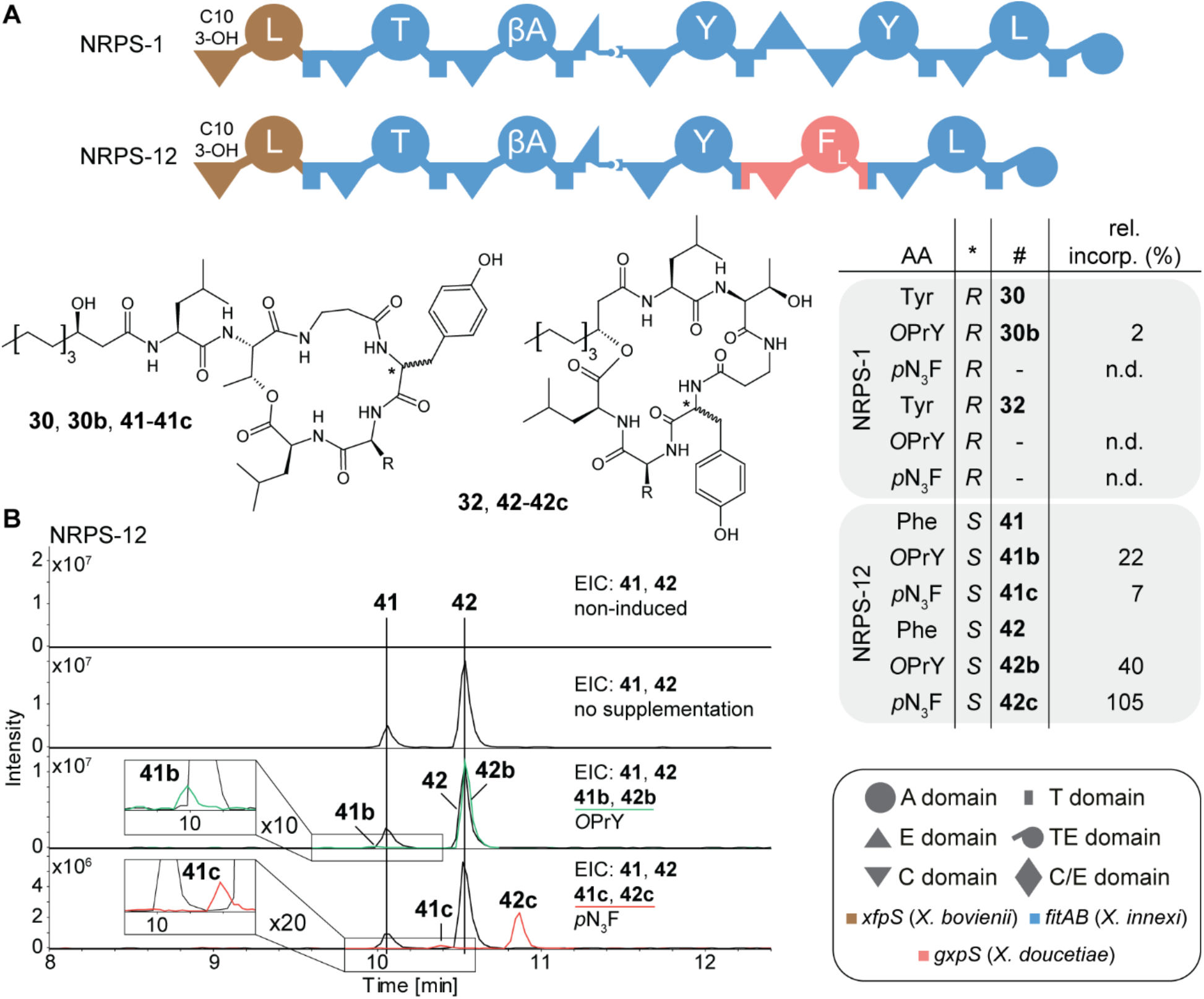
Increasing ncAA incorporation rates using NRPS engineering. **A.** Schematic depiction of NRPS-1 and NRPS-12 and the structure of produced peptides **30, 32, 41**, and **42**. Precursor-directed biosynthesis with **12** (pN_3_F) and **13** (OprY) resulted in no or poor incorporation for NRPS-1 (**30b**) but were well accepted by NRPS-12 (**41b, 41c, 42b, 42c**). The synthetic hybrid NRPS-12 uses the promiscuous A3 domain of GxpS. βA: beta-alanine. **B**. Extracted ion chromatograms (EIC) of **41**-**41c** and **42**-**42c** produced by NRPS-12 in precursor-directed biosynthesis. For NRPS-1 data, see Fig. S35.

This A-domain was further used in five other hybrid NRPS producing twelve additional lipopeptides (**50, 52**-**55, 58, 59a**/**b, 62**-**63**, and **65**; see structures) as well as six truncated peptides (**51, 56, 57, 60, 61**, and **66**; Figure S39-S51). In all five cases, the GxpS A-domain was able to incorporate **12** and **13** into the respective lipopeptides with a median relative incorporation rate of 59 and 55%, respectively (Figure S39-S51). For NRPS-16, which harbors this module twice, it was possible to incorporate **12** and **13** twice (**58g**/**58d**) or, when supplementing both simultaneously, both into the same peptide (**58h**/**i**; Figure S44B, C, Figure S46). This underlines the flexibility and compatibility of this module for NRPS engineering rendering it a valuable tool for NRP-derivatization.

Finally, we tested whether the modified peptides could be further chemically modified due to their reactive handles. Indeed, a crude culture extract containing **19m** carrying 2-Fua (**7**) reacted with *N*-phenyl-maleimide (**67**) in the expected Diels-Alder reaction as previously described for 3-Fua containing peptides (Figure 5A)^49,50^ but at a low yield due to non-optimized reaction conditions. Furthermore, a crude culture extract containing **50b** derived from PDB of NRPS-14 (Figure S39, S40) with **13** could be quantitatively reacted with 2-azido-2-desoxy-*D*-glucopyranose (N_3_Gluc, **69**) to the respective triazoles via copper-based Click-chemistry (Figure 5B, Figure S52). We could also show that a synthetic peptide **58h**, which was produced in small amounts when both **12** and **13** were fed to NRPS-16 (Figure S44), can even be cyclized (Figure S53). While this offers the possibility to introduce ring structures in parts of the peptide currently not accessible by biocatalysis, the reaction must probably be further optimized with respect to the ring size and configuration of the peptide chain as dimerization was observed as well. It has been reported previously that optimal conditions for chemical peptide macrocyclization are hard to predict and that there is no rule whether protecting groups are necessary or the beneficial effect of resin-bound peptide is needed^51^.

**Figure 5.**
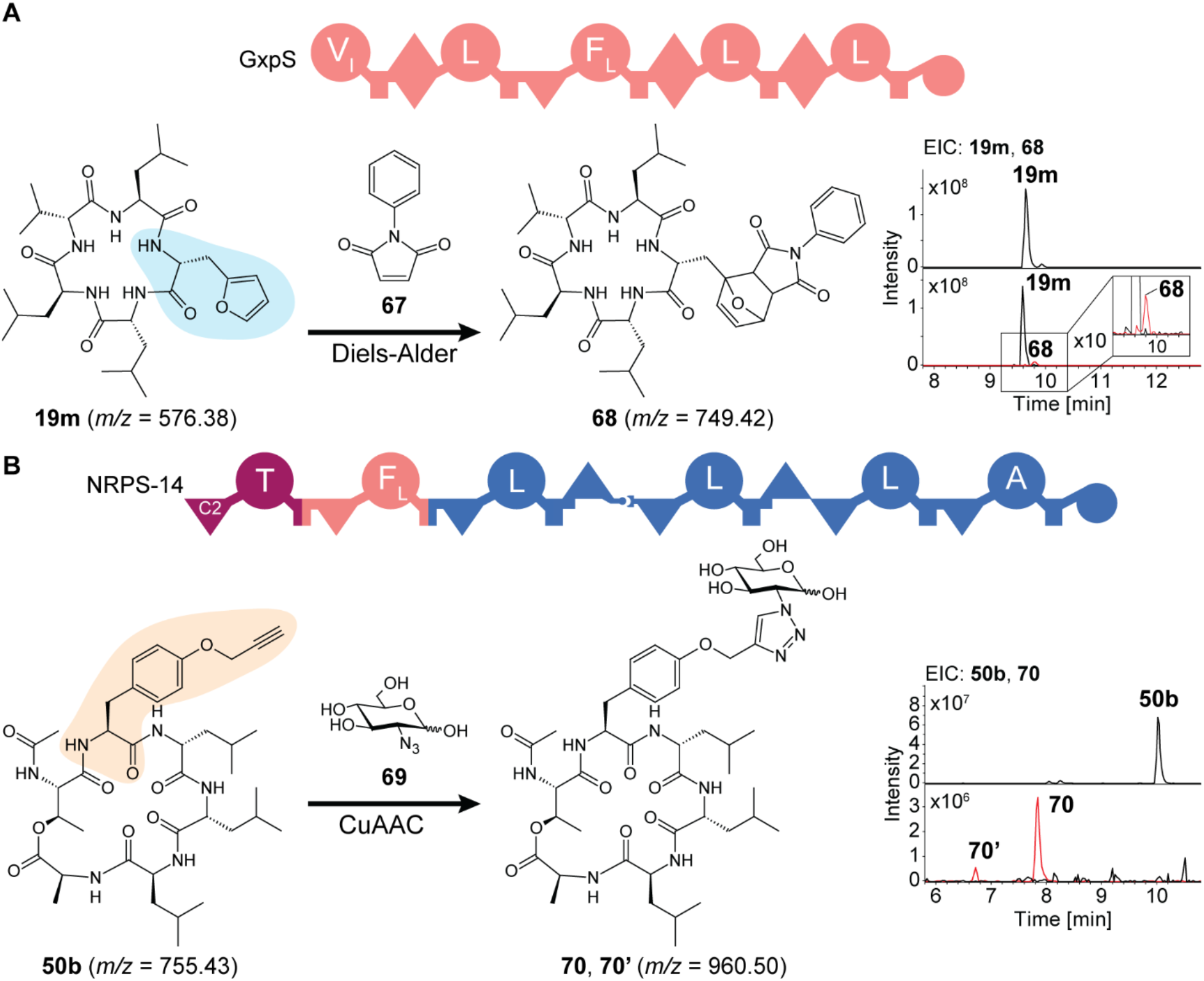
Chemical modification of cyclic peptides harboring ncAAs directly in crude extracts obtained from heterologous expression of selected NRPS in *E. coli*. **A.** The 2-Fua containing GXP-derivative (**19m**, black chromatogram) was modified with the dienophile *N*-phenylmaleimide (**67**) using a Diels-Alder reaction to give **68** (red chromatogram). **B**. The engineered alkyne-containing depsipeptide (**50b**, black chromatogram) was glycosylated quantitatively with 2-azido-2-desoxy-*D*-glucopyranose (**69**) in a copper-catalyzed azide-alkyne cycloaddition (CuAAC) to give **70**/**70’** (red chromatogram).

## Conclusions

We have shown that natural NRPS systems are able to accept ncAAs as substrates leading to backbone-modified peptide variants and that these features can be applied as tool in NRPS engineering campaigns. Probably many more NRPS parts show such a large promiscuity also for other ncAAs but have not been detected yet. The workflow described here (Figure 1) would allow their rapid identification. Furthermore, our results suggests that it should be possible to further increase the specificity of the respective A domains towards the ncAA applying targeted A-domain engineering as it has been shown recently for the optimized incorporation of 3-Fua^49^ or **13**^52^ or by high-throughput screening as shown for the incorporation of β-Phe^23^. Since also the condensation domain can be optimized to accept non-canonical building blocks^53^, future peptides derived from engineered NRPS systems might go way beyond what is currently known. Especially, for *de novo* design of peptide-based pharmaceuticals similar to orally available peptides like enlicitide^54^, rare non-cognate or non-natural amino acids (ncAA) are important to improve their pharmakokinetics^55^ as they can directly shape the properties of a compound or act as handles for later modification in a semi-synthetic approach. Applying engineered NRPS pathways incorporating such ncAAs might allow an even more biological production process than the chemo-enzymatic approach recently described for enlicitide production^56^.

Our work also gives additional insights into the role of thioesterases (TEs) responsible for release of the linear or cyclic peptide from the NRPS enzyme^57^. The FitB-TE domain allows cyclizations of the terminal Leu-6 carboxylic acid with the Thr-2 side chain as in NRPS-1 (WT) or the β-hydroxy moiety of the C10 acyl starter unit as in NRPS-12 (Figure 4). Both peptide chains differ only in the configuration of Tyr-4, far away from both ring-closing moieties. Most likely, different TEs show different substrate specificity rules for cyclization that must be studied in more detail in the future to apply them as versatile cyclization catalyst.

ncAA incorporation can also be applied to achieve post-NRPS peptide modifications like glycosylations (see Figure 5) for enhanced pharmacokinetics^58^, drug distribution, stability or crossing the blood-brain-barrier^59^. Even ring topologies beyond the natural N-terminal or side-chain to C-terminal cyclization might be possible (see Figure S53) and might even offer the generation of bicyclic systems, which are frequently found in the RiPP world^60,61^ and have been shown to have beneficial pharmacological properties^62^. In summary, there is a clear need for A-domains that are able to activate non-canonical amino acids and are compatible to NRPS engineering approaches.

## Acknowledgements

Work in the Bode lab was supported by the Max Planck Society. The authors are grateful to Edna Bode, Petra Happel, and Weihua Jiao for providing promoter exchange mutants.

---

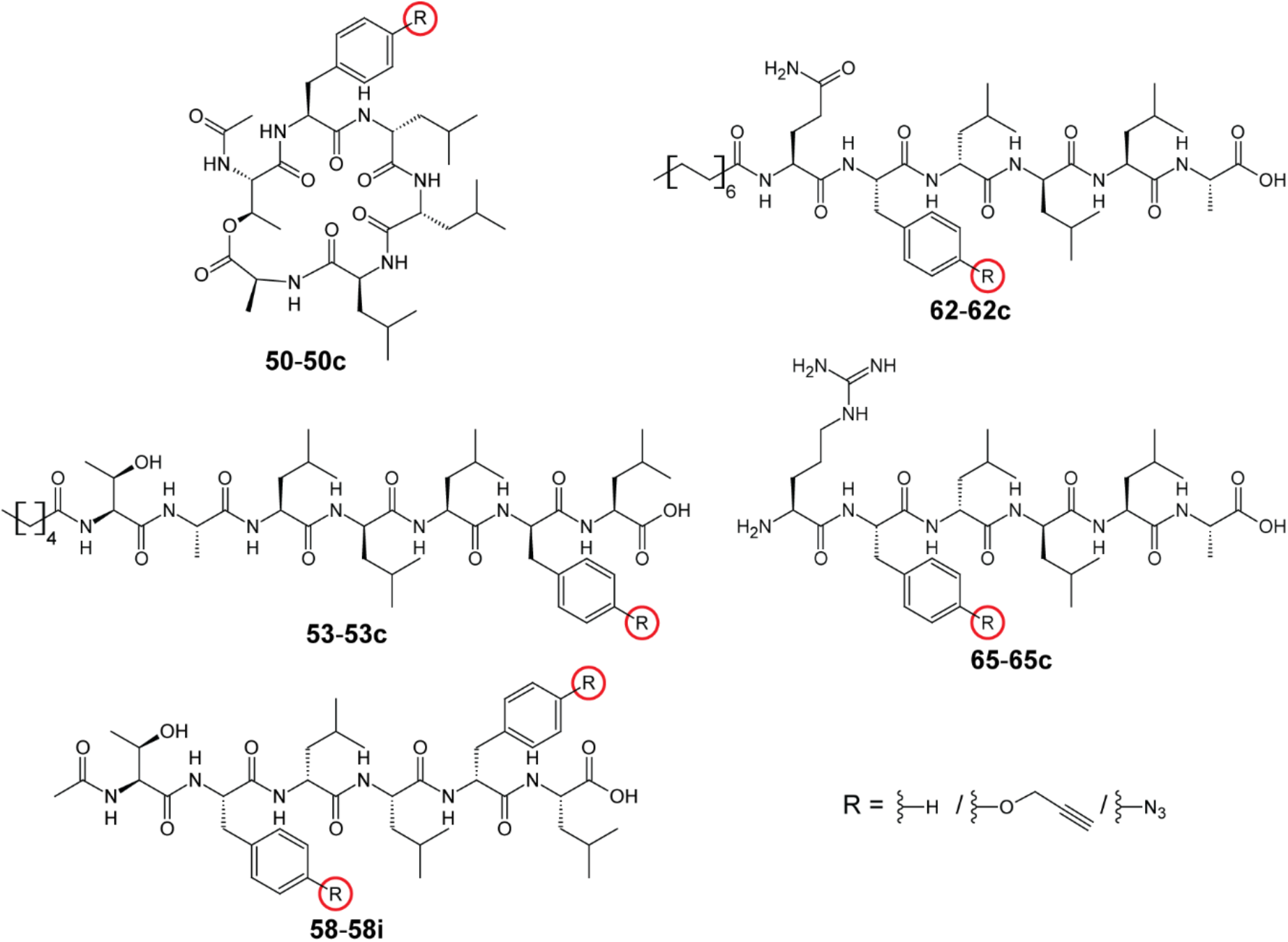

