## Supplementary Methods, Tables & Figures for "Site-Specific Introduction of Non-Canonical Amino Acids into natural and engineered Non-Ribosomal Peptides"

### Supplementary Information

#### Selection of ncAA

The selection of ncAA (Figure S1) was mainly based on substrates with reactive side-chain residues suitable for bioorthogonal click chemistry[1] with the focus on azides (**6**, **8-12**) and alkynes (**3-4**, **13**) for copper-catalyzed azide-alkyne cycloaddition (CuAAC)[2,3]. Here, propargylglycine (Pra, **4**) is the only naturally occurring amino acid[4–6]. The terminal alkynes can additionally be used in Sonogashira cross couplings[7,8] with $sp^2$-carbon-bearing halides like *p*-bromo-phenylalanine (*p*BrF, **16**). This reaction is also possible copper-free in water and *in vivo*[9].

The diene 3-(2-furyl)-alanine (2-Fua, **7**) can be used together with the dienophile *O*-allyl-tyrosine (*O*AlY, **18**) in Diels-Alder reactions[10,11]. Its analogue, 3-(3-furyl)-alanine (Fua), occurs in natural NRP[6] and is the key residue in the hepatotoxic peptide rhizonin[12,13].

3,4-dihydroxy-phenylalanine (DOPA, **15**), also a natural occurring amino acid, can be enzymatically functionalized with tyrosinase to a 1,2-benzoquinone, which can react with nucleophiles as Michael-acceptor[14,15]. Additionally, it can be *O*-alkylated using the mammalian catechol-*O*-methyl transferase using *S*-adenosyl methionine (SAM) derivatives[16].

5-chloro-tryptophan (5ClW, **2**) and *p*-amino-phenylalanine (PAPA, **14**) are the only used ncAA occurring in some *Xenorhabdus* or *Photorhabdus* strains. From the tested stains, only *P. laumondii* TTO1 has a functional PAPA-operon. In *X. szentirmaii* DSM 16338, there is the flavin-dependent halogenase *Xszen*FHal that can generate 5ClW (**2**) from free tryptophan[17] which is incorporated into a diketopiperazine[18]. The tested szentiamide (**21**) naturally harbors non-chlorinated tryptophan[19]. Although there are several halogenated NRP known[20], natural 5ClW-harbouring NRP are extremely rare but 5ClW occurs in ribosomally-produced post-translational modified peptides (RiPP) like the lantibiotic microbisporicin (NAI-107)[21].

The remaining ncAA (**1**, **17**) do not harbor reactive side-chains and only add structural complexity. 2-azetidine carboxylic acid (Aze, **1**) is a natural but rare proline analogue that is found in NRPS-derived alkaloids[22]. Homophenylalanine (HPhe, **17**) also naturally occurs in NRP[23–25] and is used as the central pharmacophore in angiotensin-converting enzyme inhibitors like ramipril[26].

#### Detailed analysis of ncAA incorporation in the selected NRPs

##### Szentiamide

The szentiamide synthetase SzeS from *Xenorhabdus szentirmaii* is a hexamodular NRPS (Figure S7A) producing the cyclic depsipeptide szentiamide (**21**)[19,27,28]. Figure 2A shows the structure of szentiamide (**21**) as well as its derivatives (**21b**-**21e**) generated by PDB supplementing the production medium with 5ClW (**2**), PAPA (**14**), and 2-Fua (**7**), respectively (Figure S8, Figure S9, Table S14). This renders three out of six A-domains promiscuous towards ncAA producing five derivatives with moderate incorporation rates (Figure 2A, Table S14). No incorporation was observed for **12**, **13**, **15-17** (Table S14).

**Xenoamicin**

The xenoamicin synthetase XabABC from *X. doucetiae* DSM 17909 consists of three separate proteins forming a tridecamodular NRPS (Figure S7B) producing the depsipeptides xenoamicin A-H[29]. Figure 2B shows the structure of xenoamicin C (**22**) as well as its derivatives (**22b**-**22f**) generated by PDB supplementing the production medium with Aha (**9**), HPra (**5**), and Aze (**1**), respectively (Figure S10-Figure S12, Table S15). For **9** and **5**, the A-domain responsible for incorporation could not be determined by the $MS^2$ fragmentation data (Figure S11) since the they got only incorporated in trace amounts. **1** was accepted very well by both proline specific A-domains (Figure S10-Figure S12, Table S15) and no incorporation was observed for **3**, **4**, **6**, and **8** (Table S15).

**Xeneprotide**

The xeneprotide synthetase XnpAB from *X. stockiae* KJ12.1 consists of two separate proteins forming a tetramodular NRPS (Figure S7C) producing the cyclic depsipeptides xeneprotide A-C[30]. Figure 2C shows the structure of xeneprotide A (**27**) as well as its derivatives **27b**-**27q** generated by PDB supplementing the production medium with *p*$N_3$F (**12**), *O*PrY (**13**), PAPA (**14**), *p*BrF (**16**), HPhe (**17**), and Aze (**1**), respectively (Figure S13-Figure S15, Table S16). For all mentioned ncAA (except **1**), an ion corresponding to a single and double incorporation could be detected (Figure S13-Figure S15). In case of double incorporations, both tryptophan residues were replaced and **1** was incorporated 12-fold better in **27q** compared to Pro in **27** (Figure 2C, Figure S13, Figure S15A, Table S16). Addition of ncAAs **2** and **7** did not lead to novel derivatives.

**TaxIllaid**

The taxIllaid synthetase TxlAB from *X. indica* consists of two separate proteins forming a heptamodular NRPS (Figure S7D) producing the cyclic depsipeptides taxIllaid A-E[31]. Figure 2D shows the structure of taxIllaid E (**28**) as well as its derivatives **28b**-**28d** generated by precursor-directed biosynthesis supplementing the production medium with *p*$N_3$F (**12**), PAPA (**14**), and 2-Fua (**7**), respectively (Figure S16, Figure S17, Table S17). Their $MS^2$ fragmentation spectra (Figure S17) indicate that the A3-domain, naturally accepting Phe, Leu and Tyr, is responsible for the incorporation of **12**, **14**, and **7**, respectively. Ten additional substrates (**3-6**, **8**, **9**, **13**, **15-17**) were tested but no incorporations were observed (Table S17).

**Fitayylide**

The fitayylide synthetase FitAB from *X. innexi* DSM 16336 consists of two proteins forming a hexamodular NRPS (Figure S7E) producing the cyclic depsipeptide fitayylide (**29**)[32], which is a close xeinamide homologue[33]. The structure of it as well as its derivatives **29b-29e** generated by precursor-directed biosynthesis supplementing the production medium with *O*PrY (**13**), PAPA

(**14**), and *p*BrF (**16**), respectively, can be seen in Figure 2E (Figure S18-Figure S21, Table S18). However, the relative incorporation rates for all substrates were poor. Four additional substrates (**7**, **12**, **15**, **17**) were tested but no incorporations were detected (Table S18). The $MS^2$ fragmentation spectra of **29b** and **29c** do not clearly indicate which tyrosine is replaced by **16** and **13**, respectively (Figure S19). To elucidate the promiscuous Tyr-specific A-domain responsible for incorporation of **16** and **13**, an additional experiment with the two synthetic NRPS (NRPS-1 and NRPS-2) was performed (Figure S20, Figure S21). Both encoded NRPS produce a fitayylide derivative with a different starting module (**30** and **31**). Additionally, in NRPS-2 a phenylalanine-specific module from a different NRPS of *X. mauleonii* DSM 17908 replaces the second tyrosine-specific module (Figure S20A). Heterologous expression of *NRPS-1* and *NRPS-2* in *E. coli* DH10B::*mtaA*[34] produced **30** and **31**, respectively. Supplementing **13** to the production medium, only for NRPS-1 an incorporation could be determined (**30b**, Figure S20 B, C). Thus, the responsible A-domain for the incorporation **13** is the A5-domain.

#### Xenortide

The xenortide synthetase XndAB from *X. nematophila* ATCC 19061 consists of two separate proteins forming a bimodular NRPS (Figure S7F) producing the highly modified dipeptides xenortide C-D[27,35–37]. The second NRPS (XndB) ends with a terminal C-domain ($C_{term}$) instead of a TE-domain. This $C_{term}$-domain releases the peptide by attaching phenylethylamine (PEA) or tryptamine (TRA) to the *C*-terminus of the dipeptide[27,37]. With this, the xenortides are structurally similar to rhabdopeptides[36,38,39]. Figure 2G shows the structure of xenortide A (**34**) as well as its derivative **34h** generated by precursor-directed biosynthesis (Figure S22) supplementing the production medium with 2-Fua (**7**). Noteworthy, for the incorporation of **7** in **34h** also the *N*-methylation took place (Figure S22C). The relative production of **32h** compared to **34** was 46.7% (Table S19). No incorporation by an A-domain was observed for **12-17**.

#### PAX

The PAX synthetase PaxABC from *X. doucetiae* DSM 17909 consists of three proteins and produces the highly cationic, cyclic lipopeptide PAX (peptide-antimicrobial-*Xenorhabdus*, **35**, Figure S23)[40–42]. PAX is widespread among the genus of *Xenorhabdus*[30] and shows antimicrobial activity against Gram-positive bacterium *Micrococcus luteus*, the fungus *Fusarium oxysporum*[40], and the amoeba *Acanthamoeba castellanii*[43]. Additionally, PAX has been shown to associate to the cell-wall of its producer, thereby protecting it against other antimicrobial peptides by repelling them[44]. Precursor-directed biosynthesis supplementing the production medium with **8**, **9**, **6**, **10**, and **11** (Figure 1B) did not show any derivatives.

##### Protegomycin

The protegomycin synthetase PrtAB from *X. doucetiae* DSM 17909 consists of two separate proteins forming a hexamodular NRPS producing the cyclic lipopeptide protegomycin (Figure S24A)[45]. Precursor-directed biosynthesis supplementing the production medium with *O*PrY (**13**), PAPA (**14**), and DOPA (**15**) caused the generation of the protegomycin derivatives **36b**-**36j** (Figure S24). The tryptophan-specific A6-domain is able to incorporate **13** and **14** resulting in **36b** and **36d** (**36f**), respectively (Figure S24, Figure S25). This can be confirmed by the absence of the tryptophan-specific fragment ion of 159.09 *m/z* in their respective $MS^2$ spectra (Figure S25). Additionally, there were exchanges of up to three tyrosines detected (**36c**, **36e**-**36j**, Figure S24) with good incorporation rates (Table S20). For these, however, no position could be assigned using $MS^2$ fragmentation spectra (Figure S25). Supplementation of **12** and **17** did not cause any derivatives (Table S20).

### Methods

#### Molecular Biology

##### Polymerase Chain Reaction

The components for polymerase chain reaction (PCR) were mixed in a final volume of 20 µl in quadruplicates, each three percent DMSO and processed according to Table S1 and Table S2. Success of PCR was monitored by agarose gel electrophoresis.

##### Agarose gel Electrophoresis

For gel electrophoresis, 1% Biozym LE Agarose (w/v) gels with MIDORI Green Advance (Nippon Genetics) in 1x TRIS-acetate-EDTA (TAE) buffer were used. The sample size was 2.5 µl with 1x Gel Loading Dye Purple (NEB). Electrophoresis was performed at 140 V for 20-30 min. Afterwards, the DNA was visualized using UV light. 1 kb Plus DNA Ladder (NEB) was used as marker.

##### PCR purification

For PCR clean-up, samples were subjected to preparative agarose gel electrophoresis (1% Biozym LE Agarose (w/v), 1x TAE buffer, MIDORI Green Direct 1:10 (Nippon Genetics), 45 min, 120 V). DNA was extracted using the Monarch® DNA Gel Extraction Kit (NEB) according to the manufacturer's protocol.

**Restriction digest**

Plasmids were digested in a total volume of 20 µl with 1x rCutSmart™ (NEB), for 1 h at 37 °C. pMS095 was digested using1 µl of I-CeuI and I-SceI. pCEP_Km was digested with NdeI and PstI-HF with subsequent enzyme inactivation at 80 °C for 20 min.

**HiFi assembly**

PCR fragments were mixed with linearized vectors in an appropriate molar ratio, equal volume of NEBuilder® 2xHiFi Assembly Master Mix (NEB) was added, and subsequently incubated for 1 h at 50 °C. Vector linearization was obtained using PCR or restriction digest. Correct assembly was verified by sequencing (Microsynth).

**Golden Gate assembly**

Golden Gate assembly was done with NEBridge® Golden Gate Assembly Kit (BsaI-HF®v2) (NEB) using conditions shown in

**Table S3**.

##### Plasmid preparation

For plasmid preparation, 4 ml of LB medium ($Kan^{50}/Cm^{34}$) was inoculated with the respective colony and incubated (37 °C, 180-200 rpm) overnight. Plasmids were isolated using the Monarch® Spin Plasmid Miniprep Kit (NEB) according to the manufacturer's protocol.

#### Microbiology

##### Preparation of chemical competent *E. coli* cells

To prepare chemical competent cells, LB substituted with 1 M $MgCl_2$ and 1 M $MgSO_4$ was inoculated (1:50) with an overnight culture of *E. coli* DH10B::*mtaA*[34] or *E. coli* ST18[46] and incubated (37 °C, 200 rpm) to an $OD_{600}$ = 0.6. Cells were then chilled on ice for 30 min and centrifuged for 8 min at 1,230 g and 4 °C. The supernatant was discarded and the cells were resuspended in 1/3 culture volume of cold RF-I solution. After 30 min of incubation at 4 °C, cells were centrifuged (1,230 g, 8 min, 4 °C) and resuspended in 1/20 culture volume of cold RF-II solution. After additional 30 min at 0-4 °C, cells were frozen in liquid nitrogen in 50 µl aliquots, and stored at -75 °C.

##### Transformation

Chemical competent *E. coli* DH10B::*mtaA*[34] or *E. coli* ST18[46] cells were thawed on ice, mixed with 0.5-2.0 µl plasmid, HiFi, or Golden Gate mix. The cells were incubated on ice for 10-15 min and subsequently heat shocked for 45 s at 42 °C. For recovery, cells were cooled down on ice for 1 min, diluted with 900 ml antibiotic-free LB medium, and incubated for 1 h at 37 °C and 750 rpm. After recovery, cells were plated on selective LB-agar plates (including 50 µg/ml 5-Aminolevulinic acid hydrochloride (5-ALA) for ST18 cells)[46–49].

##### Heterologous expression

For heterologous expression of recombinant NRPS, *E. coli* DH10B::*mtaA*[34] is used since it co-expresses the PPTase MtaA[50,51] necessary for the activation of T-domains. Production cultures were inoculated with preculture 1:100 and grown for 72 h at 22 °C and 180-220 rpm in XPP3 medium or XPP medium supplemented with respective antibiotics and 0.02% *L*-(+)-arabinose (w/v) for induction and with or without 4% XAD-16N adsorber resin (v/v) to capture hydrophobic compounds.

##### Promotor Exchange

All promotor exchanges were carried out in the respective Δ*hfq* strain[48,52] following the protocol as described in Bode *et al.*, **2023**[49]. The biosynthetic gene clusters (BGC) for xeneprotide production in *X. stockiae* KJ12.1 and for taxIlIaid production in X. indica DSM 17382 were activated using the plasmids pPH4 (pCEP_Kan_*xnpA*) and pWJ6 (pCEP_Kan_*txlA*), respectively (Table S8, Table S9).

**Identification of promiscuous Adenylation domains**

To identify promiscuous A-domains, precursor directed biosynthesis experiments with non-cognate building blocks were performed. For the screening, promotor exchange strains[47,49] of *X. doucetiae* DSM 17909, *X. indica* DSM 17382, *X. szentirmaii* DSM 26338, *P. laumondii* TTO1, *X. stockiae* KJ12.1, and *X. nematophila* ATCC 19061 with Δ*hfq* background[48] were used. The *fitAB* genes were heterologously expressed in *E. coli* DH10B::*mtaA*[34]. 4 ml XPP medium supplemented with 1 mM of a non-cognate building block (in DMSO or water with up to 0.5% HCl (v/v)), 0.2% *L*-(+)-arabinose (w/v), and respective antibiotics were inoculated with 40 µl of pre-culture. The cultures were grown for 72 h at 28 °C and 220 rpm in a 24-well deep-well plate. Extracts were measured by liquid chromatography (LC) high-resolution-mass spectrometry (HR-MS) to identify natural products in which one or more of the supplemented building blocks are incorporated. Identification of new derivatives was based on feature-based molecular network analysis[53] and $MS^2$ fragmentation spectra.

#### Mass Spectrometry

##### Sample Preparation

**Cell cultures *without* XAD-16N adsorber resin**

Cultures were mixed with one volume equivalent of acetonitrile (ACN) and incubated for 20-30 min at 1,200-1,800 rpm at room temperature. Samples were subsequently centrifuged for 30 min at 13,000 g. The supernatant was diluted 1:5 in ACN and used for LC-(HR)-MS.

**Cell cultures *with* XAD-16N adsorber resin**

Liquid cultures were decanted and the remaining XAD-16N adsorber resin was taken up in the culture volume of methanol (MeOH) and incubated for 20-30 min at 160-180 rpm at room temperature. Samples were subsequently centrifuged for 30 min at 13,000 g. The supernatant was diluted 1:10 in MeOH and used for LC-(HR)-MS.

**PAX extraction (adapted from Vo *et al.*, 2021[44])**

1 ml cell culture of *X. doucetiae* DSM 17909::Δ*hfq*_pCEP_Kan_*paxA* grown 72 h at 28 °C and 220 rpm in XPPM with 0.2% *L*-(+)-arabinose was transferred into a 2 ml glass vail and centrifuged (5,000 g, 5 min, room temperature). The supernatant was removed and the cell pellet was sonicated for 15 min in a water bath. Afterwards, the pellet was freeze-dried and resuspended in 1 ml chloroform with the addition of 200 µl of a 1% formic acid solution (v/v). The sample was sonicated for 15 min in a water bath. For phase separation, the glass vail was centrifuged (5,000 g, 5 min, room temperature) and the aqueous phase was transferred to a fresh tube. Remaining debris and chloroform were precipitated by centrifugation (3 min, 10,000 g, room temperature). The aqueous supernatant was transferred into a fresh tube and freeze-dried. The pellet was resuspended in 200 µl of a 1% formic acid solution (v/v), centrifuged (20,000 g, 20 min, room temperature) and used for LC-HR-MS.

##### Mass Spectrometry Methods

**amaZon ion trap**

Extracts were measured by ultra-high pressure liquid chromatography mass spectrometry (UPLC MS). Separation by liquid chromatography was performed with an Agilent 1290 Infinity II UPLC system using a $C_{18}$-column (ACQUITY UPLCTM BEH, 130 Å, 2.1 mm x 100 mm, Waters) at 40 °C. 5 µl sample were injected and separated over a period of 20 min using a ACN gradient (0-2 min 5%, 2-14 min 5-95%, 14-15 min 95-100%, 15-18 min 100%, 18-20 min 5%) with 0.1% (v/v) formic acid at a flow rate of 0.4 ml/min. Data was acquired between 1-15 min. Mass spectrometric analysis was performed on a ESI (electrospray ionization) ion trap mass spectrometer (amaZon speed or amZon speed ETD, Bruker Daltonics). ESI-MS spectra were recorded in positive ion mode over a mass range of 100-1,200 *m/z* or 200-1,800 *m/z*, respectively. These spectra were analyzed using DataAnalysis 6.1 (Bruker Daltonics).

**High-resolution mass spectrometry**

Extracts were measured by UPLC MS. Separation by liquid chromatography was performed on a Bruker Elute UPLC system using a $C_{18}$-column (ACQUITY UPLCTM BEH, 130 Å, 2.1 mm x 100 mm, Waters) at 40 °C. 2 µl sample were injected and separated over a period of 19 min using a ACN gradient (0-2 min 5%, 2-14 min 5-95%, 14.0-14.1 min 95-100%, 14.1-16 min 100%, 16-19 min 5%) with 0.1% (v/v) formic acid at a flow rate of 0.4 ml/min. Data was acquired between 0-14 min. For internal calibration, a 10 mM sodium formate solution ($H_2O$:isopropanole, 1:1) was injected in each run with a flow rate of 0.4 ml/min between 16.0-16.3 min. Mass spectrometric analysis was performed on a VIP-HESI (vacuum insulated probe heated-electrospray ionization) quadrupole time-of-flight (qTOF) mass spectrometer (timsTOF fleX MALDI-2, Bruker Daltonics). ESI-MS spectra were recorded in positive ion mode over a mass range of 100-2,000 *m/z*. These spectra were analyzed using DataAnalysis 6.1 (Bruker Daltonics), GNPS[53], MZmine 3.9.0[54], and Cytoscape 3.9.1[55].

**Trapped ion mobility (tims) spectrometry**

Tims data was acquired using the timsTOF fleX MALDI-2 (Bruker Daltonics) mass spectrometer. For sample separation, the same UPLC system and gradient were used as stated above. Injection volume was 10 µl. The system was additionally calibrated manually using 10 mM sodium formate and ESI-L Low Concentration Tuning Mix (Agilent). Detection was performed in "Ultra Mode" measuring inversed reduced ion mobility ($1/K_0$) between 1.10-1.30 $Vs/cm^2$ with a ramp time of 589.7 ms and PASEF (parallel accumulation serial fragmentation). These spectra were analyzed using DataAnalysis 6.1 (Bruker Daltonics).

<u>Feature-based Molecular Network</u>

Spectrometry raw files from the timsTOF fleX MALDI-2 were converted to .mzXML files using MSConvert 3.0.24100-098a63b (Proteowizard). Those files were processed as following with MZmine 3.9.0[54]. Unless stated otherwise, the default settings were used in all steps. For an initial network, "mass detection" thresholds for MS and $MS^2$ were set to $10^3$ and $10^1$, respectively. For later analysis, the thresholds were increased. Feature detection was performed using "ADAP Chromatogram Builder" with a "min group size in # of scans" of five and "min highest intensity" of $10^1$ (for initial analysis). Generated chromatograms were deconvoluted using "local minimum resolver" with "min # of data points" of >5. Isotopes were grouped using the "$^{13}C$ isotope filter". Next, all generated feature lists were aligned using the "join aligner tool". "Weight for *m/z*" and "weight for RT" were set to one and the "retention time tolerance" was >3 min. The aligned feature list was filtered for “features with $MS^2$ scan” using the “feature list row filter”. Additionally, “gap filtering” was performed with the “same RT and *m/z* range gap filter” tool. A non-induced control measurement was used for “feature list blank subtraction”. The feature list generated in this way was exported in a format suitable for GNPS[53]. All files were uploaded to GNPS[53] using FileZilla® 3.66.5. Feature-based molecular networks were calculated using GNPS[53] with following settings: Mass tolerance of 0.02 Da, “min pairs cos” of 0.5 and “minimum matched fragment ions” of three. The obtained network was visualized using Cytoscape 3.9.1[55].

#### Synthesis

##### Inter-molecular Diels-Alder reaction for peptide derivatization

*X. doucetiae* DSM 17909::*Δhfq*_pCEP_Kan_*gspS*[48] was cultivated in 50 ml XPP3 medium for 4 h, 180 rpm, and 28 °C. After that, the production medium was supplemented 2 mM 2-Fua (**7**), 0.2% *L*-(+)-arabinose (w/v), and 4% (v/v) XAD-16N adsorber resin and incubated for 70 h at 28 °C and 180 rpm. After production, cells and medium were discarded and the resin was incubated with 50 ml methanol for 1 h at room temperature and 180 rpm. Methanol and resin were separated and the liquid fraction was centrifuged (13,000 g, 30 min) to precipitate particles. Subsequently, the methanol was evaporated using a V10-Touch system (Biotage) in 10 ml aliquots. Diels-Alder reaction was carried out using *N*-phenylmaleimide (**67**) under the conditions described in Ehinger *et al.*, **2024**[11]. Progress of the reaction was monitored by UPLC-MS (amaZon speed, Bruker Daltonics) operating in positive ion mode (Figure 5A, Figure S52A).

##### Inter-molecular CuAAC for peptide derivatization

*NRPS-14* was heterologously expressed by *E. coli* DH10B::*mtaA*[34] in 50 ml XPP3 medium for 2 h, 180 rpm, and 37 °C. After that, the production medium was supplemented 2 mM *O*PrY (**13**), 0.02% *L*-(+)-arabinose (w/v), and 4% (v/v) XAD-16N adsorber resin and incubated for 70 h at 22 °C and 180 rpm. After production, cells and medium were discarded and the resin was incubated with 50 ml methanol for 1 h at room temperature and 180 rpm. Methanol and resin were separated and the liquid fraction was centrifuged (13,000 g, 30 min) to precipitate particles. Subsequently, the methanol was evaporated using a V10-Touch system (Biotage) in 10 ml aliquots. The extract as well as 40 µmol 2-azido-2-desoxy-*D*-glucopyranose ($N_3$Gluc) were dissolved in 1 ml dry DMSO. Under argon atmosphere, 120 µmol DBU and 20 µmol Cu(I)Br were added subsequently. The reaction mix was stirred for 6 h at room temperature under argon atmosphere. Progress of the reaction was monitored by UPLC-MS (amaZon speed, Bruker Daltonics) operating in positive ion mode (Figure 5B, Figure S52B).

**Solid phase peptide synthesis (SPPS)**

For SPPS, the microwave-assisted peptide synthesizer Liberty Prime 2.0 (CEM) was used with iterative Fmoc standard strategy[56] utilizing DIC (*N*,*N'*-diisopropylcarbodiimide)/Oxyma/DIPEA (*N*,*N*-diisopropylethylamine) as coupling agents. SPPS was used to synthesize the peptide $C_2$-Thr(*L*)-*O*PrY(*L*)-Leu(*D*)-Leu(*L*)-*p*$N_3$F(*D*)-Leu(*L*) (**58h**). It was synthesized on a Fmoc-*L*-Leu pre-loaded Wang resin (50 µmol) in automated iterative cycles of coupling (90 °C, 5 min), deprotection (50 °C, 10 min), and washing. Amino acid deprotection was done with 25% (v/v) pyrrolidine in DMF (*N*,*N*-dimethylformamide). For final deprotection and cleavage (1 h, room temperature), 5 ml of a solution of trifuroacetic acid (TFA), $H_2O$, and triisopropylsilane (TIS) (95:2.5:2.5, v/v) was used.

**Intra-molecular CuAAC for peptide cyclisation**

0.22 mg (0.25 µmol, 1.0 equiv) of the synthetic peptide $C_2$-Thr(*L*)-*O*PrY(*L*)-Leu(*D*)-Leu(*L*)-*p*$N_3$F(*D*)-Leu(*L*) (**58h**) was dissolved in DMSO/$H_2O$ (4:1) in a final volume of 1 ml. Aqueous Cu(II)$SO_4$ solution (10 mM, 12.5 µL, 0.125 µmol, 0.5 equiv) was added to the peptide solution, and the mixture was briefly stirred. Subsequently, freshly prepared aqueous sodium ascorbate solution (10 mM, 50 µL, 0.50 µmol, 2.0 equiv) was added to initiate the reaction. The reaction mixture was stirred at room temperature for 16 hours. Progress of the reaction was monitored by UPLC-MS (amaZon speed, Bruker Daltonics) operating in positive ion mode (Figure S53).

### Tables

#### Methods

**Table S1.** Two-step PCR program for generation of homologues overhangs.

| Components | Steps | Temperature [°C] | Time [s] | |
|---|---|---|---|---|
| ○ forward Primer (4 pmol) | 1. | | | |
| ○ reverse Primer (4 pmol) | Initial denaturation | 98 | 180 | |
| ○ dNTPs (10 mM) | Denaturation | 98 | 10 | |
| ○ 1x Q5® Reaction Buffer / HF Buffer | Annealing | $T_m$ - 2 | 15 | x5 |
| ○ Q5® HF / Phusion® Hot Start Flex DNA Pol (2 U/µl) | Elongation | 72 | 30/1 kb | |
| ○ (g)DNA template (0.1 µl) | 2. | | | |
| ○ DMSO (3%) | Denaturation | 98 | 8 | |
| ○ $ddH_2O$ | Annealing | 72 | 12 | x25 |
| | Elongation | 72 | 30/1 kb | |
| | Final Elongation | 72 | 300 | |

**Table S2.** One-step PCR program.

| Components | Steps | Temperature [°C] | Time [s] | |
|---|---|---|---|---|
| ○ Forward primer (4 pmol) | Initial denaturation | 98 | 30 | |
| ○ Reverse primer (4 pmol) | Denaturation | 98 | 10 | |
| ○ dNTPs (10 mM) | Annealing | $T_m$ - 2 | 15 | x30 |
| ○ 1x Q5® Reaction Buffer / HF Buffer | Elongation | 72 | 30/1 kb | |
| ○ Q5® HF DN Q5® HF / Phusion® Hot Start Flex DNA Pol (2 U/µl) | Final elongation | 72 | 300 | |
| ○ (g)DNA template (0.1 µl) | | | | |
| ○ DMSO (3%) | | | | |
| ○ $ddH_2O$ | | | | |

**Table S3.** Golden Gate assembly protocol. Based on Podolski *et al*., **2025**[57,58].

| Components (10 µl) | Steps | Temperature [°C] | Time [s] | |
|---|---|---|---|---|
| ○ Acceptor Plasmid (38 ng) | Digestion | 37 | 60 | x30 |
| ○ donor Plasmid (38 ng) | Ligation | 16 | 60 | |
| ○ 1x T4 DNA ligase Buffer | Heat inactivation | 60 | 300 | |
| ○ NEB Golden Gate Assembly Mix (0.5 µl) | | | | |
| ○ $ddH_2O$ | | | | |

#### Materials

**Table S4.** List of all chemicals used.

| Chemical | Supplier |
|---|---|
| acetic acid ≥99.0% | Merck KGaA, Darmstadt, Germany |
| acetonitrile ≥99.9% (ACN) | Sigma-Aldrich, St. Louis, USA |
| *L*-alanine (A) ≥98.0% | Sigma-Aldrich, St. Louis, USA |
| β-alanine (β-Ala) ≥99.0% | Sigma-Aldrich, St. Louis, USA |
| Amberlite® XAD16N 20-60 mesh | Sigma-Aldrich, St. Louis, USA |
| (2*S*,3*S*)-2-amino-3-azidobutanoic acid (Abu) ≥99.0% | Iris Biotech GmbH, Marktredwitz, Germany |
| *p*-aminobenzoic acid ≥99.0% | Sigma-Aldrich, St. Louis, USA |
| 5-Aminolevulinic acid hydrochloride (5-ALA) ≥98.0% | Carl Roth GmbH & Co. KG, Karlsruhe, Germany |
| *p*-amino-*L*-phenylalanine*HCl (PAPA) ≥96.0% | Sigma-Aldrich, St. Louis, USA |
| ammonium heptamolybdate tetra hydrate ≥99.0%; $(NH_4)_6Mo_7O_{24}$*4 $H_2O$ | Carl Roth GmbH & Co. KG, Karlsruhe, Germany |
| ammonium sulfate ≥99.5%; $(NH_4)_2SO_4$ | Carl Roth GmbH & Co. KG, Karlsruhe, Germany |
| *L*-(+)-arabinose (ara) ≥99.0% | Carl Roth GmbH & Co. KG, Karlsruhe, Germany |
| *L*-arginine (R) ≥98.5% | Sigma-Aldrich, St. Louis, USA |
| *L*-(+)-ascorbic acid ≥99.0% | Carl Roth GmbH & Co. KG, Karlsruhe, Germany |
| *L*-asparagine (N) ≥98.0% | Sigma-Aldrich, St. Louis, USA |
| *L*-aspartate (D) ≥98.0% | Sigma-Aldrich, St. Louis, USA |
| *L*-2-azetidine carboxylic acid (Aze) ≥99.0% | Sigma-Aldrich, St. Louis, USA |
| 3-azido-*L*-alanine*HCl hydrate (Aza) ≥98.0% | Iris Biotech GmbH, Marktredwitz, Germany |
| 2-azido-2-desoxy-*D*-glucopyranose ($N_3$Gluc) ≥97.0% | Sigma-Aldrich, St. Louis, USA |
| 4-azido-*L*-homoalanine*HCl (Aha) ≥98.0% | Iris Biotech GmbH, Marktredwitz, Germany |
| $N_\varepsilon$-azido-*L*-lysine ($N_3$Lys) ≥99.0% | Iris Biotech GmbH, Marktredwitz, Germany |
| $N_\delta$-azido-*L*-ornithine*HCl ($N_3$Orn) ≥98.0% | Iris Biotech GmbH, Marktredwitz, Germany |
| *p*-azido-*L*-phenylalanine (*p*$N_3$F) ≥98.0% | Bachem AG, Bubendorf, Switzerland |

| Chemical | Supplier |
|---|---|
| Bacto$^{TM}$ Proteose Peptone No.3 | Thermo Fisher Scientific, Waltham, USA |
| *D*-(+)-biotin ≥98.5% | Carl Roth GmbH & Co. KG, Karlsruhe, Germany |
| Biozym LE Agarose | Biozym Scientific GmbH, Oldendorf, Germany |
| *p*-bromo-*DL*-phenylalanine (*p*BrF) | Bachem AG, Bubendorf, Switzerland |
| calcium (II) chloride dihydrate ≥99.0%; $CaCl_2$*2 $H_2O$ | Carl Roth GmbH & Co. KG, Karlsruhe, Germany |
| chloramphenicol (Cm) ≥98.5% | Carl Roth GmbH & Co. KG, Karlsruhe, Germany |
| 5-chloro-*DL*-tryptophan (5ClW) | Santa Cruz Biotechnology, Inc., Dallas, USA |
| cobalamin ≥96.0% | Carl Roth GmbH & Co. KG, Karlsruhe, Germany |
| copper (I) bromide ≥98.0%; Cu(I)Br | TCI Deutschland GmbH, Eschborn, Germany |
| copper (II) chloride dihydrate ≥99.0%; $CuCl_2$*2 $H_2O$ | Carl Roth GmbH & Co. KG, Karlsruhe, Germany |
| copper (II) sulfate ≥99.0%; $CuSO_4$ | Carl Roth GmbH & Co. KG, Karlsruhe, Germany |
| 10x rCutSmart® buffer | New England Biolabs, Ipswich, USA |
| *L*-cysteine (C) ≥98.0% | Sigma-Aldrich, St. Louis, USA |
| deoxy nucleoside triphosphates (dNTPs) | Carl Roth GmbH & Co. KG, Karlsruhe, Germany |
| 1,8-diazabicyclo[5.4.0]undec-7-ene (DBU) | Fisher Scientific GmbH, Schwerte, Germany |
| Difco$^{TM}$ agar | Becton, Dickinson and Company, Sparks, USA |
| 3,4-dihydroxy-*DL*-phenylalanine (DOPA) ≥98.0% | Fisher Scientific GmbH, Schwerte, Germany |
| *N*,*N*'-diisopropylcarbodiimide (DIC) | Iris Biotech GmbH, Marktredwitz, Germany |
| *N*,*N*-diisopropylethylamine (DIPEA) | Sigma-Aldrich, St. Louis, US |
| *N*,*N*-dimethylformamide (DMF) ≥99.8 % | Thermo Fisher Scientific, Waltham, USA |
| dimethyl sulfoxide (DMSO) ≥99.0% | Carl Roth GmbH & Co. KG, Karlsruhe, Germany |
| dipotassium (I) hydrogen phosphate ≥99.0%; $K_2HPO_4$ | Carl Roth GmbH & Co. KG, Karlsruhe, Germany |
| 1 kb Plus DNA Ladder | New England Biolabs, Ipswich, USA |
| ethanol ≥99.5% | Carl Roth GmbH & Co. KG, Karlsruhe, Germany |
| ethanol 70%, denatured | Carl Roth GmbH & Co. KG, Karlsruhe, Germany |
| ethylenediaminetetraacetic acid, (EDTA) ≥99.0% | Carl Roth GmbH & Co. KG, Karlsruhe, Germany |
| $N_\alpha$-(9-fluorenylmethoxycarbonyl)-*O*-(*tert*-butyl)-*L*-threonine (Fmoc-*L*-Thr(*t*Bu)-OH) | GL Biochem Ltd., Shanghai, China |
| $N_\alpha$-(9-fluorenylmethoxycarbonyl)-*L*-leucine (Fmoc-*L*-Leu-OH) | GL Biochem Ltd., Shanghai, China |
| $N_\alpha$-(9-fluorenylmethoxycarbonyl)-*D*-leucine (Fmoc-*D*-Leu-OH) | GL Biochem Ltd., Shanghai, China |

| Chemical | Supplier |
|---|---|
| Fmoc-*L*-Leu-Wang resin; 100-200 mesh 0.5-1.3 mmol/g | Iris Biotech GmbH, Marktredwitz, Germany |
| $N_\alpha$-(9-fluorenylmethoxycarbonyl)-4-azido-*D*-phenylalanine (Fmoc-*D*-Phe(4-$N_3$)-OH) | Iris Biotech GmbH, Marktredwitz, Germany |
| $N_\alpha$-(9-fluorenylmethoxycarbonyl)-*O*-propargyl-*L*-tyrosine (Fmoc-*L*-Tyr(Propargyl)-OH) | Iris Biotech GmbH, Marktredwitz, Germany |
| folic acid ≥96.0% | Thermo Fisher Scientific, Waltham, USA |
| formic acid (FA) ≥99.0% | VWR International GmbH, Darmstadt, Germany |
| 3-(2-Furyl)-*L*-alanine (2-Fua) | Iris Biotech GmbH, Marktredwitz, Germany |
| Gel Loading Dye Purple (6x) | New England Biolabs, Ipswich, USA |
| GibcoTM Bacto$^{TM}$ tryptone | Life Technologies Corporation, Miami, USA |
| GibcoTM Bacto$^{TM}$ yeast extract, technical | Life Technologies Corporation, Miami, USA |
| *L*-glutamine (Q) ≥99.0% | Sigma-Aldrich, St. Louis, US |
| *L*-glutamate (E) ≥99.0% | Sigma-Aldrich, St. Louis, US |
| glycerol ≥98.0% | Carl Roth GmbH & Co. KG, Karlsruhe, Germany |
| glycine (G) ≥99.0% | Carl Roth GmbH & Co. KG, Karlsruhe, Germany |
| 5x HF Buffer | New England Biolabs, Ipswich, USA |
| *L*-histidine (H) ≥99.0% | Sigma-Aldrich, St. Louis, US |
| *L*-homopropargylglycine*HCl (HPra) | Sigma-Aldrich, St. Louis, US |
| *L*-homophenylalanine (HPhe) ≥98.0% | Fisher Scientific GmbH, Schwerte, Germany |
| hydrochloric acid 25%; HCl | Carl Roth GmbH & Co. KG, Karlsruhe, Germany |
| iron (III) chloride hexahydride; $FeCl_3$*6 $H_2O$ | Merck AG, Darmstadt, Germany |
| *L*-isoleucine (I) ≥98.0% | Sigma-Aldrich, St. Louis, US |
| kanamycin sulfate (Km) | Sigma-Aldrich, St. Louis, USA |
| *L*-leucine (L) ≥98.0% | Sigma-Aldrich, St. Louis, USA |
| *L*-lysine hydrochloride (K) ≥98.0% | Sigma-Aldrich, St. Louis, USA |
| magnesium (II) chloride ≥99.0%; $MgCl_2$ | Carl Roth GmbH & Co. KG, Karlsruhe, Germany |
| magnesium (II) sulfate hepta hydrate ≥99.0%; $MgSO_4$*7 $H_2O$ | Carl Roth GmbH & Co. KG, Karlsruhe, Germany |
| manganese (II) chloride tetra hydrate ≥99.0%; $MnCl_2$*4 $H_2O$ | Sigma-Aldrich, St. Louis, USA |
| methanol (MeOH) ≥99.9% | Fisher Scientific GmbH, Schwerte, Germany |
| *L*-methionine (M) ≥98.0% | Sigma-Aldrich, St. Louis, USA |
| MIDORI Green Advance | Nippon Genetics Europe GmbH, Düren, Germany |
| MIDORI Green Direct | Nippon Genetics Europe GmbH, Düren, Germany |
| 3-(*N*-morpholino)-propanesulfonic acid (MOPS) ≥99.5% | Carl Roth GmbH & Co. KG, Karlsruhe, Germany |
| nicotinic acid ≥98.0% | Carl Roth GmbH & Co. KG, Karlsruhe, Germany |

| Chemical | Supplier |
|---|---|
| Oxyma Pure ≥99.5% (ethyl cyano(hydroxyimino)acetate) | Sigma-Aldrich, St. Louis, USA |
| *D*-pantothenic acid hemi calcium salt ≥98.0% | Thermo Fisher Scientific, Waltham, USA |
| *L*-phenylalanine (F) ≥98.0% | Sigma-Aldrich, St. Louis, USA |
| potassium (I) acetate ≥99.0% | Carl Roth GmbH & Co. KG, Karlsruhe, Germany |
| potassium (I) dihydro phosphate ≥99.0%; $KH_2PO_4$ | Carl Roth GmbH & Co. KG, Karlsruhe, Germany |
| *DL*-propargylglycine (Pra) ≥98.0% | Sigma-Aldrich, St. Louis, USA |
| *O*-propargyl-*L*-serine*HCl (*O*PrS) | Iris Biotech GmbH, Marktredwitz, Germany |
| *O*-propargyl-*L*-tyrosine (*O*PrY) | Iris Biotech GmbH, Marktredwitz, Germany |
| *L*-proline (P) ≥99.0% | Sigma-Aldrich, St. Louis, USA |
| pyridoxine hydrochloride ≥99.0% | SERVA Electrophoresis GmbH, Heidelberg, Germany |
| pyrrolidine 99% | Sigma-Aldrich, St. Louis, USA |
| Q5® Reaction Buffer | New England Biolabs, Ipswich, USA |
| riboflavin ≥97.0% | Carl Roth GmbH & Co. KG, Karlsruhe, Germany |
| rubidium chloride ≥99.0%; RbCl | Sigma-Aldrich, St. Louis, USA |
| *L*-serine (S) ≥99.0% | Sigma-Aldrich, St. Louis, USA |
| sodium (I) chloride ≥99.5%; NaCl | Carl Roth GmbH & Co. KG, Karlsruhe, Germany |
| sodium (I) hydroxide ≥98.0%; NaOH | Carl Roth GmbH & Co. KG, Karlsruhe, Germany |
| sodium (I) pyruvate ≥99.0% | Carl Roth GmbH & Co. KG, Karlsruhe, Germany |
| sodium (I) tetraborate decahydrate; $Na_2B_4O_7$*10 $H_2O$ | Merck KGaA, Darmstadt, Germany |
| thiamine hydrochloride ≥99.0% | Sigma-Aldrich, St. Louis, USA |
| *L*-threonine (T) ≥99.5% | Sigma-Aldrich, St. Louis, USA |
| trifuroacetic acid (TFA) ≥99.9% | Carl Roth GmbH & Co. KG, Karlsruhe, Germany |
| triisopropylsilane (TIS) ≥98.0% | Sigma-Aldrich, St. Louis, USA |
| tris(hydroxymethyl)aminomethane (TRIS) ≥99.9% | Sigma-Aldrich, St. Louis, USA |
| trisodium (I) citrate dihydrate ≥99.0% | Carl Roth GmbH & Co. KG, Karlsruhe, Germany |
| *L*-tryptophan (W) ≥98.0% | Sigma-Aldrich, St. Louis, USA |
| *L*-tyrosine (Y) ≥98.0% | Sigma-Aldrich, St. Louis, USA |
| *L*-valine (V) ≥98.0% | Sigma-Aldrich, St. Louis, USA |
| zinc (II) chloride ≥97%; $ZnCl_2$ | Sigma-Aldrich, St. Louis, USA |

#### Kits and Enzymes

**Table S5.** Kits and enzymes used in this work.

| Kits and Enzymes | Supplier |
|---|---|
| Monarch® DNA Gel Extraction Kit | New England Biolabs, Ipswich, USA |
| Monarch® Spin Plasmid Miniprep Kit | |
| NEBridge® Golden Gate Assembly Kit (BsaI-HF®v2) | |
| NEBuilder® 2xHiFi Assembly Master Mix | |
| I-CeuI | |
| I-SceI | |
| NdeI | |
| PstI-HF | |
| Phusion® Hot Start Flex DNA Polymerase | |
| Q5® High Fidelity DNA Polymerase | |

#### Media, and Buffes

**Table S6.** Composition of used media.

| Media | Components |
|---|---|
| Lysogeny broth (LB) medium [agar]<br>pH 7.0 | 10.0 g/l Gibco™ Bacto™ tryptone |
| | 5.0 g/l Gibco™ Bacto™ yeast extract |
| | 5.0 g/l NaCl |
| | [1.5% Difco™ agar (w/v)] |
| *Xenorhabdus*/*Photorhabdus* production medium (XPPM)[48]<br>pH 7.0 | 10.0 g/l glycerol |
| | 2.0 g/l amino acid mix (5% (w/w) of each proteinogenic *L*-amino acid) |
| | 1.0 g/l sodium pyruvate |
| | 2.0% Salt A solution (v/v) |
| | 2.0% Salt B solution (v/v) |
| | 0.2% vitamin solution (v/v) |
| | 0.1% trace element solution (v/v) |
| XPP3 medium[59]<br>pH 7.0 | 10.0 g/l glycerol |
| | 10.0 g/l Bacto™ Proteose Peptone No.3 |
| | 1.0 g/l sodium pyruvate (optional) |
| | 2.0% Salt A solution (v/v) |
| | 2.0% Salt B solution (v/v) |
| | 0.2% vitamin solution (v/v) |
| | 0.1% trace element solution (v/v) |

**Table S7.** Composition of buffers and solutions used for gel electrophoresis, media, and competent cells.

| Buffers and Solutions | Components |
|---|---|
| 50x TRIS-acetate-EDTA (TAE) buffer pH 8.6 | 242 g/l TRIS |
| | 18.6 g/l EDTA |
| | 5.7% acetic acid (v/v) |
| Salt A solution | 350 g/l $K_2HPO_4$ |
| | 100 g/l $KH_2PO_4$ |
| Salt B solution | 50.0 g/l $(NH_4)_2SO_4$ |
| | 29.4 g/l trisodium citrate*2 $H_2O$ |
| | 5.0 g/l $MgSO_4$ |
| trace element solution | 0.2 g/l $FeCl_3$*6 $H_2O$ |
| | 40.0 mg/l $ZnCl_2$ |
| | 10.0 mg/l $CuCl_2$*2 $H_2O$ |
| | 10.0 mg/l $MnCl_2$*4 $H_2O$ |
| | 10.0 mg/l $Na_2B_4O_7$*10 $H_2O$ |
| | 10.0 mg/l $(NH_4)_6Mo_7O_{24}$*4 $H_2O$ |
| vitamin solution | 12.0 g/l pyridoxine hydrochloride |
| | 2.3 g/l nicotinic acid |
| | 1.2 g/l *D*-pantothenic acid hemi calcium salt |
| | 1.0 g/l thiamine hydrochloride |
| | 0.2 g/l *p*-aminobenzoic acid |
| | 0.2 g/l riboflavin |
| | 20.0 mg/l cobalamin |
| | 10.0 mg/l folic acid |
| | 6.0 mg/l *D*-(+)-biotin |
| RF-I solution pH 5.8 | 150 g/l glycerol |
| | 12.0 g/l RbCl |
| | 9.9 g/l $MnCl_2$*4$H_2O$ |
| | 2.9 g/l potassium acetate |
| | 1.5 g/l $CaCl_2$*2$H_2O$ |
| RF-II solution pH 6.8 | 150 g/l glycerol |
| | 11.0 g/l $CaCl_2$*2$H_2O$ |
| | 2.1 g/l MOPS |
| | 1.2 g/l RbCl |

#### Plasmids

**Table S8.** Overview of used plasmid. GG: Golden Gate.

| Plasmids | Genotype and Description | Source |
|---|---|---|
| pACYC_SEVA | ori p15A, OriT, $cm^R$, *araE*, *araC*-$P_{BAD}$ | [60] |
| pAP37 | ori p15A, $cm^R$, *araC*-$P_{BAD}$, [XINDV2_04300]_C1-T1/2 - BsaI site - [PLUMV2_16690]_T4/2-TE; Golden Gate (GG) acceptor for A/B overhangs. | [58] |
| pAP37_AP41 | ori p15A, $cm^R$, *araC*-$P_{BAD}$, [XINDV2_04300]_C1-T1/2 - [XSZEV2_13025]_T4/2-T5/2 - [PLUMV2_16690]_T4/2-TE | |
| pAP39 | ori pUC, $gent^R$, [XSZEV2_13025]_T4/2-T5/2; GG donor for A/B overhangs. | |
| pAP41 | ori pUC, $gent^R$, [XMIRV2_10175]_T1/2-T2/2; GG donor with A/B overhangs. | |
| pCEP_Km | ori R6K, oriT, $kan^R$, *traI, araC*-$P_{BAD}$ | [47] |
| pCK_0431 | ori p15A, $cm^R$, *araC*-$P_{BAD}$, *tacI* (pACYC empty) | [59] |
| pCK_0433 | ori ColA, $kan^R$, *araC*-$P_{BAD}$, *tacI* (pCOLA empty) | |
| pCK_0447 | ori p15A and OriT, $cm^R$, *araC*-$P_{BAD}$, *tacI* (pACYC empty) | this work |
| pCK_0449 | ori ColA and OriT, $kan^R$, *araC*-$P_{BAD}$, *tacI* (pCOLA empty) | |
| pCK_0706 | ori p15A, $cm^R$, *araC*-$P_{BAD}$, [XMAUV2_08560] | |
| pCK_0683 | ori p15A, $cm^R$, *araC*-$P_{BAD}$, [XINV2_12405] - [XINV2_12410] | [32] |
| pCK_1099-c* | ori ColA and oriT, $kan^R$, *araC*-$P_{BAD}$, [XINNV2_12410] | this work |
| pCK_1134-a | ori p15A, $cm^R$, *araC*-$P_{BAD}$, [XBOVV2_06145]_C$_{start}$-A1 - [XINNV2_12405]_T1-E4-DD | |
| pMS091* | ori p15A, $cm^R$, *araC*-$P_{BAD}$, [XINDV2_04300]_C1-T1/2 - BsaI site - [XINDV2_06865]_T4/2-E5 | |
| pMS091 | ori p15A, $cm^R$, *araC*-$P_{BAD}$, [XINDV2_04300]_C1-T1/2 - BsaI site - [XINDV2_06865]_T4/2-E5_C6272T (silent point mutation to remove BsaI restriction site); GG acceptor for A/B overhangs. | |
| pMS094 | ori p15A, $cm^R$, *araC*-$P_{BAD}$, [XBUDV2_10055]_A1-T1/2 - BsaI site - [XINDV2_06865]_T4/2-E5_C6272T; GG acceptor for A/B overhangs. | |
| pMS095 | ori p15A, $cm^R$, *araC*-$P_{BAD}$, XINDV2_06870]_C5-T5/2 - BsaI site - [PLUMV2_16690]_T4/2-TE; GG acceptor for A/B overhangs. | |
| pMS099 | ori pUC, $gent^R$, [XDOUV2_17010]_T2/2-T3/2; GG donor with A/B overhangs. | |
| pMS100 | ori pUC, $gent^R$, [XINDV2_06865]_T2/2-T3/2; GG donor with A/B overhangs. | |
| pMS101 | ori pUC, $gent^R$, [XDOUV2_09055]_T5/2-T6/2; GG donor with A/B overhangs. | |

| Plasmids | Genotype and Description | Source |
|---|---|---|
| pMS102 | ori pUC, *gent*$^R$, [XEKJV2_09200]_T1/2-T2/2; GG donor with A/B overhangs. | |
| pMS103 | ori pUC, *gent*$^R$, [XEKJV2_09200]_T2/2-T3/2; GG donor with A/B overhangs. | |
| pMS104 | ori pUC, *gent*$^R$, [XDOUV2_10795]_T2/2-T3/2; GG donor with A/B overhangs. | |
| pMS105 | ori pUC, *gent*$^R$, [XSZEV2_13025]_T5/2-T6/2; GG donor with A/B overhangs. | |
| pMS107 | ori pUC, *gent*$^R$, [XINNV2_12410]_T4/2-T5/2; GG donor with A/B overhangs. | |
| pMS108 | ori pUC, *gent*$^R$, [XDOUV2_10775]_T3/2-T4/2; GG donor with A/B overhangs. | |
| pMS109 | ori ColA, oriT, *kan*$^R$, *araC-P*$_{BAD}$, [XINDV2_06870]_C5-T5/2 - BsaI site - [PLUMV2_16690]_T4/2-TE; GG acceptor for A/B overhangs. | |
| pMS113 | ori p15A, OriT, *cm*$^R$, *araC-P*$_{BAD}$, [XINDV2_06865]_C1-E5 | |
| pMS114 | ori ColA, OriT, *kan*$^R$, *araC-P*$_{BAD}$, [XINDV2_06870]_C5-TE | |
| pMS121 | ori p15A, OriT, *cm*$^R$, *araE*, *araC-P*$_{BAD}$, [XMAUV2_12890]_C1-T1/2 - BsaI site - [XINDV2_06865]_T4/2-E5_C6272T; GG acceptor for A/B overhangs. | this work |
| pAP37_MS099 | ori p15A, *cm*$^R$, *araC-P*$_{BAD}$, [XINDV2_04300]_C1-T1/2 - [XDOUV2_17010]_T2/2-T3/2 - [PLUMV2_16690]_T4/2-TE | |
| pAP37_MS100 | ori p15A, *cm*$^R$, *araC-P*$_{BAD}$, [XINDV2_04300]_C1-T1/2 - [XINDV2_06865]_T2/2-T3/2 - [PLUMV2_16690]_T4/2-TE | |
| pAP37_MS101 | ori p15A, *cm*$^R$, *araC-P*$_{BAD}$, [XINDV2_04300]_C1-T1/2 - [XDOUV2_09055]_T5/2-T6/2 - [PLUMV2_16690]_T4/2-TE | |
| pAP37_MS102 | ori p15A, *cm*$^R$, *araC-P*$_{BAD}$, [XINDV2_04300]_C1-T1/2 - [XEKJV2_09200]_T1/2-T2/2 - [PLUMV2_16690]_T4/2-TE | |
| pAP37_MS103 | ori p15A, *cm*$^R$, *araC-P*$_{BAD}$, [XINDV2_04300]_C1-T1/2 - [XEKJV2_09200]_T2/2-T3/2 - [PLUMV2_16690]_T4/2-TE | |
| pAP37_MS104 | ori p15A, *cm*$^R$, *araC-P*$_{BAD}$, [XINDV2_04300]_C1-T1/2 - [XDOUV2_10795]_T2/2-T3/2 - [PLUMV2_16690]_T4/2-TE | |
| pAP37_MS105 | ori p15A, *cm*$^R$, *araC-P*$_{BAD}$, XINDV2_04300]_C1-T1/2 - [XSZEV2_13025]_T5/2-T6/2 - [PLUMV2_16690]_T4/2-TE | |

| Plasmids | Genotype and Description | Source |
|---|---|---|
| pAP37_MS107 | ori p15A, $cm^R$, $araC\text{-}P_{BAD}$, [XINDV2_04300]_C1-T1/2 - [XINNV2_12410]_T4/2-T5/2 - [PLUMV2_16690]_T4/2-TE | this work |
| pAP37_MS108 | ori p15A, $cm^R$, $araC\text{-}P_{BAD}$, [XINDV2_04300]_C1-T1/2 - [XDOUV2_10775]_T3/2-T4/2 - [PLUMV2_16690]_T4/2-TE | |
| pMS091_MS099 | ori p15A, $cm^R$, $araC\text{-}P_{BAD}$, [XINDV2_04300]_C1-T1/2 - [XDOUV2_17010]_T2/2-T3/2 - [XINDV2_06865]_T4/2-E5_C6272T | |
| pMS094_MS099 | ori p15A, $cm^R$, $araC\text{-}P_{BAD}$, [XBUDV2_10055]_A1-T1/2 - [XDOUV2_17010]_T2/2-T3/2 - [XINDV2_06865]_T4/2-E5_C6272T | |
| pMS109_MS099 | ori ColA, $kan^R$, $araC\text{-}P_{BAD}$, [XINDV2_06870]_C5-T5/2 - [XDOUV2_17010]_T2/2-T3/2 - [PLUMV2_16690]_T4/2-TE | |
| pMS121_MS099 | ori p15A, OriT, $cm^R$, *araE*, $araC\text{-}P_{BAD}$, [XMAUV2_12890]_C1-T1/2 - [XDOUV2_17010]_T2/2-T3/2 - [XINDV2_06865]_T4/2-E5_C6272T | |
| pMT001 | ori pUC, $gent^R$, [XMAUV2_08560]_T2/2-T3/2; GG donor for C/O overhangs. | |
| pMT006 | ori ColA, $kan^R$, $araC\text{-}P_{BAD}$, [XINNV2_12410]_DD-T4/2 - [XMAUV2_08560]_T2/2-T3/2 - [XINNV2_12410]_T5/2-TE | |
| pMT010 | ori ColA, OriT, $kan^R$, $araC\text{-}P_{BAD}$, [XINNV2_12410]_DD-T4/2 - BsaI site - [XINNV2_12410]_T5/2-TE; GG acceptor for C/O overhangs. | |
| pMT012 | ori ColA, OriT, $kan^R$, $araC\text{-}P_{BAD}$, [XINNV2_12410]_DD-T4/2 - BsaI site - [XINNV2_12410]_T5/2-TE; GG acceptor for A/B overhangs. | |
| pMT012_MS099 | ori ColA, OriT, $kan^R$, $araC\text{-}P_{BAD}$, [XINNV2_12410]_DD-T4/2 - [XDOUV2_17010]_T2/2-T3/2 - [XINNV2_12410]_T4/2-TE | |
| pPH4 (pCEP_Kan_*xnpA*) | ori R6K, oriT, $kan^R$, *traI*, $araC\text{-}P_{BAD}$, [XEKJV2_09200]_712 bp for promotor exchange | |
| pSEVA681 | ori pUC, $gent^R$ | [61] |
| pWJ9 (pCEP_Kan_*txlA*) | ori R6K, oriT, $kan^R$, *traI*, $araC\text{-}P_{BAD}$, [XINDV2_06865] 541 bp; for promotor exchange | this work |

**Table S9.** Cloning of all Plasmids generated in this work. All plasmids cyclized using NEBuilder® HiFi Assembly (NEB) or NEBridge® Golden Gate Assembly (NEB). Primer sequences given in red were used to create homologues overhangs; in green for point mutations; in yellow for insertions. Primers produced by Eurofins Scientific, Sigma Aldrich, and Metabion.

| Plasmid | Primer | Sequence (5' → 3') | Template |
|---|---|---|---|
| pCK_0447 | ck1230 | GGAAAAGGACAACGAAGAGAATCAGCTGTCCCTCCTGTTCAG | pCK_0431 |
| | ck1231 | CGAGGCTGGCCGTATGCATATAGAAACAGAAGCCACTGGAGC | |
| | ck1228 | TATGCATACGGCCAGCCTCGCAGA | pSEVA681 |
| | ck1229 | TTCTCTTCGTTGTCCTTTTCCGCTGCATAAC | |
| pCK_0449 | ck1228 | TATGCATACGGCCAGCCTCGCAGA | pSEVA681 |
| | ck1229 | TTCTCTTCGTTGTCCTTTTCCGCTGCATAAC | |
| | ck1232 | GGAAAAGGACAACGAAGAGAAAAACGTCCTAGAAGATGCCAGG | pCK_0433 |
| | ck1233 | GCGAGGCTGGCCGTATGCATACTGCGTCTAGCATGCCTATTTG | |
| pCK_0706 | ck0537 | CAGCTATTTTCATTGAATTCCTCCTGTTAGCCCAAA | pCK_0431 |
| | ck0538 | GTGAGGTGATGATTAATTAACCTAGGCTGCTGCC | |
| | ck0539 | CTAGGTTAATTAATCATCACCTCACACCAACAACTAC | gDNA *X. mauleonii* DSM 17908 |
| | ck0540 | CAGGAGGAATTCAATGAAAATAGCTGAGAAAGCGTTGGA | |
| pCK_1099-c* | ck1007 | CATGGAATTCCTCCTGTTAGCCCAAAAAAAC | pCK_0449 |
| | ck1010 | TTAATTAACCTAGGCTGCTGCCACCG | |
| | ck1372 | TTTTGGGCTAACAGGAGGAATTCCATGTTGATGCTATTCAAAGAACTCAATGAAAGTC | gDNA *X. innexi* DSM 16336 |
| | ck1600 | GCAGCAGCCTAGGTTAATTAAGCCAATACCTTTTCCTGACTTTCTGA | |
| pCK_1134-a | ck1007 | CATGGAATTCCTCCTGTTAGCCCAAAAAAAC | pCK_0431 |
| | ck1010 | TTAATTAACCTAGGCTGCTGCCACCG | |
| | ck1565 | TTTTGGGCTAACAGGAGGAATTCCATGGATAACATTCTGGCCTCGCCA | gDNA *X. bovienii* SS-2004 |

| Plasmid | Primer | Sequence (5' → 3') | Template |
|---|---|---|---|
| pCK_1134-a continued | ck1566 | TGCGTTACGGGGGGCAACATA AACATAGCGGCTCTGTTTAAAA TCTGG | |
| | ck0618 | TATGTTGCCCCCCGTAACGCA | gDNA *X. innexi* DSM 16336 |
| | ck1371 | GGTGGCAGCAGCCTAGGTTAA TTAATTACACATCCAAAATAGT TTTCTGTTCTGAGTGG | |
| pMS091* | ms341 | ACCAACGAATGCAAGGTCTCA TGGTAATTCAATTATGGCCATC AAGCTG | gDNA *X. indica* DSM 17382 |
| | ms342 | GGTGGCAGCAGCCTAGGTTAA TTAATTACACATCCAAAATGGT TTTCTTTTC | |
| | ms339 | TTAATTAACCTAGGCTGCTGCC ACC | pAP37 |
| | ms340 | ACCATGAGACCTTGCATTCGTT GGT | |
| pMS091 | ms385 | ATTCAATTGGTTTCTAAACTCC GGCAGG | pMS091* |
| | ms388 | TCAGGTAGGATCCGCTAATCTT ATGG | |
| | ms386 | GCCGGAGTTTAGAAACCAATT GAATACTG | |
| | ms387 | CATAAGATTAGCGGATCCTAC CTGACG | |
| pMS094 | ms349 | TTTTTGGGCTAACAGGAGGAA TTCCATGAAAGATAACATTGCT ACAGTG | gDNA *X. budapestensis* DSM 16342 |
| | ms350 | CCTTGCATTCGTTGGTCTCGTC CTCCCAATGCAAAAAAGCTGT CAT | |
| | ms342 | GGTGGCAGCAGCCTAGGTTAA TTAATTACACATCCAAAATGGT TTTCTTTTC | pMS091 |
| | ms344 | GGAATTCCTCCTGTTAGCCCA AAAAAAC | |
| pMS095 | ms351 | TTTTTGGGCTAACAGGAGGAA TTCCATGTTGAAACTATTAGAA GAACTGAATAAAAATC | gDNA *X. indica* DSM 17382 |
| | ms352 | ACCTTGCATTCGTTGGTCTCGT CCTCCGATACGGAAGAAGTTA TCCTC | |
| | ms343 | AGGACGAGACCAACGAATGCA AG | pAP37 |
| | ms344 | GGAATTCCTCCTGTTAGCCCA AAAAAAC | |

| Plasmid | Primer | | Sequence (5' → 3') | Template |
|---|---|---|---|---|
| pMS099 | ms359 | | CACACAGGAAAGAAGGTCTCAAGGACATTCTTTGTTGGCGGTACGGATG | gDNA *X. doucetiae* DSM 17909 |
| | ms360 | | CAGTCACGACCTTTGGTCTCTACCACCCAAGGCGAAGAAACTGTCATG | |
| | AP119 | | TCCTTGAGACCTTCTTTCCTGTGTGAAATTGTTATCCGCT | pAP39 |
| | AP120 | | TGGTAGAGACCAAAGGTCGTGACTGGGAAAACCC | |
| pMS100 | ms361 | | CACACAGGAAAGAAGGTCTCAAGGAAACTCCCTGCTGGCGATTAAACTG | gDNA *X. indica* DSM 17382 |
| | ms362 | | CAGTCACGACCTTTGGTCTCTACCACCAATGCGGAAGAAATTGTCCTCAATG | |
| | AP119 | | TCCTTGAGACCTTCTTTCCTGTGTGAAATTGTTATCCGCT | pAP39 |
| | AP120 | | TGGTAGAGACCAAAGGTCGTGACTGGGAAAACCC | |
| pMS101 | nested1 | ms332 | GCGATTTGGTGCGCCGGTTACC | gDNA *X. doucetiae* DSM 17909 |
| | | ms333 | GGCGAGGAGTAAGGGAGTGAGC | |
| | ms363 | | CACACAGGAAAGAAGGTCTCAAGGAAACTCCCTGACCGCCATTAAGC | nested1 |
| | ms364 | | CAGTCACGACCTTTGGTCTCTACCACCAATGCGGAAGAAATTATCTTCAATACC | |
| | AP119 | | TCCTTGAGACCTTCTTTCCTGTGTGAAATTGTTATCCGCT | pAP39 |
| | AP120 | | TGGTAGAGACCAAAGGTCGTGACTGGGAAAACCC | |
| pMS102 | ms365 | | CACACAGGAAAGAAGGTCTCAAGGAAATTCTCTGACCGCCATTAAACTGATTGC | gDNA *X. stockiae* KJ12.1 |
| | ms366 | | CAGTCACGACCTTTGGTCTCTACCACCGAGACGGAAGAAATTATCGTCAATAC | |
| | AP119 | | TCCTTGAGACCTTCTTTCCTGTGTGAAATTGTTATCCGCT | pAP39 |
| | AP120 | | TGGTAGAGACCAAAGGTCGTGACTGGGAAAACCC | |

| Plasmid | Primer | Sequence (5' → 3') | Template |
|---|---|---|---|
| pMS103 | ms367 | CACACAGGAAAGAAGGTCTCAAGGAAATTCTCTGAGCGCCATTAAACTGATTGCC | gDNA *X. stockiae* KJ12.1 |
| | ms368 | CAGTCACGACCTTTGGTCTCTACCACCAATGCGGAAGAAATTATCGTGGATGC | |
| | AP119 | TCCTTGAGACCTTCTTTCCTGTGTGAAATTGTTATCCGCT | pAP39 |
| | AP120 | TGGTAGAGACCAAAGGTCGTGACTGGGAAAACCC | |
| pMS104 | ms369 | CACACAGGAAAGAAGGTCTCAAGGACACTCGCTGATGATTGTCAGCCTGATCG | gDNA *X. doucetiae* DSM 17909 |
| | ms370 | CAGTCACGACCTTTGGTCTCTACCACCAAGCTCAAAGAAGTGATCATGGCGGC | |
| | AP119 | TCCTTGAGACCTTCTTTCCTGTGTGAAATTGTTATCCGCT | pAP39 |
| | AP120 | TGGTAGAGACCAAAGGTCGTGACTGGGAAAACCC | |
| pMS105 | ms371 | CACACAGGAAAGAAGGTCTCAAGGACATTCGTTGCTGGCGGTACGG | gDNA *X. szentirmaii* DSM 16338 |
| | ms372 | CAGTCACGACCTTTGGTCTCTACCACCCAAATCAAAGAAGCTGTCATGGCG | |
| | AP119 | TCCTTGAGACCTTCTTTCCTGTGTGAAATTGTTATCCGCT | pAP39 |
| | AP120 | TGGTAGAGACCAAAGGTCGTGACTGGGAAAACCC | |
| pMS107 | ms375 | CACACAGGAAAGAAGGTCTCAAGGAGATTCAATTATCAGTATTCAGTTAGTTTCCAGAC | gDNA *X. innexi* DSM 16336 |
| | ms376 | CAGTCACGACCTTTGGTCTCTACCACCGATACGGAAGAAATTATCTTCAATACCG | |
| | AP119 | TCCTTGAGACCTTCTTTCCTGTGTGAAATTGTTATCCGCT | pAP39 |
| | AP120 | TGGTAGAGACCAAAGGTCGTGACTGGGAAAACCC | |
| pMS108 | nested2: ms379 | GCCGGATTCAACAGATGTGGCAGTTC | gDNA *X. doucetiae* DSM 17909 |
| | nested2: ms380 | CAACATCGCGGGAGTGATGGCC | |
| | ms377 | CACACAGGAAAGAAGGTCTCAAGGACACTCATTGCTGGCTGTCCAACTCAT | nested2 |

| Plasmid | Primer | | Sequence (5' → 3') | Template |
|---|---|---|---|---|
| pMS108 continued | ms378 | | CAGTCACGACCTTTGGTCTCTACCACCGAGTTCAAAGAAATGATCATGGCGAC | |
| | AP119 | | TCCTTGAGACCTTCTTTCCTGTGTGAAATTGTTATCCGCT | pAP39 |
| | AP120 | | TGGTAGAGACCAAAGGTCGTGACTGGGAAAACCC | |
| pMS109 | Restric. Digest | I-CeuI | - | pMS095 |
| | | I-SceI | - | |
| | ck1097 | | GAGGGGTTTTTGCTAACTATAACGGTC | pCK_0449 |
| | ck1098 | | TAGCCGTCAAGTTGTCATAATTACCCTG | |
| pMS113 | ms393 | | TTTTTGGGCTAACAGGAGGAATTCCATGAGAATACCTGAAAGTGCGTTG | gDNA *X. indica* DSM 17382 |
| | ms396 | | TAGAGCGTGGTAACACTCCTCCAGC | |
| | ms342 | | GGTGGCAGCAGCCTAGGTTAATTAATTACACATCCAAAATGGTTTTCTTTTC | |
| | ms395 | | GCTGGAGGAGTGTTACCACGCTCTA | |
| | ms339 | | TTAATTAACCTAGGCTGCTGCCACC | pCK_0447 |
| | ms344 | | GGAATTCCTCCTGTTAGCCCAAAAAAAC | |
| pMS114 | ms351 | | TTTTTGGGCTAACAGGAGGAATTCCATGTTGAAACTATTAGAAGAACTGAATAAAAATC | gDNA *X. indica* DSM 17382 |
| | ms398 | | GGCAAGCTCGATATGAGGGATGAC | |
| | ms394 | | GGTGGCAGCAGCCTAGGTTAATTAATTATTCAGTAAATATTGATACACCTTGCTTAAC | |
| | ms397 | | GTCATCCCTCATATCGAGCTTGCC | |
| | ms339 | | TTAATTAACCTAGGCTGCTGCCACC | pCK_0449 |
| | ms344 | | GGAATTCCTCCTGTTAGCCCAAAAAAAC | |
| pMS121 | ms443 | | TTTTTGGGCTAACAGGAGGAATTCCATGACAAAATCTGAATATTTAGTAAGTTC | gDNA *X. mauleonii* DSM 17908 |
| | ms444 | | ACCATGAGACCTTGCATTCGTTGGTCTCGTCCTCCAATGCGGAAGAAATTATCG | |

| Plasmid | Primer | Sequence (5' → 3') | Template |
|---|---|---|---|
| pMS121 continued | ms341 | ACCAACGAATGCAAGGTCTCATGGTAATTCAATTATGGCCATCAAGCTG | pMS091 |
| | ms438 | TCGCCAGGGTTTTCCCAGTCACGACTTACACATCCAAAATGGTTTTCTTTTCTG | |
| | ms344 | GGAATTCCTCCTGTTAGCCCAAAAAAC | pACYC_SEVA |
| | ms435 | GTCGTGACTGGGAAAACCCTG | |
| pAP37_MS099 | Golden Gate | - | pAP37 |
| | | - | pMS099 |
| pAP37_MS100 | Golden Gate | - | pAP37 |
| | | - | pMS100 |
| pAP37_MS101 | Golden Gate | - | pAP37 |
| | | - | pMS101 |
| pAP37_MS102 | Golden Gate | - | pAP37 |
| | | - | pMS102 |
| pAP37_MS103 | Golden Gate | - | pAP37 |
| | | - | pMS103 |
| pAP37_MS104 | Golden Gate | - | pAP37 |
| | | - | pMS104 |
| pAP37_MS105 | Golden Gate | - | pAP37 |
| | | - | pMS105 |
| pAP37_MS107 | Golden Gate | - | pAP37 |
| | | - | pMS107 |
| pAP37_MS108 | Golden Gate | - | pAP37 |
| | | - | pMS108 |
| pMS091_MS099 | Golden Gate | - | pMS091 |
| | | - | pMS099 |
| pMS094_MS099 | Golden Gate | - | pMS094 |
| | | - | pMS099 |
| pMS109_MS099 | Golden Gate | - | pMS109 |
| | | - | pMS099 |
| pMS121_MS099 | Golden Gate | - | pMS121 |
| | | - | pMS099 |
| pMT001 | MT002 | CACACAGGAAAGAAGGTCTCACGGCGATTCCATTATCAGCATC | pCK_0706 |
| | MT003 | CACGACCTTTGGTCTCTACCGCCGATGCGGAAGAAATTATCTTCAA | |
| | MT011 | GCCGTGAGACCTTCTTTCCTGTGTGAAATTGTTATCCGC | pAP39 |
| | MT012 | CGGTAGAGACCAAAGGTCGTGACTGGGAAAAC | |

| Plasmid | Primer | Sequence (5' → 3') | Template |
|---|---|---|---|
| pMT006 | Golden Gate | - | pMT010 |
| | | - | pMT001 |
| pMT010 | ck1007 | CATGGAATTCCTCCTGTTAGCCCAAAAAAAC | pCK_0449 |
| | ck1010 | TTAATTAACCTAGGCTGCTGCCACCG | |
| | ck1598 | GCTAACAGGAGGAATTCCATGTTGATGCTATTCAAAGAACTCAATGAAAGTC | pCK_1099-c |
| | MT001 | TGAGACCTTTTTTTGGTCTCGGCCGCCGATACGGAAGAAATTATCTTCGATACC | |
| | MT004 | CGAGACCAAAAAAAGGTCTCACGGTAATTCCCTGACCGCCATTAAGC | |
| | ck1584 | CGGTGGCAGCAGCCTAGGTTAA | |
| pMT012 | oMT010F | AGGACGAGACCAAAAAAAGGTCTCATGGTAATTCCCTGACCGCCATTAAGC | pMT010 |
| | oMT009R | GAGACCTTTTTTTGGTCTCGTCCTCCGATACGGAAGAAATTATCTTCGATACCG | |
| pMT012_MS099 | Golden Gate | - | pMT012 |
| | | - | pMS099 |
| pWJ6 (pCEP_Kan_ *txlA*) | WJ031 | TTTGGGCTAACAGGAGGCTAGCATATGAGAATACCTGAAAGTGCGTTG | gDNA *X. indica* DSM 17382 |
| | WJ032 | TCTGCAGAGCTCGAGCATGCACATGAATATCCGATAACCATGCTGTTG | |
| | Restric. Digest NdeI | - | pCEP_Km |
| | Restric. Digest PstI-HF | - | |
| pPH4 (pCEP_Kan_ *xnpA*) | O21 | TTTGGGCTAACAGGAGGCTAGCATATGAGAAAAGCTGAGGATCA | gDNA *X. stockiae* KJ12.1 |
| | O22 | CCGTTTAAACATTTAAATCTGCAGCTGAGAGTGGTAATGTCAGCG | |
| | Restric. Digest NdeI | - | pCEP_Km |
| | Restric. Digest PstI-HF | - | |

#### Strains

**Table S10.** Overview of strains used in this work.

| **Strain** | **Reference** |
|---|---|
| *E. coli* DH10B::*mtaA* | [62] |
| *E. coli* ST18 | [46] |
| *Photorhabdus laumondii* TTO1 | [63] |
| *P. laumondii* TTO1::pCEP_Kan_*plu3263*ATG | [47] |
| *P. laumondii* TTO1::Δ*hfq*_pCEP_Kan_*plu3263* | [48] |
| *Xenorhabdus bovienii* SS-2004 | [64] |
| *X. budapestensis* DSM 16342 | [65] |
| *X. doucetiae* DSM 17909 | [66] |
| *X. doucetiae* DSM 17909::Δ*hfq* | [48] |
| *X. doucetiae* DSM 17909::*Δhfq*_pCEP_Kan_*gspS* | |
| *X. doucetiae* DSM 17909::Δ*hfq*_pCEP_Kan_paxA | |
| *X. doucetiae* DSM 17909::Δ*hfq*_pCEP_Kan_*prtA* | [45] |
| *X. doucetiae* DSM 17909::Δ*hfq*_pCEP_Kan_*xabA* | [48] |
| *X. indica* DSM 17382 | [67] |
| *X. indica* DSM 17382::Δ*hfq*_pCEP_Kan_*taxlA* | this work |
| *X. innexi* DSM 16336 | [65] |
| *X. mauleonii* DSM 17908 | [66] |
| *X. nematophila* ATCC 19061 | [68] |
| *X. nematophila* ATCC 19061::Δ*hfq*_pCEP_Kan_*XNC1_xndA* | [*48*] |
| *X. stockiae* KJ12.1 | [52] |
| *X. stockiae* KJ12.1::Δ*hfq*_pCEP_Kan_*xnpA* | this work |
| *X. szentirmaii* DSM 16338 | [65] |
| *X. szentirmaii* DSM 16338::Δ*hfq*_pCEP_Kan_*3460* | [*48*] |

#### NRPS

**Table S11.** Overview of all synthetic hybrid NRPS tested in this work.

| NRPS | Plasmids | NRPS | Plasmids |
|---|---|---|---|
| NRPS-1 | pCK_1134-a + pCK_1099-c | NRPS-2 | pCK_1134-a + pMT006 |
| NRPS-3 | pAP37_MS099 | NRPS-4 | pAP37_MS100 |
| NRPS-5 | pAP37_MS101 | NRPS-6 | pAP37_MS102 |
| NRPS-7 | pAP37_MS103 | NRPS-8 | pAP37_MS105 |
| NRPS-9 | pAP37_MS107 | NRPS-10 | pAP37_MS108 |
| NRPS-11 | pAP37_AP41 | NRPS-12 | pCK_1134-a + pMT012_MS099 |
| NRPS-13 | pMS113 + pMS114 | NRPS-14 | pMS121_MS099 + pMS114 |
| NRPS-15 | pMS113 + pMS109_MS099 | NRPS-16 | pMS121_MS099 + pMS109_MS099 |
| NRPS-17 | pMS091_MS099 + pMS114 | NRPS-18 | pMS094_MS099 + pMS114 |

#### Identification of Promiscuous A-Domains

**Table S12.** Summary of precursor-directed biosynthesis of GameXPeptide derivatives in *P. laumondii* TTO1::Δ*hfq*_pCEP_Kan_*plu3263*. Substrates as specified in Figure S1; n.d.: not detected; molecular formulas as $[M+H]^+$; Δppm: mass error.

| Product | Substrate | Production relative to 19 | Domain | Retention time [min] | Molecular formula | Δppm |
|---|---|---|---|---|---|---|
| **19b** | *p*BrF (**16**) | 6% | A3 | 10.8 | $C_{32}H_{51}BrN_5O_5$ | -0.2 |
| **19d** | *O*PrY (**13**) | 88% | A3 | 10.2 | $C_{35}H_{54}N_5O_6$ | 0.7 |
| **19c** | *p*N$_3$F (**12**) | 33% | A3 | 10.6 | $C_{32}H_{51}N_8O_5$ | 0.2 |

**Table S13.** Summary of precursor-directed biosynthesis of GameXPeptide derivatives in *X. doucetiae* DSM 17909::Δ*hfq*_pCEP_Kan_*gxpS*. Substrates as specified in Figure S1; n.d.: not detected; molecular formulas as $[M+H]^+$; Δppm: mass error.

| Product | Substrate | Production relative to 19 | Domain | Retention time [min] | Molecular formula | Δppm |
|---|---|---|---|---|---|---|
| **19c** | *p*N$_3$F (**12**) | 164% | A3 | 10.6 | $C_{32}H_{51}N_8O_5$ | 0.7 |
| **19d** | *O*PrY (**13**) | 154% | A3 | 10.2 | $C_{35}H_{54}N_5O_6$ | 1.4 |
| **19e** | Pra (**4**) | 3% | A3 | 9.4 | $C_{28}H_{48}N_5O_5$ | 1.3 |
| **19f** | HPra (**5**) | 20% | A3 | 9.6 | $C_{29}H_{50}N_5O_5$ | 1.5 |
| **19g** | *O*PrS (**3**) | 10% | A3 | 9.6 | $C_{29}H_{50}N_5O_6$ | 1.0 |
| **19h** | Aha (**9**) | 12% | A3 | 9.2 | $C_{27}H_{49}N_8O_5$ | 1.3 |
| **19i** | HPhe (**17**) | 74% | A3 | 10.6 | $C_{33}H_{54}N_5O_5$ | 0.5 |
| **19j** | PAPA (**14**) | 388% | A3 | 8.1 | $C_{32}H_{53}N_6O_5$ | 0.2 |
| **19k** | DOPA (**15**) | 2% | A3 | 9.0 | $C_{32}H_{52}N_5O_7$ | -0.9 |
| **19l** | *O*AlY (**18**) | 220% | A3 | 10.6 | $C_{35}H_{56}N_5O_6$ | 0.6 |
| **19m** | 2-Fua (**7**) | 62% | A3 | 9.6 | $C_{30}H_{50}N_5O_6$ | 0.4 |
| | Abu (**6**) | n.d. | | | | |
| | Aza (**8**) | n.d. | | | | |

**Table S14.** Summary of precursor-directed biosynthesis of szentiamide derivatives in *X. szentirmaii* DSM 16338::Δ*hfq*_pCEP_Kan_*3460*. Substrates as specified in Figure S1; n.d.: not detected; molecular formulas as $[M+H]^+$; Δppm: mass error.

| Product | Substrate | Production relative to 21 | Domain | Retention time [min] | Molecular formula | Δppm |
|---|---|---|---|---|---|---|
| **21b** | PAPA (**14**) | 42% | A5 | 7.8 | $C_{45}H_{57}N_8O_8$ | 0.3 |
| **21c** | 2-Fua (**7**) | 4% | A3 | 8.4 | $C_{43}H_{54}N_7O_{10}$ | 0.9 |
| **21d** | | 10% | A5 | 9.6 | $C_{43}H_{54}N_7O_9$ | 0.6 |
| **21e** | 5ClF (**2**) | 14% | A6 | 9.3 | $C_{45}H_{55}ClN_7O_9$ | -0.5 |
| | *p*N$_3$F (**12**) | n.d. | | | | |
| | *O*PrY (**13**) | n.d. | | | | |
| | DOPA (**15**) | n.d. | | | | |
| | *p*BrF (**16**) | n.d. | | | | |
| | HPhe (**17**) | n.d. | | | | |

**Table S15.** Summary of precursor-directed biosynthesis of xenoamicin derivatives in *X. doucetiae* DSM 17909::Δ*hfq*_pCEP_Kan_*xabA*. Substrates as specified in Figure S1; n.d.: not detected; molecular formulas as $[M+2H]^{2+}$; Δppm: mass error.

| Product | Substrate | Production relative to 22 and 23 | Domain | Retention time [min] | Molecular formula | Δppm |
|---|---|---|---|---|---|---|
| **22b** | HPra (**5**) | 4% | A4/A5/A8 | 10.0 | $C_{65}H_{109}N_{13}O_{15}$ | 5.8 |
| **22c** | Aha (**9**) | 0.4% | A4/A5/A8 | 10.2 | $C_{63}H_{108}N_{16}O_{15}$ | 6.4 |
| **22d** | Aze (**1**) | 26% | A12 | 10.1 | $C_{64}H_{111}N_{13}O_{15}$ | 5.1 |
| **22e** | | 34% | A1 | 10.5 | $C_{64}H_{111}N_{13}O_{15}$ | 5.3 |
| **22f** | | 239% | A1 + A12 | 9.5 | $C_{62}H_{107}N_{13}O_{15}$ | 6.2 |
| | *O*PrS (**3**) | n.d. | | | | |
| | Pra (**4**) | n.d. | | | | |
| | Abu (**6**) | n.d. | | | | |
| | Aza (**8**) | n.d. | | | | |

**Table S16.** Summary of precursor-directed biosynthesis of xeneprotide derivatives in *X. stockiae* KJ12.1::Δ*hfq*_pCEP_Kan_*09200*. Substrates as specified in Figure S1; n.d.: not detected; molecular formulas as $[M+H]^+$; Δppm: mass error.

| Product | Substrate | Production relative to 27 | Domain | Retention time [min] | Molecular formula | Δppm |
|---|---|---|---|---|---|---|
| **27b/27c** | *O*PrY (**14**) | 9% | A2/A3 | 9.0 | $C_{40}H_{42}N_5O_7$ | 0.4 |
| **27e/27f** | *p*$N_3$F (**7**) | 10% | A2/A3 | 9.3 | $C_{37}H_{39}N_8O_6$ | 1.5 |
| **27h/27i** | *p*BrF (**18**) | 1% | A2/A3 | 9.5 | $C_{37}H_{39}BrN_5O_6$ | 1.6 |
| **27k/27l** | HPhe (**19**) | 1% | A2/A3 | 9.2 | $C_{38}H_{42}N_5O_6$ | 0.8 |
| **27n/27o** | PAPA (**16**) | 3% | A2/A3 | 6.9 | $C_{37}H_{41}N_6O_6$ | 0.6 |
| **27q** | Aze (**25**) | 1234% | A4 | 8.4 | $C_{38}H_{39}N_6O_6$ | 0.3 |
| | 5ClW (**15**) | n.d. | | | | |
| | 2-Fua (**24**) | n.d. | | | | |

**Table S17.** Summary of precursor-directed biosynthesis of taxlllaid derivatives in *X. indica* DSM 17382::Δ*hfq*_pBAD_*XINDV2_06865*. Substrates as specified in Figure S1; n.d.: not detected; molecular formulas as $[M+H]^+$; Δppm: mass error.

| Product | Substrate | Relative Production to 28 | Domain | Retention time [min] | Molecular formula | Δppm |
|---|---|---|---|---|---|---|
| **28b** | *p*$N_3$F (**12**) | 11% | A3 | 10.8 | $C_{43}H_{69}N_{10}O_9$ | -1.9 |
| **28c** | PAPA (**14**) | 92% | A3 | 8.7 | $C_{43}H_{71}N_8O_9$ | -2.5 |
| **28d** | 2-Fua (**7**) | 2% | A3 | 10.3 | $C_{41}H_{68}N_7O_{10}$ | -2.0 |
| | *O*PrS (**3**) | n.d. | | | | |
| | Pra (**4**) | n.d. | | | | |
| | HPra (**5**) | n.d. | | | | |
| | Abu (**6**) | n.d. | | | | |
| | Aza (**8**) | n.d. | | | | |
| | Aha (**9**) | n.d. | | | | |
| | *O*PrY (**13**) | n.d. | | | | |
| | DOPA (**15**) | n.d. | | | | |
| | *p*BrF (**16**) | n.d. | | | | |
| | HPhe (**17**) | n.d. | | | | |

**Table S18.** Summary of precursor-directed biosynthesis of fitayylide derivatives in *E. coli* DH10B::*mtaA*_pCK_0683. Substrates as specified in Figure S1; n.d.: not detected; Δppm: molecular formulas as $[M+H]^+$; mass error.

| Product | Substrate | Production relative to 29 | Domain | Retention time [min] | Molecular formula | Δppm |
|---|---|---|---|---|---|---|
| **29b** | *p*BrF (**16**) | 2% | A5 | 8.4 | $C_{39}H_{54}BrN_6O_9$ | 0.3 |
| **29c** | *O*PrY (**13**) | 2% | A5 | 8.0 | $C_{42}H_{57}N_6O_{10}$ | 0.8 |
| **29d** | PAPA (**14**) | 0.8% | A5 | 5.9 | $C_{39}H_{56}N_7O_9$ | 0.5 |
| **29e** | | 1% | A4 | 6.0 | $C_{39}H_{56}N_7O_9$ | 0.6 |
| | 2-Fua (**7**) | n.d. | | | | |
| | *p*N$_3$F (**12**) | n.d. | | | | |
| | DOPA (**15**) | n.d. | | | | |
| | HPhe (**17**) | n.d. | | | | |

**Table S19.** Summary of precursor-directed biosynthesis of xenortide derivatives in *X. nematophila* ATCC 19061::Δ*hfq*_pCEP_Kan_*XNC1_xndA*. Substrates as specified in Figure S1; n.d.: not detected; molecular formulas as $[M+H]^+$; Δppm: mass error.

| Product | Substrate | Relative Production to 32 | Domain | Retention time [min] | Molecular formula | Δppm |
|---|---|---|---|---|---|---|
| **32h** | 2-Fua (**24**) | 47% | A2 | 6.8 | $C_{23}H_{34}N_3O_3$ | -1.7 |
| | *p*N$_3$F (**12**) | n.d. | | | | |
| | *O*PrY (**13**) | n.d. | | | | |
| | PAPA (**14**) | n.d. | | | | |
| | DOPA (**15**) | n.d. | | | | |
| | *p*BrF (**16**) | n.d. | | | | |
| | HPhe (**17**) | n.d. | | | | |

**Table S20.** Summary of precursor-directed biosynthesis of protegomycin derivatives in *X. doucetiae* DSM 17909::Δ*hfq*_pCEP_Kan_*prtA*. Substrates as specified in Figure S1; n.d.: not detected; molecular formulas as $[M+2H]^{2+}$; Δppm: mass error.

| Product | Substrate | Production relative to 34 | Domain | Retention time [min] | Molecular formula | Δppm |
|---|---|---|---|---|---|---|
| **34b** | *O*PrY (**13**) | 15% | A6 | 8.1 | $C_{60}H_{71}N_7O_{12}$ | 1.7 |
| **34c** | | 23% | A2/3/4/5 | 8.7 | $C_{62}H_{72}N_8O_{11}$ | 2.3 |
| **34d** | PAPA (**14**) | 56% | A6 | 6.5 | $C_{57}H_{70}N_8O_{11}$ | 1.2 |
| **34e** | | 328% | A2/3/4/5 | 7.2 | $C_{59}H_{71}N_9O_{10}$ | 1.6 |
| **34i** | DOPA (**15**) | 40% | A2/3/4/5 | 7.8 | $C_{59}H_{70}N_8O_{12}$ | 1.3 |
| | *p*N$_3$F (**12**) | n.d. | | | | |
| | HPhe (**17**) | n.d. | | | | |

**Table S21.** Overview of all single incorporations of non-cognate building blocks detected in precursor-directed biosynthesis. Grouped after substrates specified in Figure S1; n.d.: not detected.

| Substrate | NRPS | Domain | Compound | Rel. incorporation (%) |
|---|---|---|---|---|
| Aze (**1**) | XabABC | A1 | **22e** | 34 |
| | | A12 | **22d** | 26 |
| | XnpAB | A4 | **27q** | 1234 |
| 5ClW (**2**) | XszeS | A6 | **21e** | 14 |
| *O*PrS (**3**) | GxpS | A3 | **19g** | 10 |
| Pra (**4**) | GxpS | A3 | **19e** | 3 |
| HPra (**5**) | GxpS | A3 | **19f** | 20 |
| Abu (**6**) | n.d. | | | |
| 2-Fua (**7**) | GxpS | A3 | **19m** | 61 |
| | XszeS | A3 | **21c** | 4 |
| | | A5 | **21d** | 10 |
| | TxlAB | A3 | **21d** | 2 |
| | XndAB | A2 | **32h** | 47 |
| Aza (**8**) | n.d. | | | |
| Aha (**9**) | GxpS | A3 | **19m** | 12 |
| | Xab | A4/A5/A8 | **22c** | 0.4 |
| $N_3$Orn (**10**) | n.d. | | | |
| $N_3$Lys (**11**) | n.d. | | | |
| *p*$N_3$F (**12**) | TxlAB | A3 | **28b** | 11 |
| | GxpS | A3 | **19c** | 164 |
| | XnpAB | A2/A3 | **27b**/**27c** | 10 |
| *O*PrY (**13**) | GxpS | A3 | **19d** | 154 |
| | FitAB | A5 | **30c** | 2 |
| | XnpAB | A2/A3 | **27e**/**27f** | 9 |
| | PrtAB | A6 | **34b** | 15 |
| | | A2/A3/A4/A5 | **34c** | 56 |
| PAPA (**14**) | XszeS | A5 | **21b** | 41 |
| | TxlAB | A3 | **28c** | 92 |
| | GxpS | A3 | **19j** | 388 |
| | FitAB | A5 | **30d** | 0.8 |
| | | A4 | **30e** | 1 |
| | XnpAB | A2/A3 | **27n**/**27o** | 3 |
| | PrtAB | A6 | **34d** | 56 |
| | | A2/A3/A4/A5 | **34e** | 328 |
| DOPA (**15**) | GxpS | A3 | **19k** | 2 |
| | PrtAB | A2/A3/A4/A5 | **34i** | 40 |
| *p*BrF (**16**) | FitAB | A5 | **30b** | 2 |
| | XnpAB | A2/A3 | **27h**/**27i** | 1 |
| HPhe (**17**) | GxpS | A3 | **19i** | 74 |
| | XnpAB | A2/A3 | **27k**/**27l** | 1 |
| *O*AlY (**18**) | GxpS | A3 | **19l** | 220 |

#### NRPS Engineering with Promiscuous A-Domains

**Table S22.** Overview of the non-cognate amino acids supplemented to production medium of the synthetic hybrid NRPS-3 and NRPS-5 to NRPS-10. +: incorporation; -: not detected

| NRPS | Suppl. building blocks | Incorporation |
|---|---|---|
| NRPS-3 | *p*$N_3$F (**12**) | + |
| | *O*PrY (**13**) | + |
| NRPS-5 | **13** | - |
| NRPS-6 | **12** | + |
| | **13** | - |
| NRPS-7 | **12** | + |
| | **13** | + |
| NRPS-8 | 5ClW (**2**) | + |
| NRPS-9 | **12** | - |
| | **13** | - |
| | *p*BrF (**16**) | - |
| NRPS-10 | Aha (**9**) | - |
| | HPra (**5**) | - |

### Figures

#### Identification of Promiscuous A-Domains

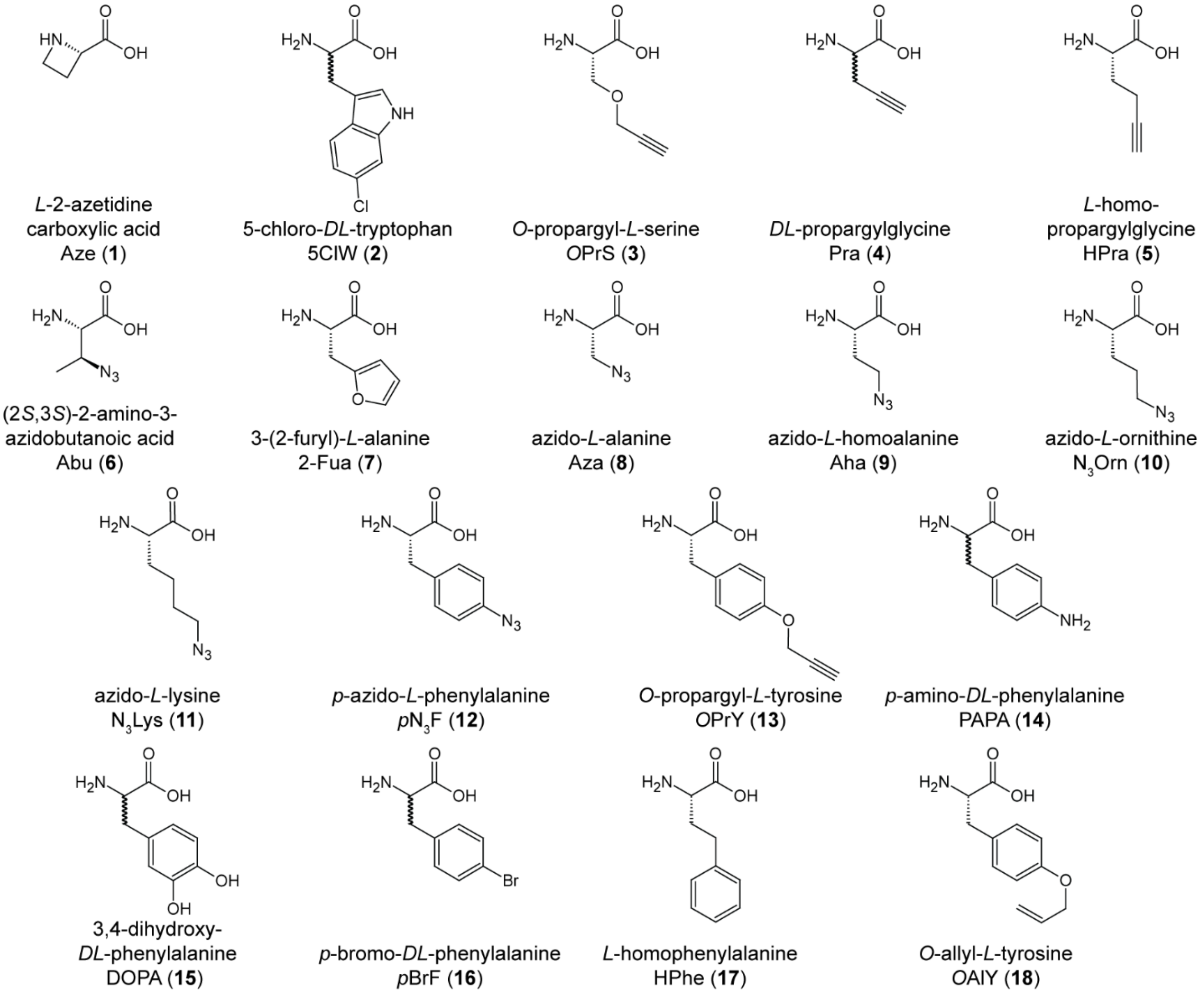

**Figure S1.** Overview of all non-cognate amino acids used for precursor-directed biosynthesis.

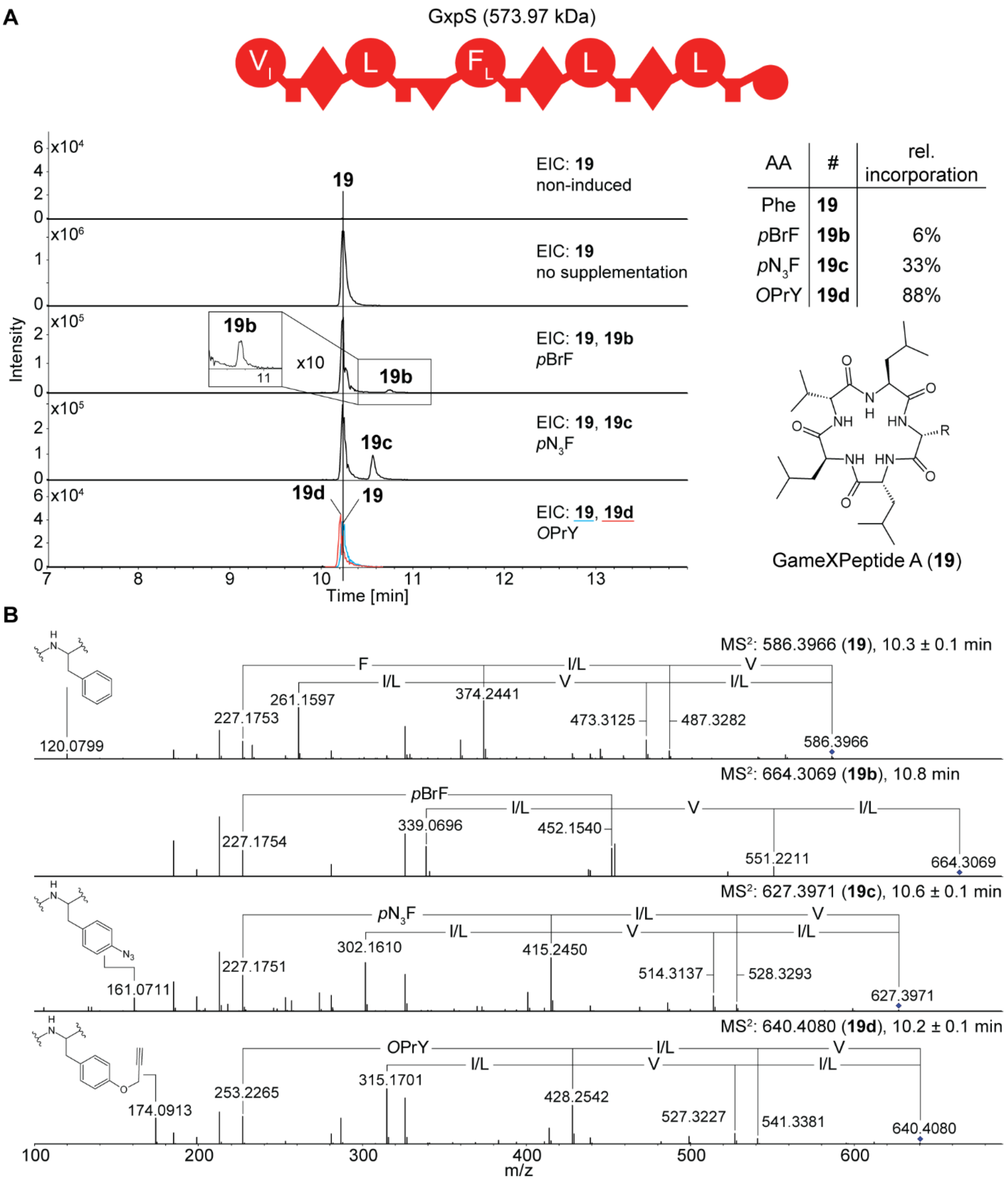

**Figure S2.** Precursor-directed biosynthesis of GameXPeptide derivatives. **A.** Schematic depiction of the NRPS GxpS from *P. laumondii* TTO1; structures and extracted ion chromatograms (EIC) of produced GameXPeptide derivatives **19-19d** identified by supplementing the production medium with indicated non-cognate amino acids. A-domain specificity indicated by the amino acid one-letter code. **B.** $MS^2$ fragmentation spectra of GameXPeptide A (**19**) and its derivatives **19b-19d**.

**A**

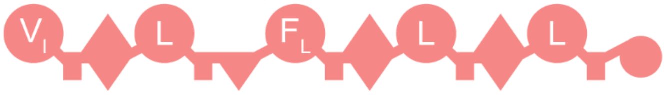

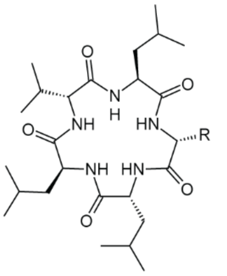

GameXPeptide A (**19**)

| AA | # | rel. incorporation | AA | # | rel. incorporation |
|---|---|---|---|---|---|
| Phe | **19** | | *O*PrS | **19g** | 10% |
| *p*N$_3$F | **19c** | 164% | Aha | **19h** | 12% |
| OPrY | **19d** | 154% | HPhe | **19i** | 74% |
| Pra | **19e** | 3% | PAPA | **19j** | 388% |
| HPra | **19f** | 20% | DOPA | **19k** | 2% |

**B**

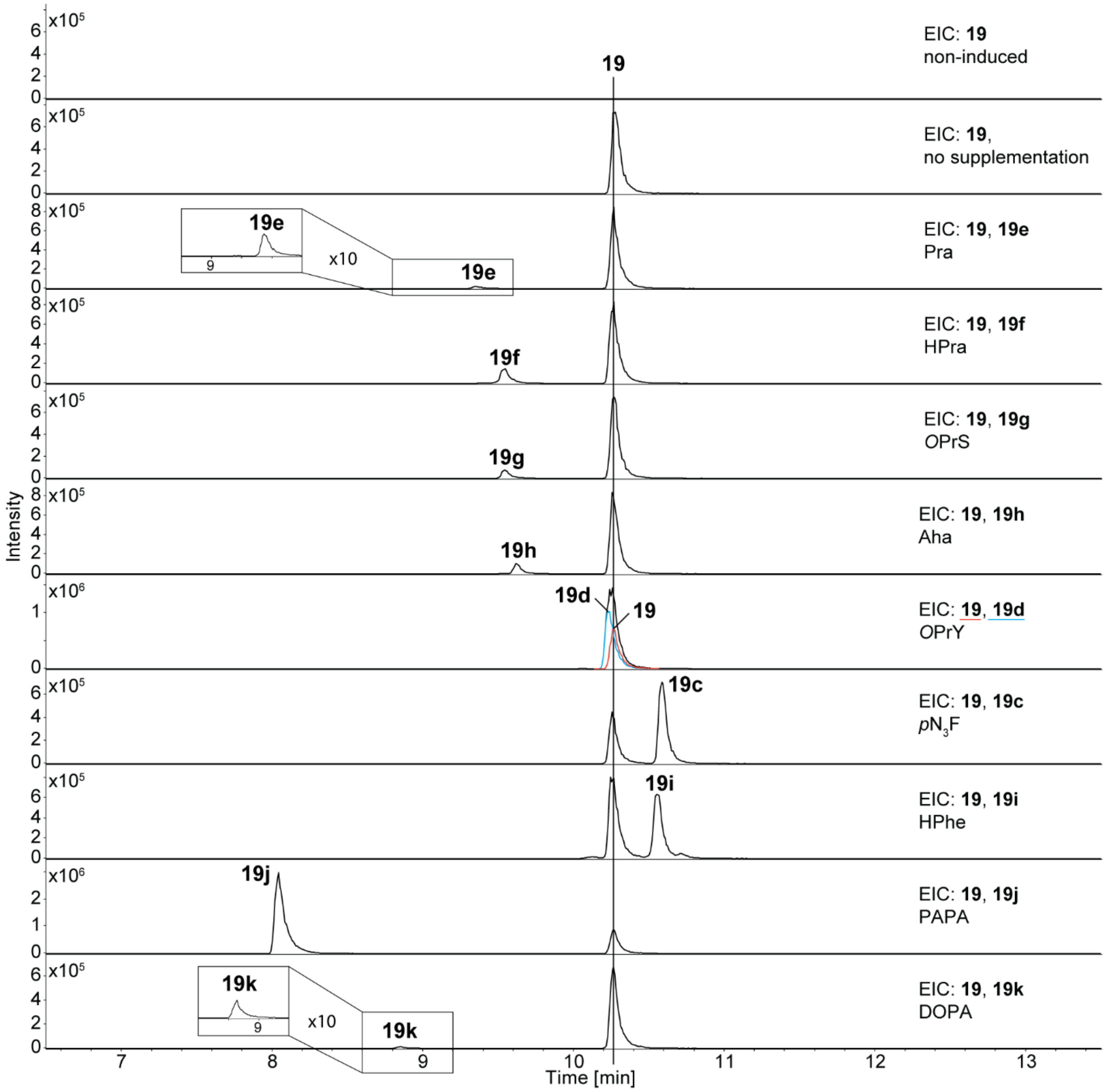

**Figure S3.** Precursor-directed biosynthesis of GameXPeptide derivatives. **A.** Schematic depiction of the NRPS GxpS from *X. doucetiae* DSM 17909 and produced GameXPeptide derivatives **19c**-**19k** identified by supplementing the production medium with indicated non-cognate amino acids. A-domain specificity indicated by the amino acid one-letter code. **B.** Extracted ion chromatograms (EIC) of the GameXPeptide derivatives **19c**-**19k**.

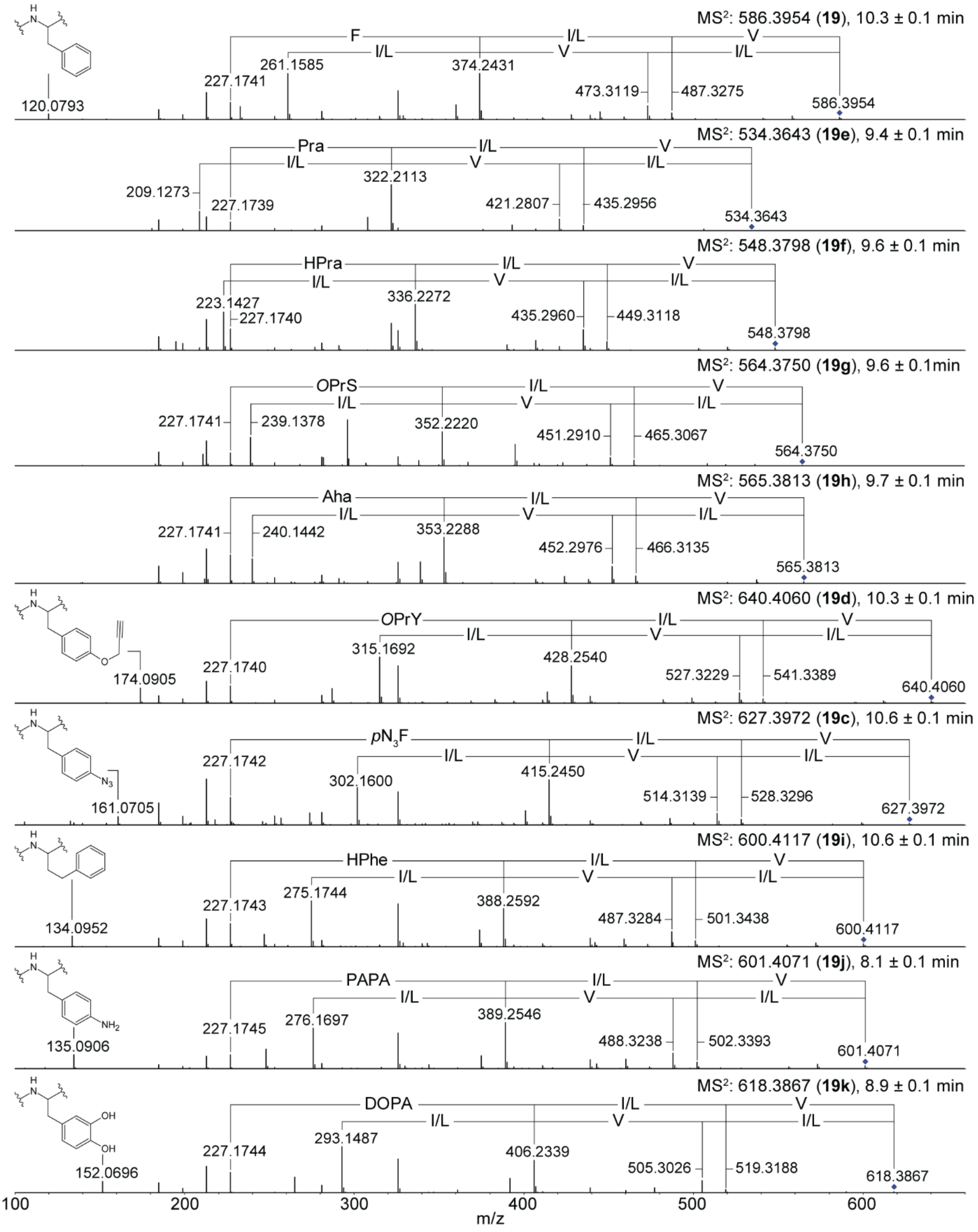

**Figure S4.** $MS^2$ fragmentation spectra of GameXPeptide derivatives **19c-19f** depicted in Figure S3 produced by precursor-directed biosynthesis compared to natural GameXPeptide A (**19**).

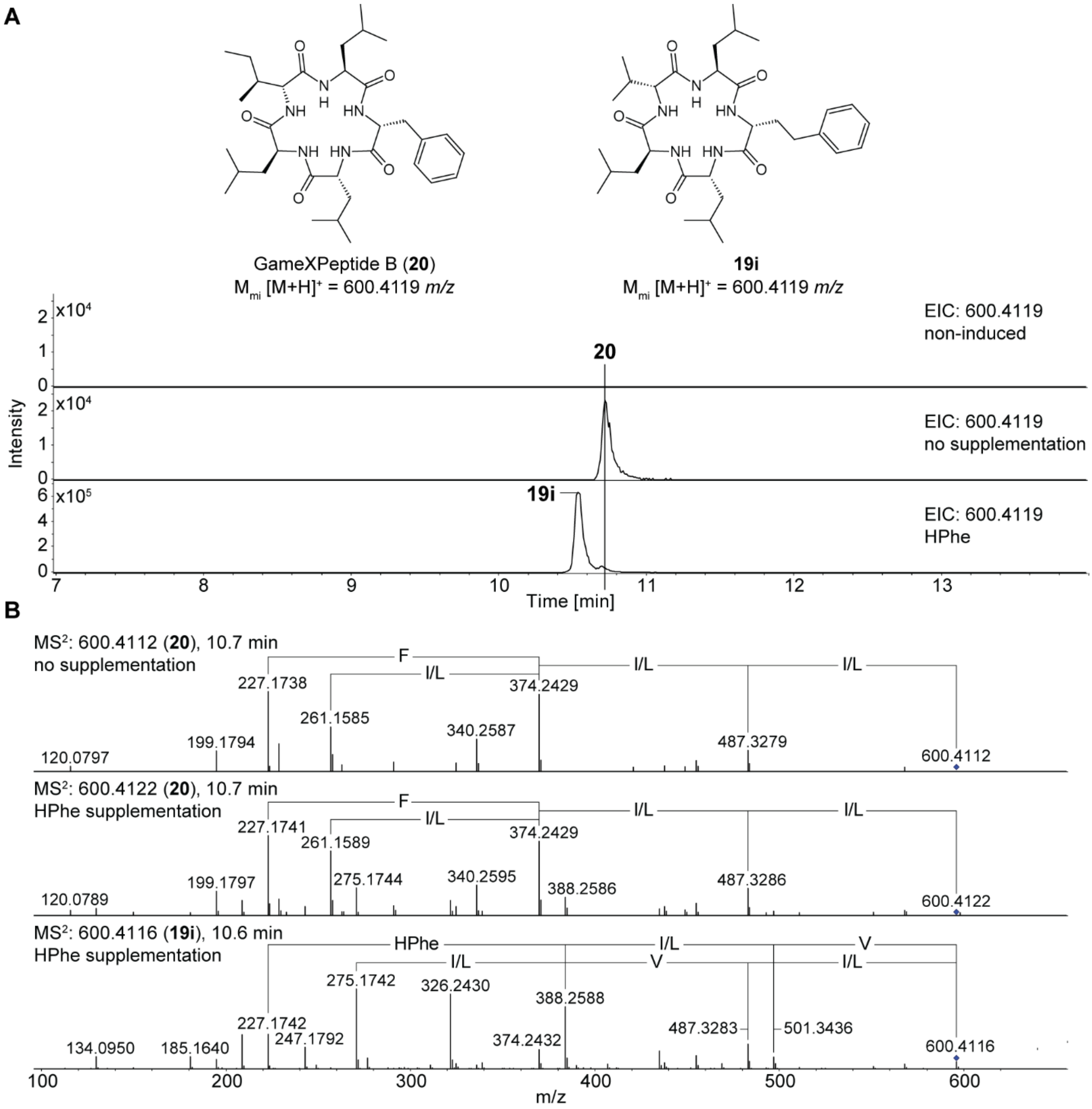

**Figure S5.** HPLC-HR-MS comparison between GameXPeptide B (**20**) and **19i**. **A.** Structure and extracted ion chromatograms (EIC) of **20** and **19i**. **B.** $MS^2$ fragmentation spectra of the GameXPeptide derivative **19i** produced by precursor-directed biosynthesis compared to natural GameXPeptide B (**20**).

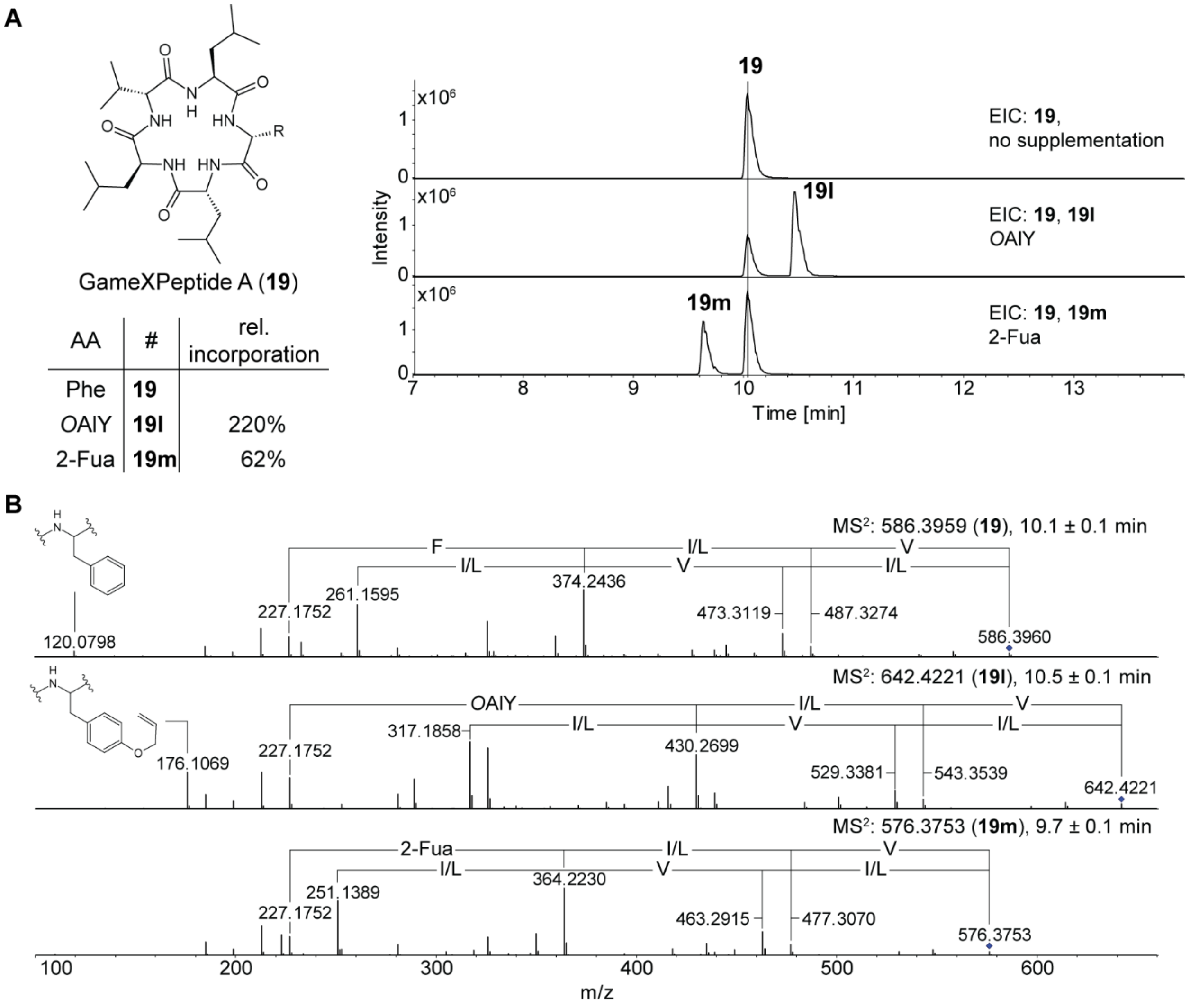

**Figure S6.** Precursor-directed biosynthesis of GameXPeptide derivatives. **A.** Structures and extracted ion chromatograms (EIC)of produced GameXPeptide derivatives **19l** and **19m** identified by supplementing the production medium with *O*AlY (**27**) and Fua (**7**). **B.** MS$^2$ fragmentation spectra of GameXPeptide derivatives **19l** and **19m** compared to natural GameXPeptide A (**19**).

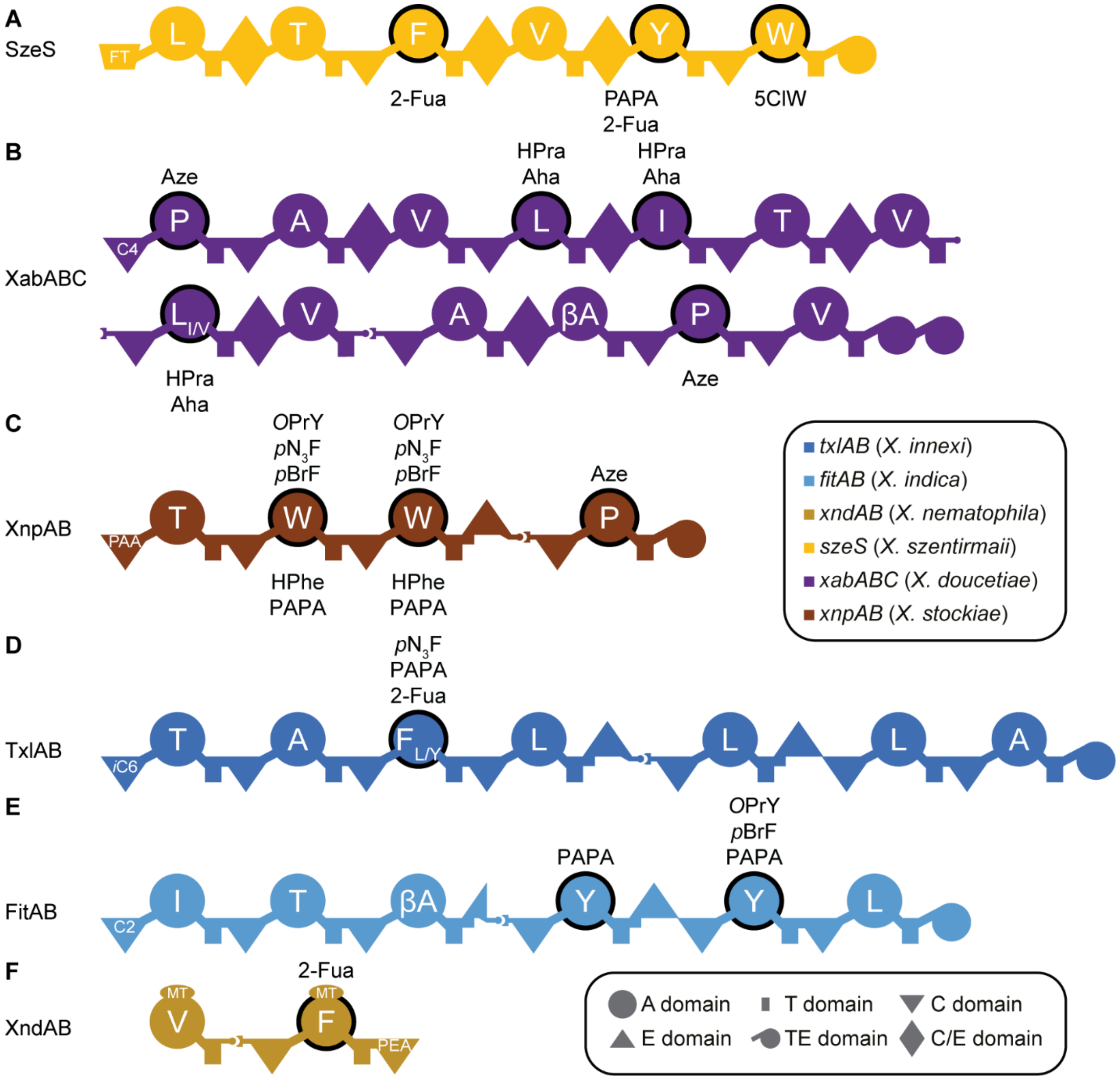

**Figure S7.** Schematic depiction of the organization of the NRPS producing the compounds shown in Figure 2. Promiscuous A-domains accepting indicated ncAAs with a black circle. Szentimide synthetase (SzeS, **A**) from *X. szentirmaii* DSM 16338, xenoamicin synthetase (XabABC, **B**) from *X. doucetiae* DSM 17909, xeneprotide synthetase (XnpAB, **C**) from *X. stockiae* KJ12.1, taxlllaid synthetase (TxlAB, **D**) from *X. inica* DSM 17382, fitayylide synthetase (FitaAB, **E**) from *X. innexi* DSM 16336, and xenortide synthetase (XndAB, **F**) from *X. nematophila* ATCC 19061.

**A**

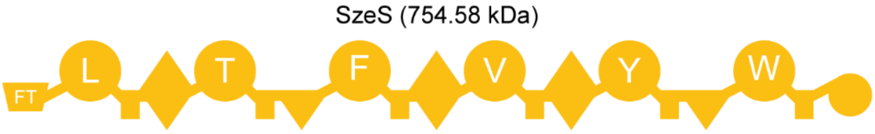

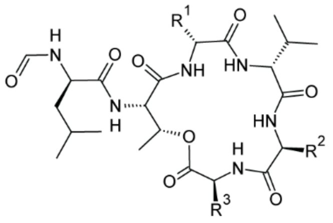

szentiamide (**21**)

| AA | R | # | rel. incorporation |
|---|---|---|---|
| Phe, Tyr, Trp | 1, 2, 3 | **21** | |
| PAPA | 2 | **21b** | 42% |
| 2-Fua | 1 / 2 | **21c/d** | 4% / 10% |
| 5ClW | 3 | **21e** | 14% |

**B**

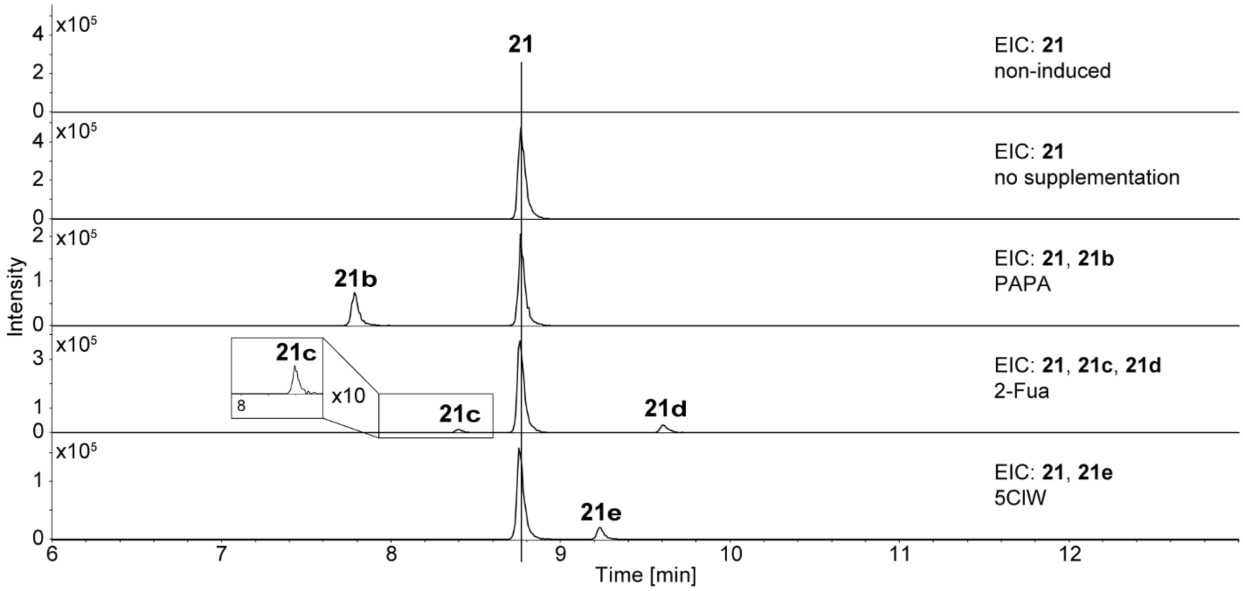

**Figure S8.** Precursor-directed biosynthesis of szentiamide derivatives. **A.** Schematic depiction of the NRPS SzeS from *X. szentirmaii* DSM 16338 and produced szentiamide derivatives **21**-**21e** identified by supplementing the production medium with indicated non-cognate amino acids. A-domain specificity indicated by the amino acid one-letter code. **B.** Extracted ion chromatograms (EIC) of the szentiamide derivatives **21**-**21e**.

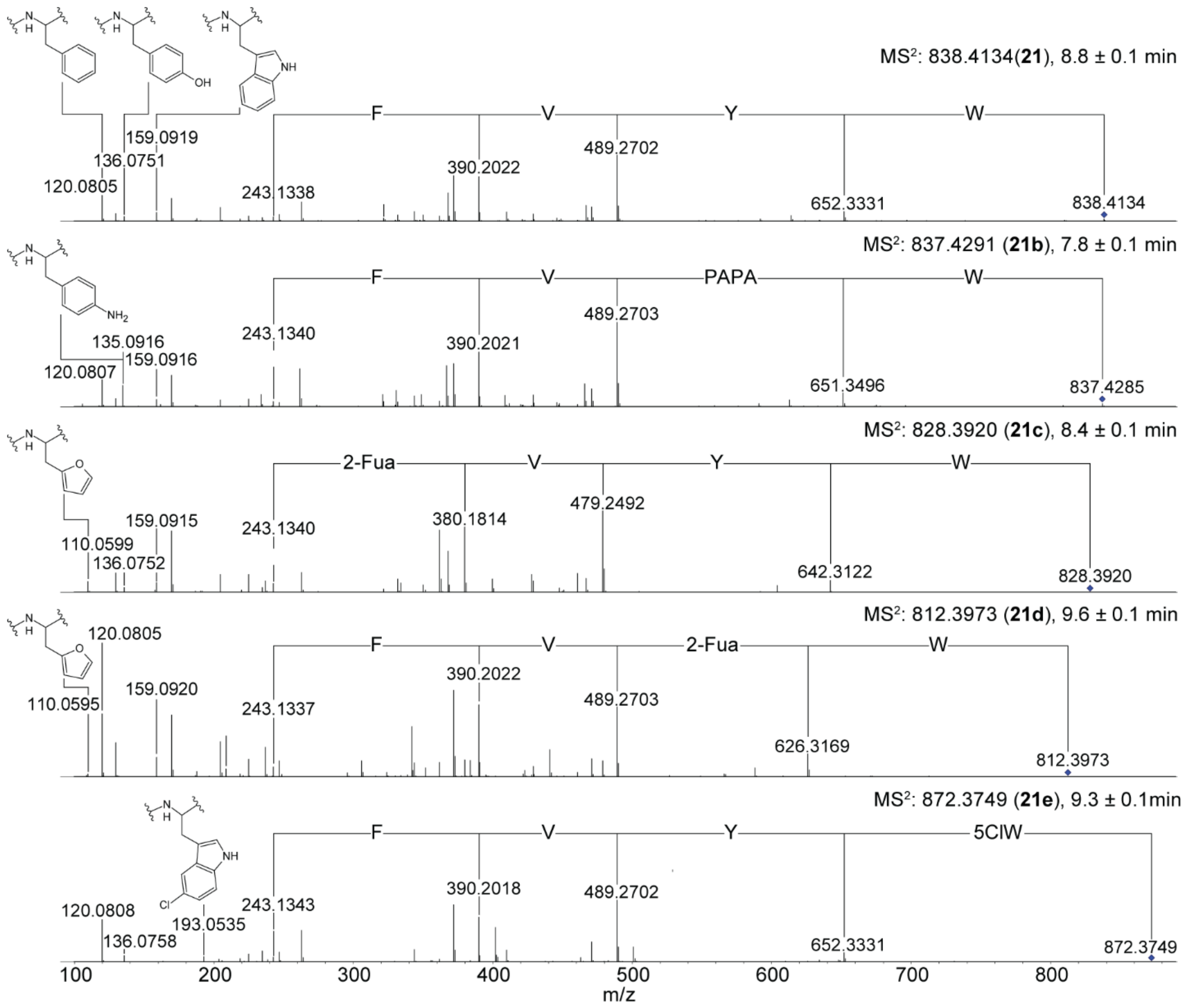

**Figure S9.** $MS^2$ fragmentation spectra of szentiamide derivatives **21b**-**21d** depicted in Figure S8 produced by precursor-directed biosynthesis compared to natural szentiamide (**21**).

**A**

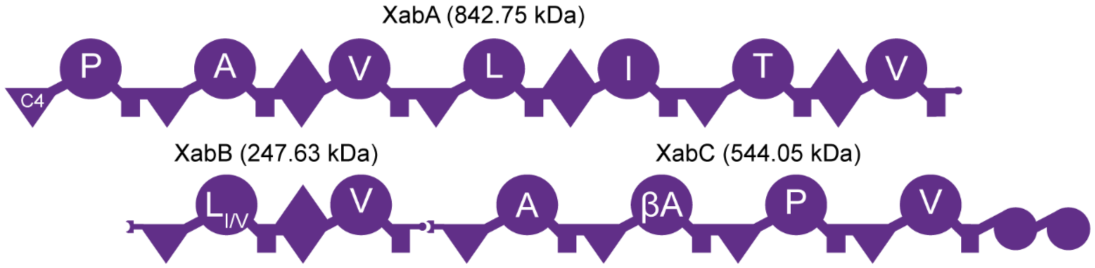

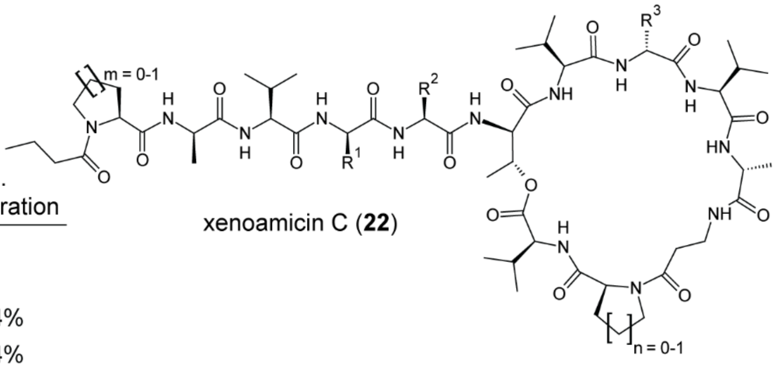

| AA | R | m | n | # | rel. incorporation |
|---|---|---|---|---|---|
| Leu, Ile, Leu | 1, 2, 3 | 1 | 1 | **22** | |
| 3x Leu | 1, 2, 3 | 1 | 1 | **23** | |
| HPra | 1 / 2 / 3 | 1 | 1 | **22b** | 4% |
| Aha | 1 / 2 / 3 | 1 | 1 | **22c** | 0.4% |
| Aze | | 1 | 0 | **22d** | 26% |
| Aze | | 0 | 1 | **22e** | 34% |
| Aze | | 0 | 0 | **22f** | 239% |

**B**

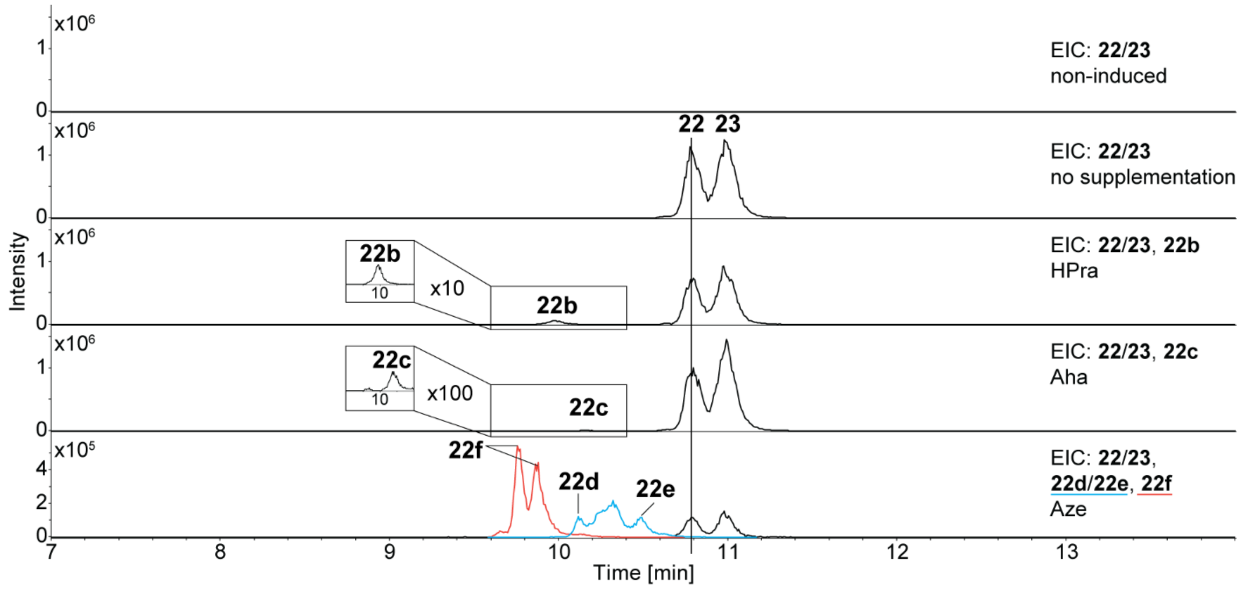

**Figure S10.** Precursor-directed biosynthesis of xenoamicin derivatives. **A.** Schematic depiction of the NRPS XabABC from *X. doucetiae* DSM 17909 and produced xenoamicin derivatives **22-22f** identified by supplementing the production medium with indicated non-cognate amino acids. A-domain specificity indicated by the amino acid one-letter code. βA: beta-alanine. **B.** Extracted ion chromatograms (EIC) of the xenoamicin derivatives **22-22f** and **23**.

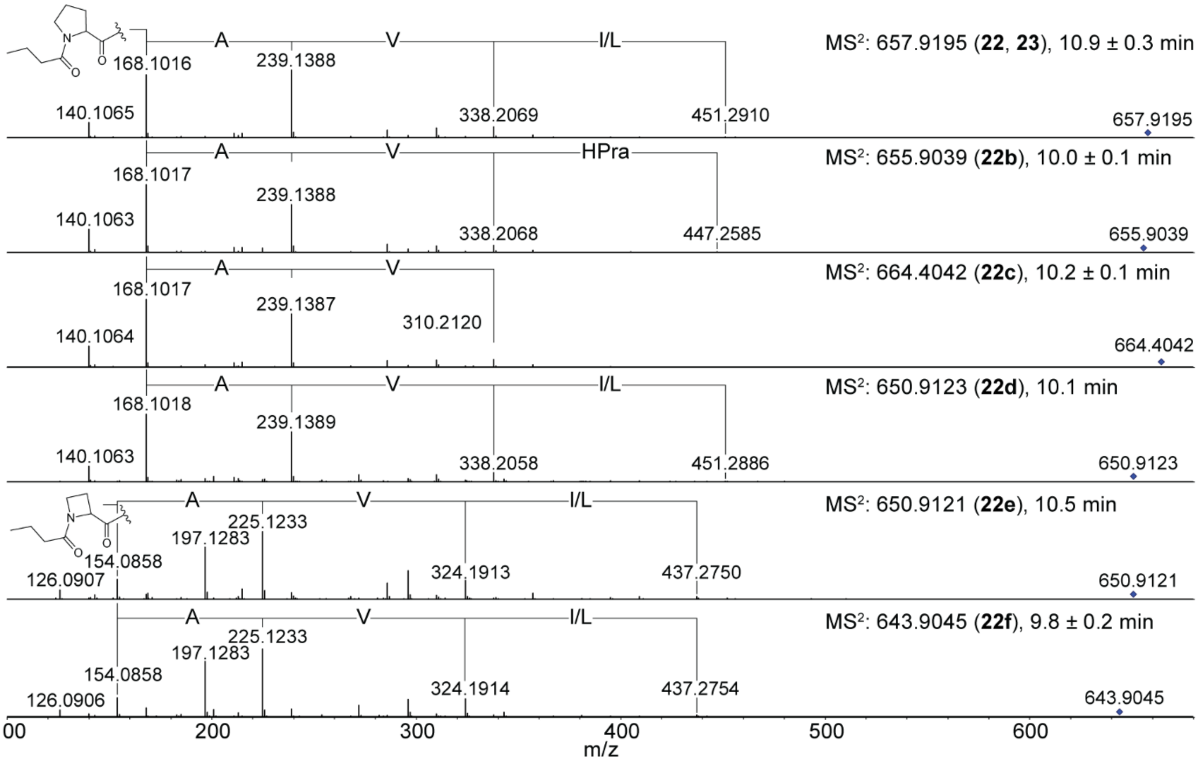

**Figure S11**. $MS^2$ fragmentation spectra of xenoamicin C (**22**) and B (**23**) and their derivatives **22b-22f**, depicted in Figure S10, produced by precursor-directed biosynthesis.

**A**

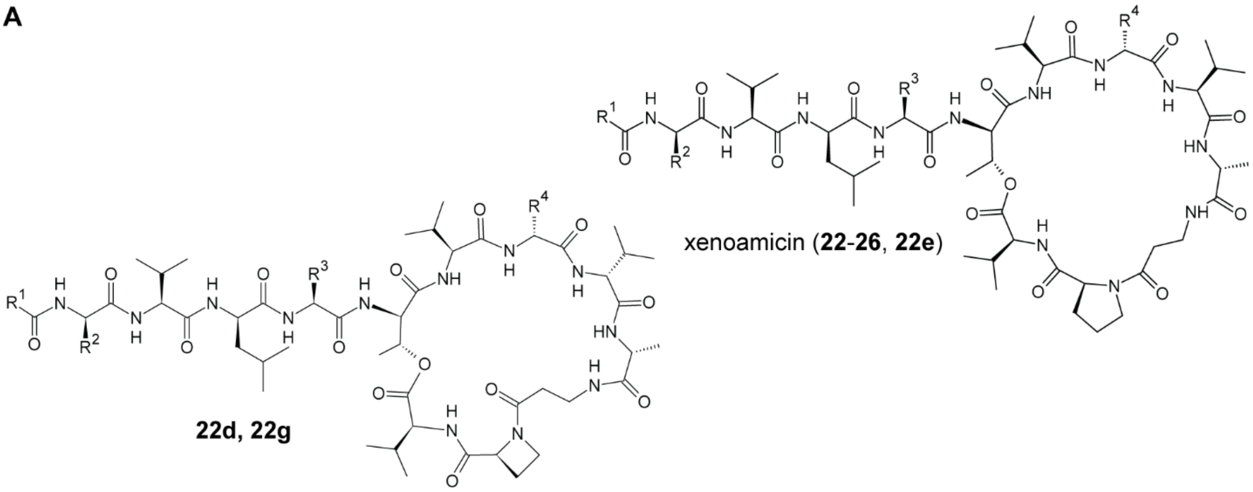

**B**

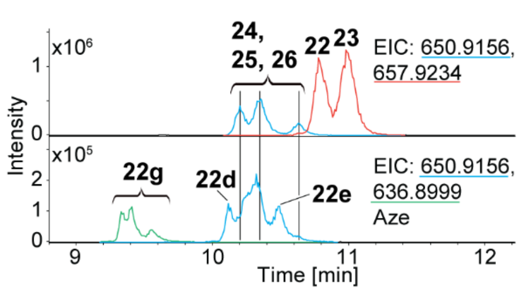

| | $R^1$ | $R^2$ | $R^3$ | $R^4$ | $M_{mi}$ $[M+2H]^{2+}$ |
|---|---|---|---|---|---|
| **22** | 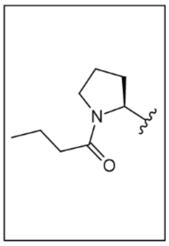 | Me | *s*Bu | *s*Bu | 657.9237 *m/z* |
| **23** | | Me | *s*Bu | *i*Bu | 657.9237 *m/z* |
| **24** | | Me | *s*Bu | *i*Pr | 650.9156 *m/z* |
| **25** | | H | *s*Bu | *s*Bu / *i*Bu | 650.9156 *m/z* |
| **26** | | Me | *i*Pr | *s*Bu / *i*Bu | 650.9156 *m/z* |
| **22d** | | Me | *s*Bu | *s*Bu / *i*Bu | 650.9156 *m/z* |
| **22e** | | Me | *s*Bu | *s*Bu / *i*Bu | 650.9156 *m/z* |
| **22g** | 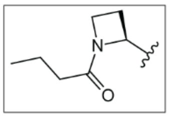 | Me / H | *s*Bu / *i*Pr | *i*Pr / *s*Bu / *i*Bu | 636.8999 *m/z* |

**C**

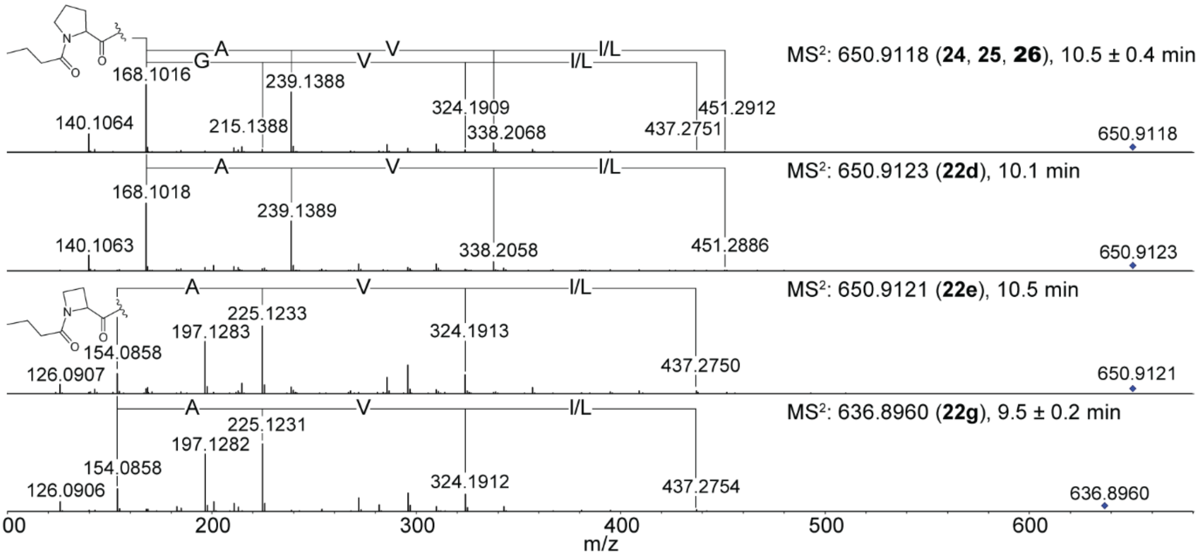

**Figure S12.** HPLC-HR-MS comparison between natural xenoamicin C, B, A, F, and E (**22-26**) and the derivatives **22d**, **22e**, and **22g** produced by precursor-directed biosynthesis. **A.** Structure of xenoamicins **22-26** and the derivatives **22d**, **22e**, and **22g**. **B.** Extracted ion chromatograms (EIC) of **22-26** and derivatives **22d**, **22e**, and **22g** produced by precursor-directed biosynthesis supplementing Aze (**1**) to the production medium. **C.** $MS^2$ fragmentation spectra of the xenoamicin derivative **22d**, **22e**, and **22g** compared to natural xenoamicin A, F, E (**24-26**).

A

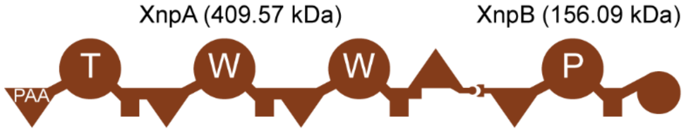

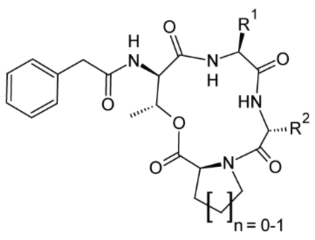

xeneprotide A (**27**)

| AA | R | n | # | rel. incorporation | AA | R | # |
|---|---|---|---|---|---|---|---|
| Trp, Trp | 1, 2 | 1 | **27** | | | | |
| OPrY | 1 / 2 | 1 | **27b/c** | 9% | OPrY | 1, 2 | **27d** |
| *p*N$_3$F | 1 / 2 | 1 | **27e/f** | 10% | *p*N$_3$F | 1, 2 | **27g** |
| *p*BrF | 1 / 2 | 1 | **27h/i** | 1% | *p*BrF | 1, 2 | **27j** |
| HPhe | 1 / 2 | 1 | **27k/l** | 1% | HPhe | 1, 2 | **27m** |
| PAPA | 1 / 2 | 1 | **27n/o** | 3% | PAPA | 1, 2 | **27p** |
| Aze | | 0 | **27q** | 1234% | | | |

**B**

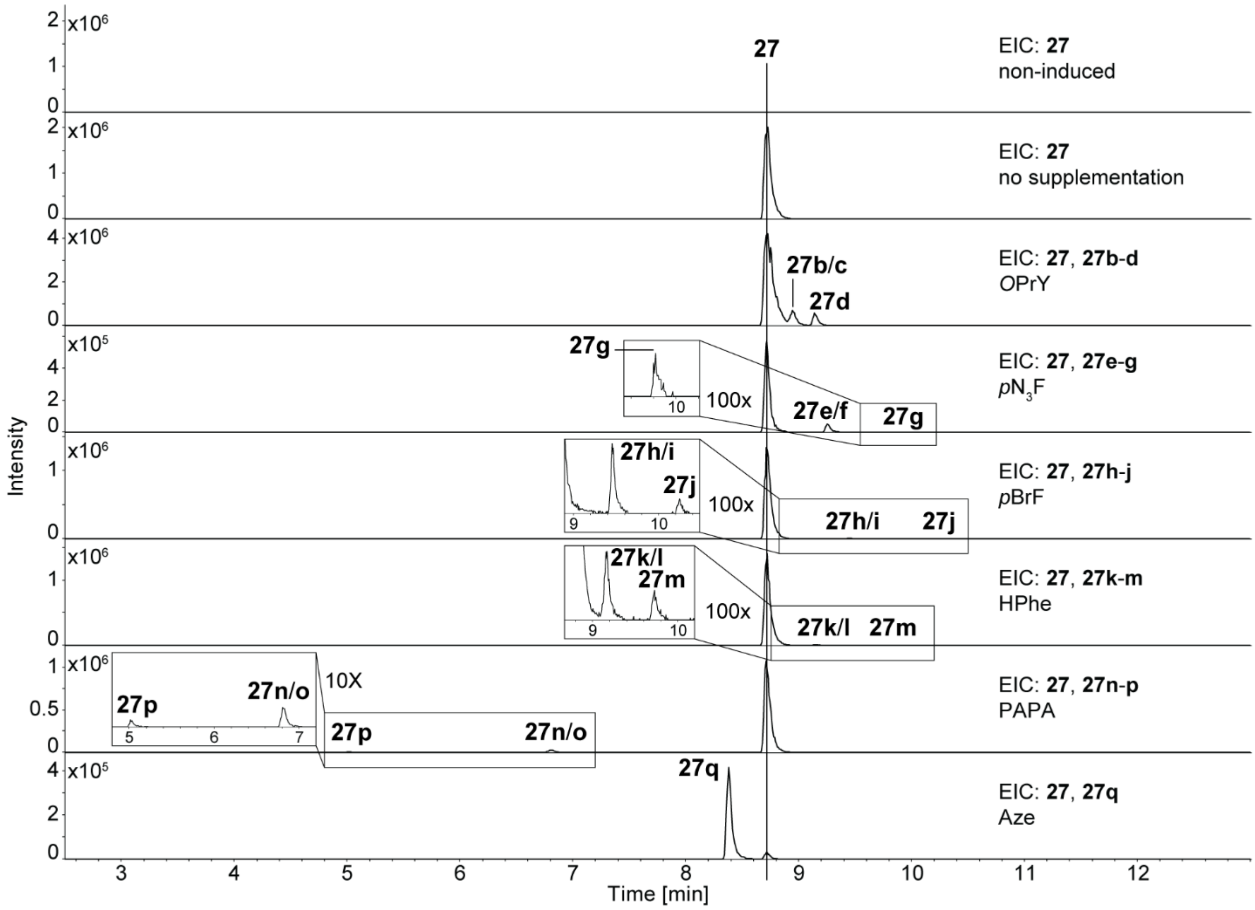

**Figure S13.** Precursor-directed biosynthesis of xeneprotide derivatives. **A.** Schematic depiction of the NRPS XnpAB from *X. stockiae* KJ12.1 and produced xeneprotide derivatives **27-27q** identified by supplementing the production medium with indicated non-cognate amino acids. A-domain specificity indicated by the amino acid one-letter code. **B.** Extracted ion chromatograms (EIC) of the xeneprotide derivatives **27-27q**.

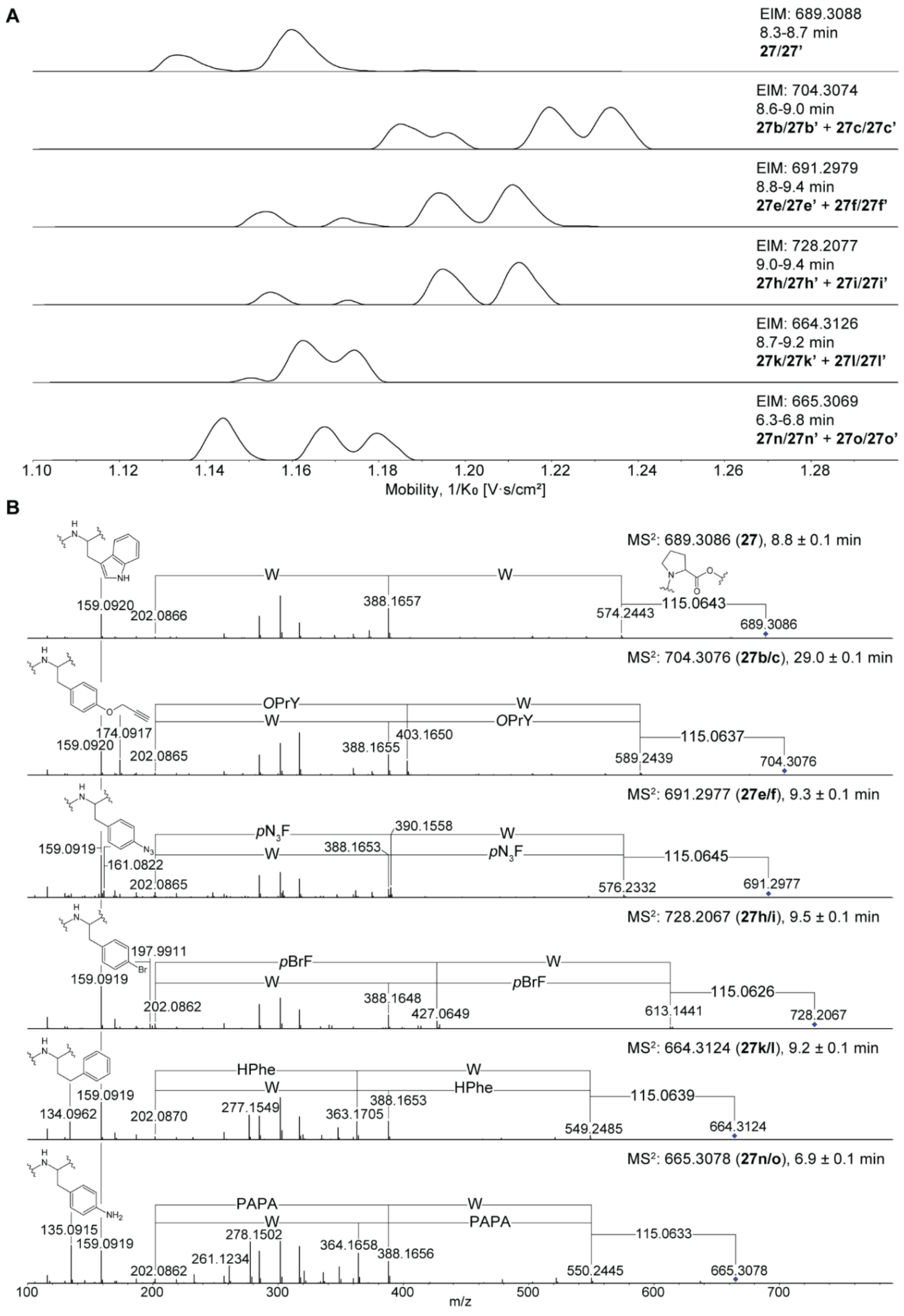

**Figure S14.** Detailed analysis of single-exchange derivatives of xeneprotide generated by precursor-directed biosynthesis. **A.** Extracted ion mobilograms (EIM) of xeneprotide derivatives **27b**/**c**, **27e**/**f**, **27h**/**i**, **27k**/**l**, and **27n**/**o**. Isomers indicated with apostrophes. **B.** $MS^2$ fragmentation spectra of the xeneprotide derivatives harboring single exchanges depicted in Figure S13.

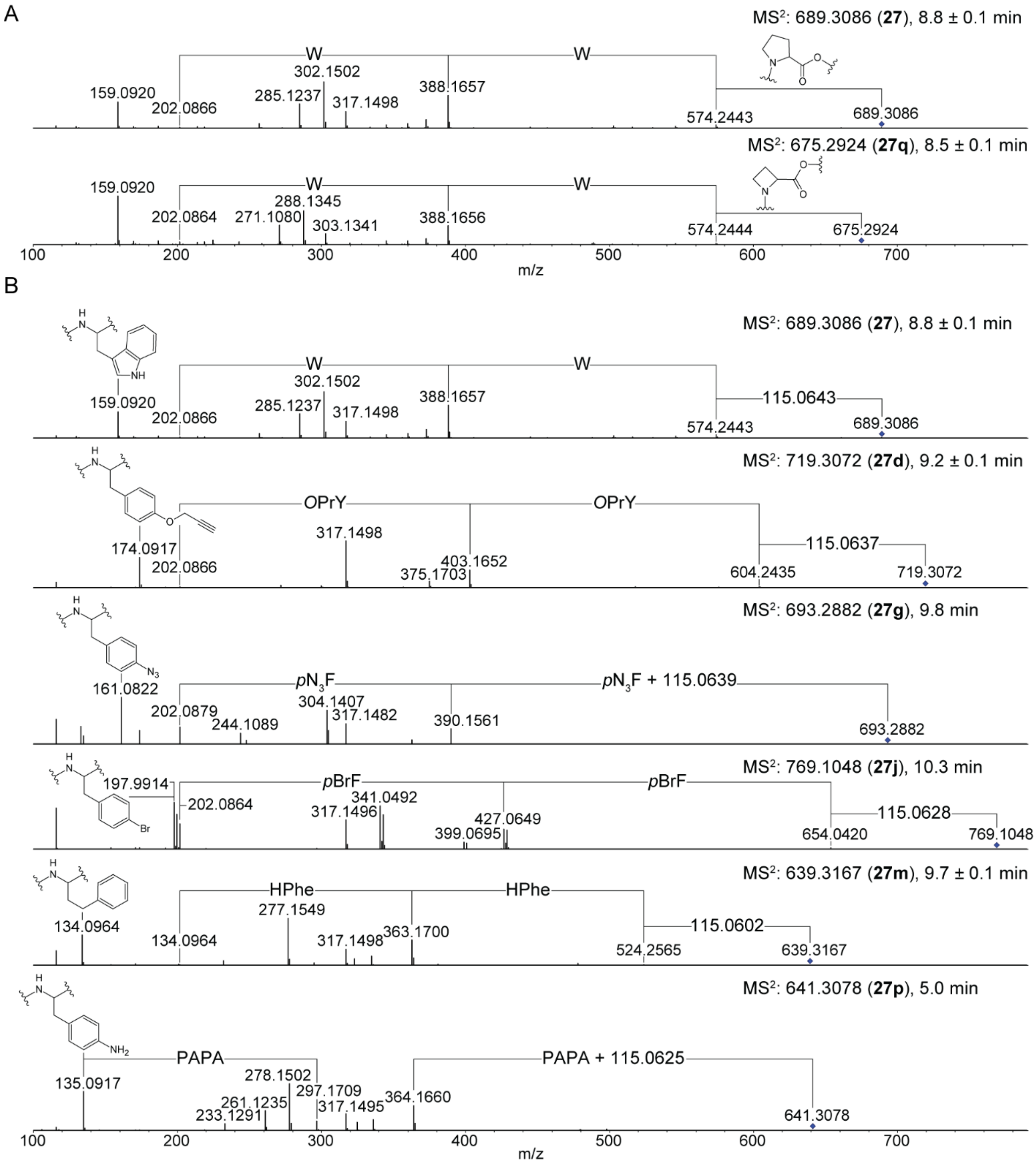

**Figure S15.** $MS^2$ fragmentation spectra of xeneprotide derivatives **27d**, **27g**, **27j**, **27m**, **27p**, and **27q** depicted in Figure S13 produced by precursor-directed biosynthesis compared to natural xeneprotide (**27**).

**A**

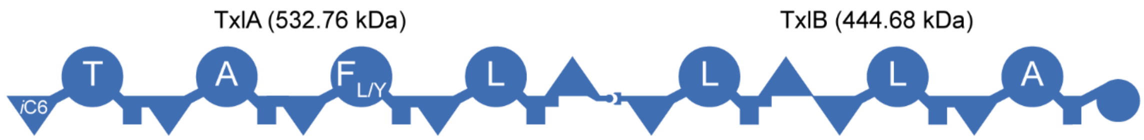

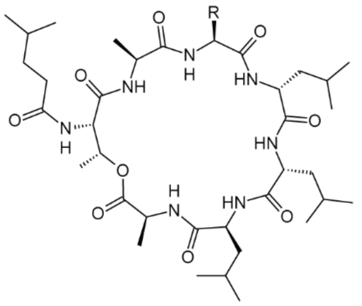

taxIllaid E (**28**)

| AA | # | rel. incorporation |
|---|---|---|
| Tyr | **28** | |
| $p$N$_3$F | **28b** | 11% |
| PAPA | **28c** | 92% |
| 2-Fua | **28d** | 2% |

**B**

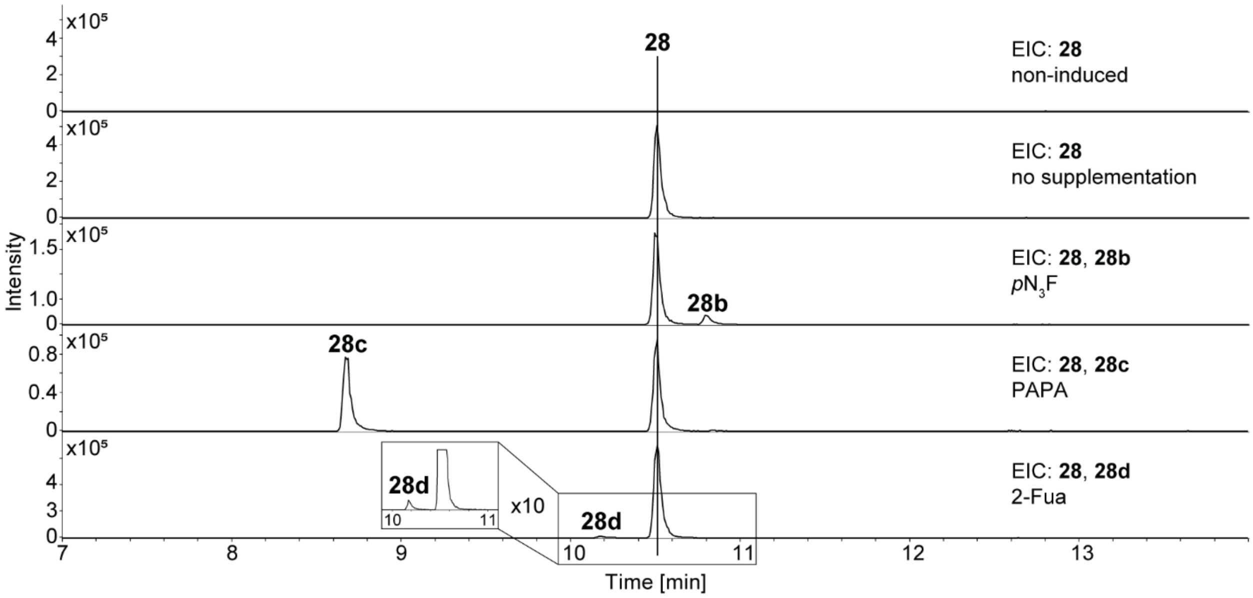

**Figure S16.** Precursor-directed biosynthesis of taxIllaid derivatives. **A.** Schematic depiction of the NRPS TxlAB from *X. indica* DSM 17382 and produced taxIllaid derivatives **28-28d** identified by supplementing the production medium with indicated non-cognate amino acids. A-domain specificity indicated by the amino acid one-letter code. **B.** Extracted ion chromatograms (EIC) of the taxIllaid derivatives **28-28d**.

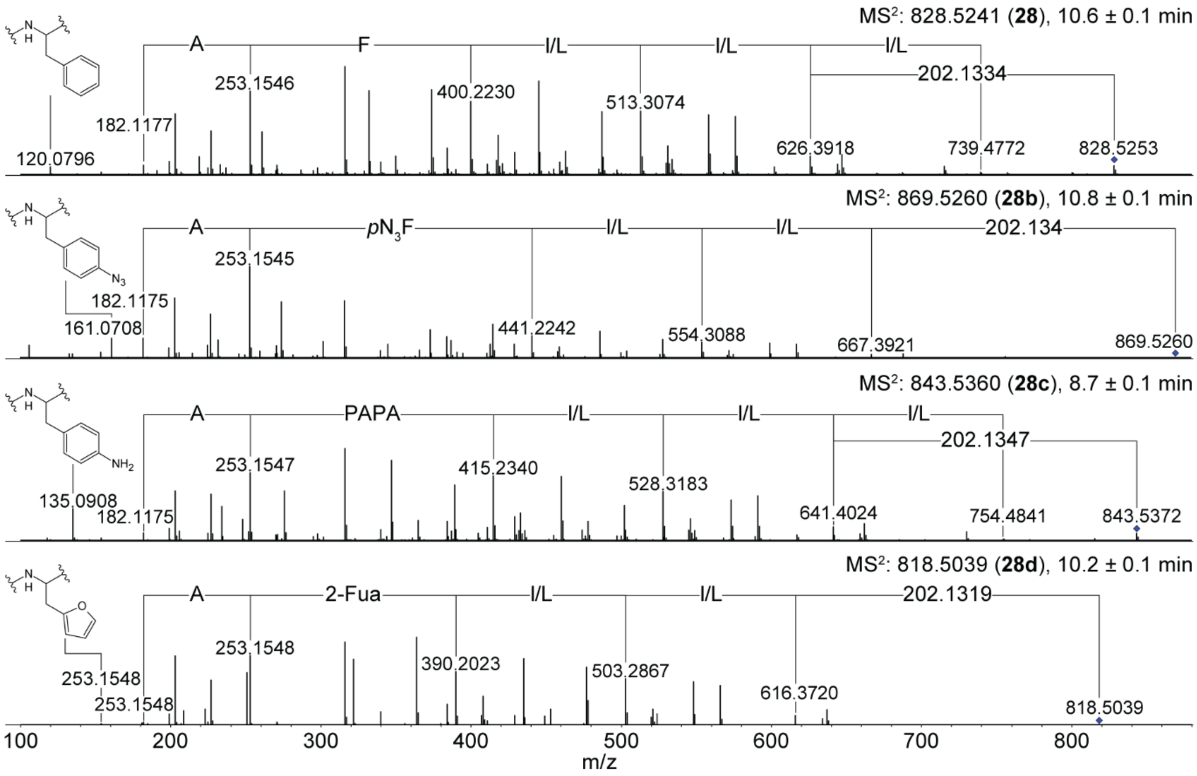

**Figure S17.** $MS^2$ fragmentation spectra of taxlllaid derivatives **28b-28d** depicted in Figure S16 produced by precursor-directed biosynthesis compared to natural taxlllaid E (**28**).

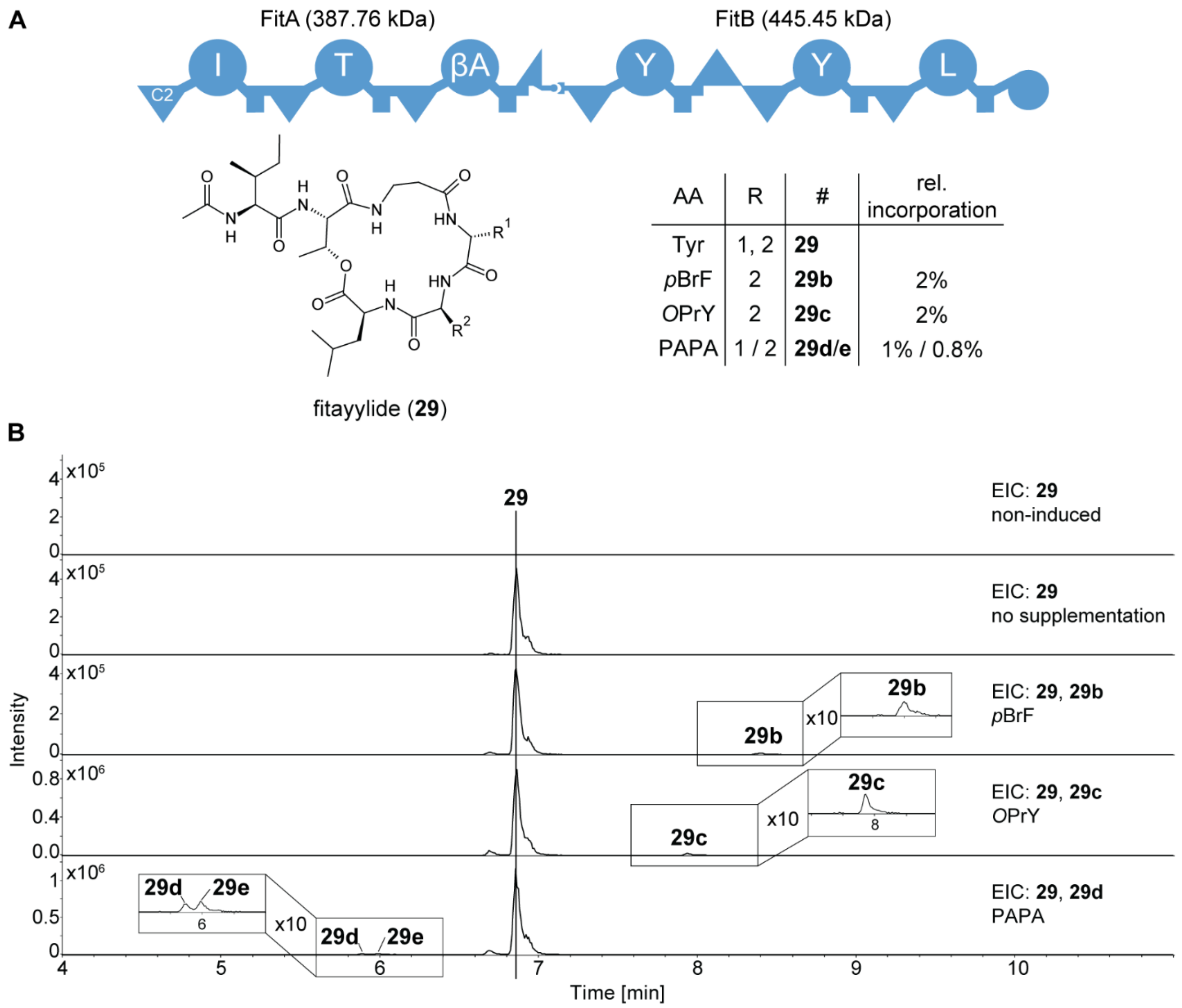

**Figure S18.** Precursor-directed biosynthesis of fitayylide derivatives. **A.** Schematic depiction of the NRPS FitAB from *X. innexi* DSM 16336 and produced fitayylide derivatives **29-29f** identified by supplementing the production medium with indicated non-cognate amino acids. Expression of *fitAB* was done in *E. coli* DH10B::*mtaA*. A-domain specificity indicated by the amino acid one-letter code. βA: beta-alanine. **B.** Extracted ion chromatograms (EIC) of the fitayylide derivatives **29-29f**.

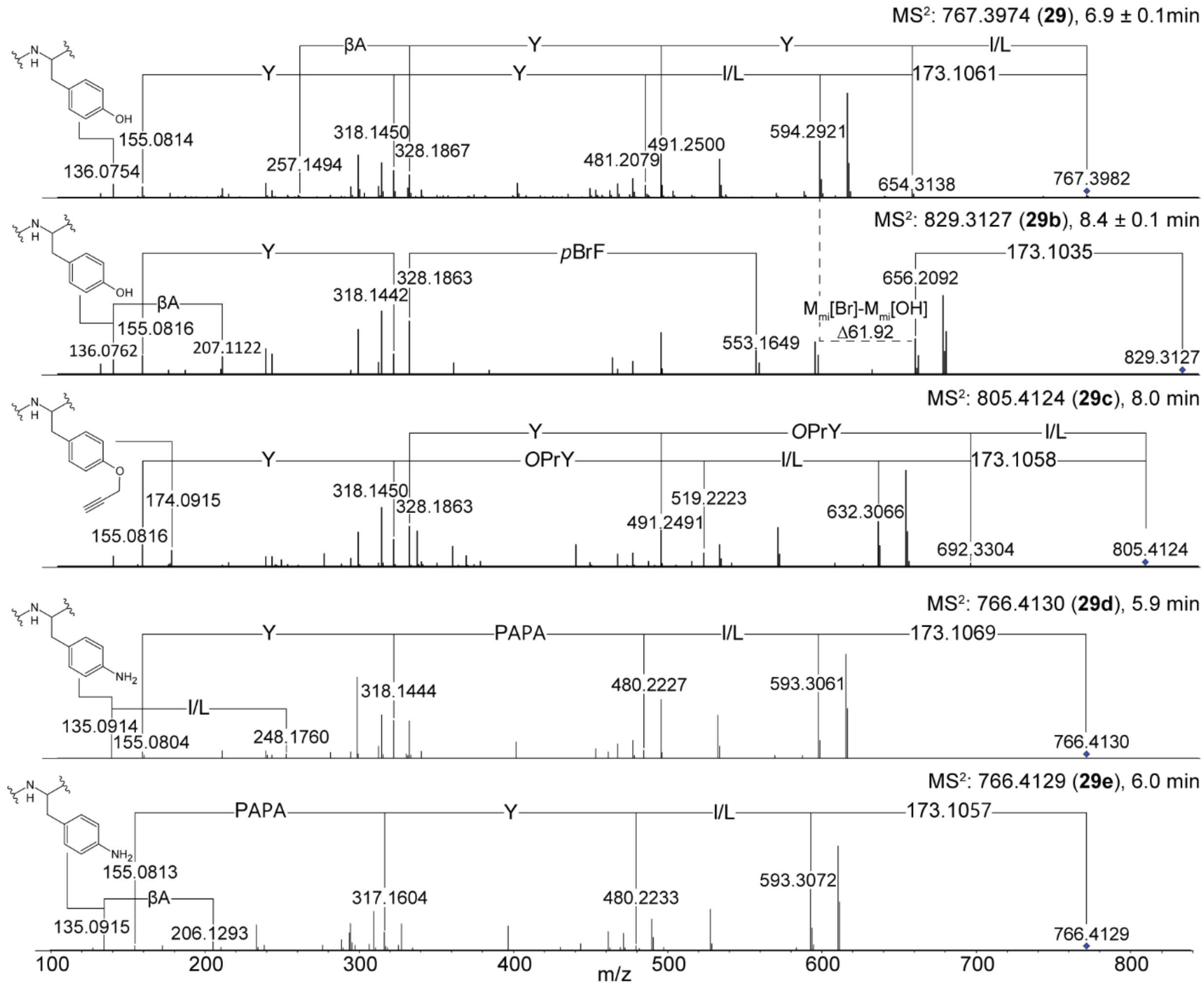

**Figure S19.** $MS^2$ fragmentation spectra of fitayylide derivatives **29b**-**29e** depicted in Figure S18 produced by precursor-directed biosynthesis compared to natural fitayylide (**29**).

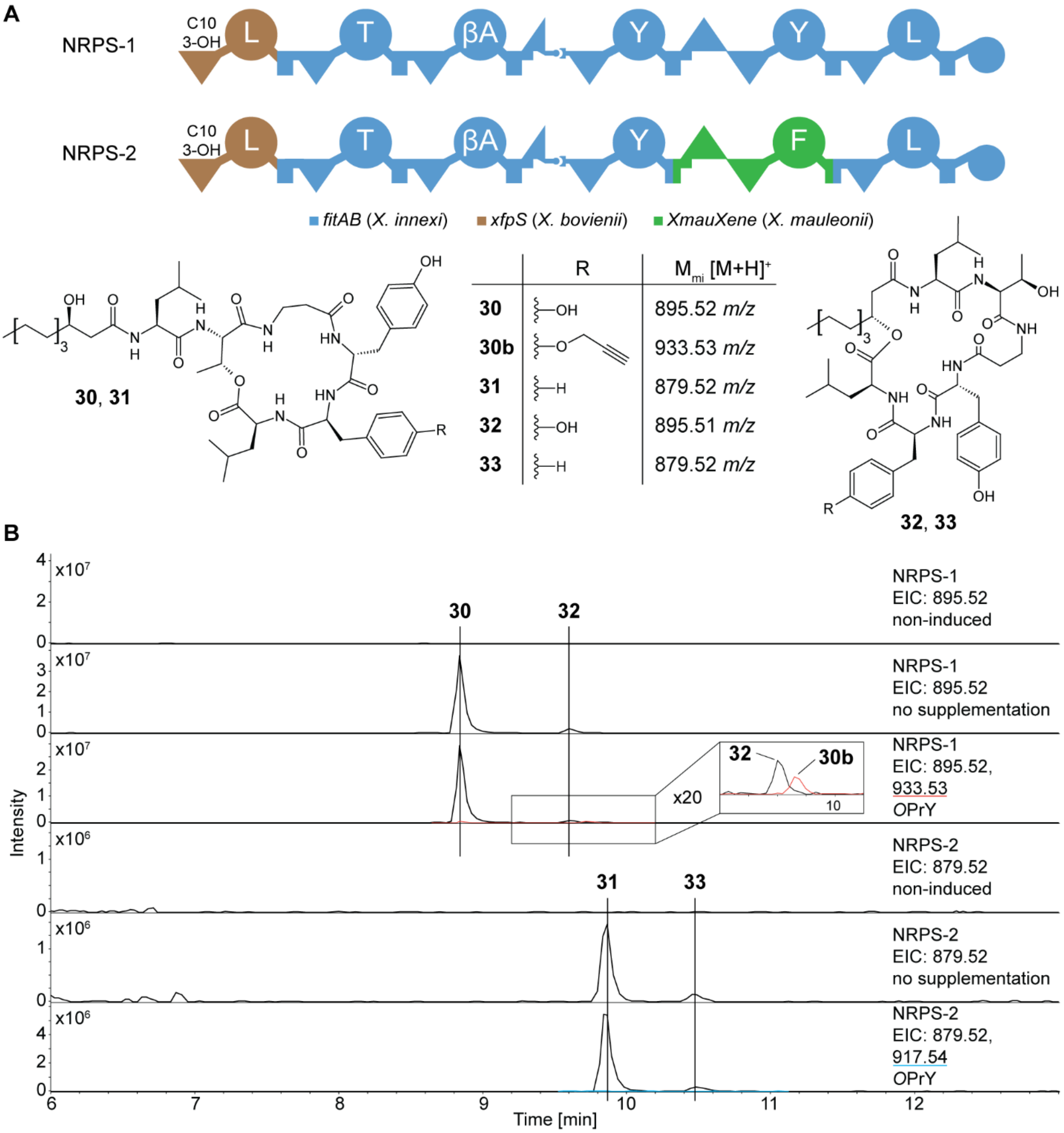

**Figure S20.** Identification of the promiscuous tyrosine-specific A-domain. **A.** Scheme and products of synthetic hybrid NRPS-1 and NRPS-2 used to identify, which tyrosine-specific A-domain of FitB is promiscuous. **B.** Extracted ion chromatograms (EIC) of products **30-33** produced by heterologous expression of *NRPS-1* and *NRPS-2* subjected to precursor-directed biosynthesis in *E. coli* DH10B::*mtaA* supplementing *O*PrY (**13**) to the production medium.

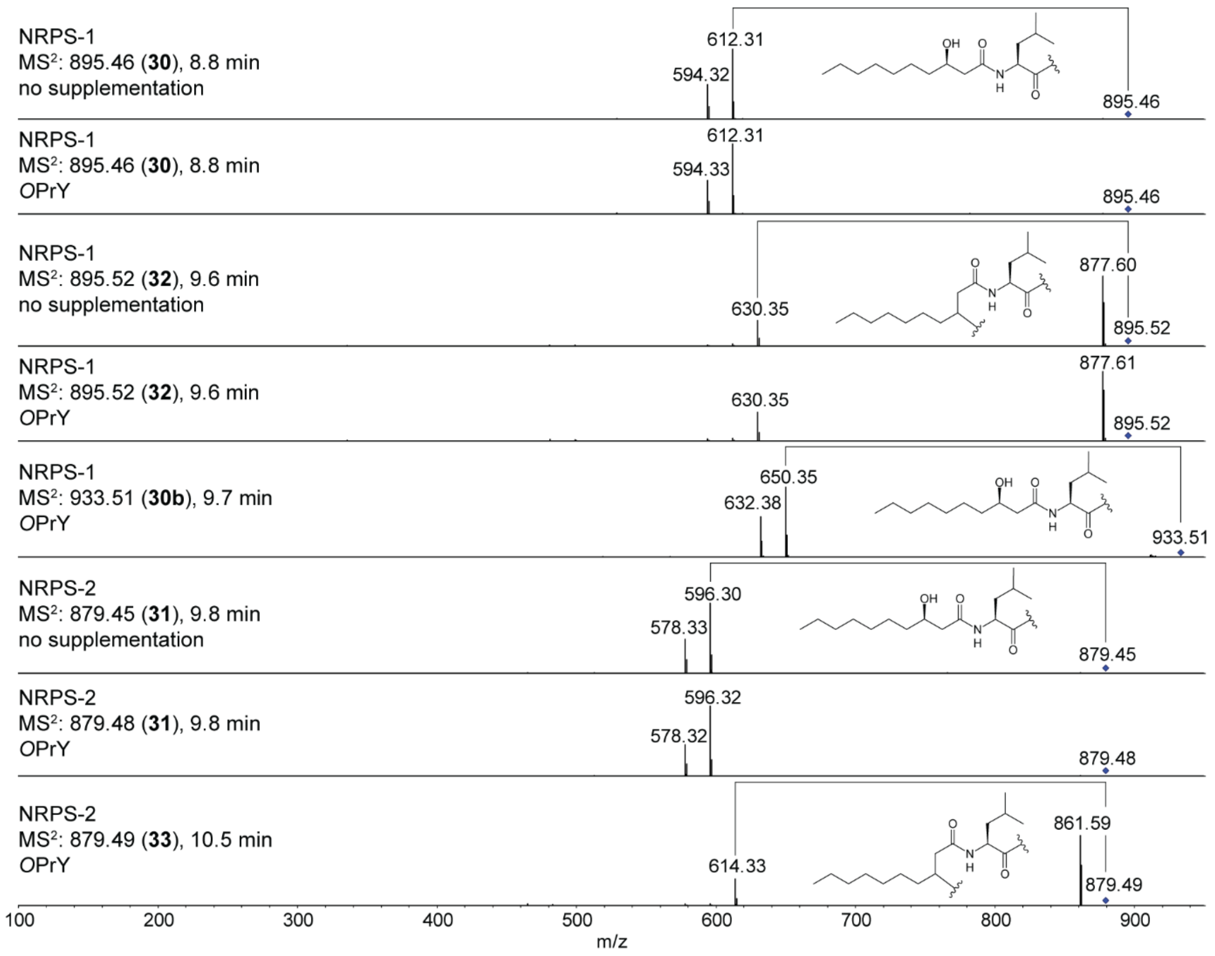

**Figure S21.** $MS^2$ fragmentation spectra of the products **30**, **30b**, and **31**-**33** depicted in Figure S20 produced by precursor-directed biosynthesis with NRPS-1 and NRPS-2. For **33** produced by NRPS-2 without supplementation, no $MS^2$ spectra could be obtained.

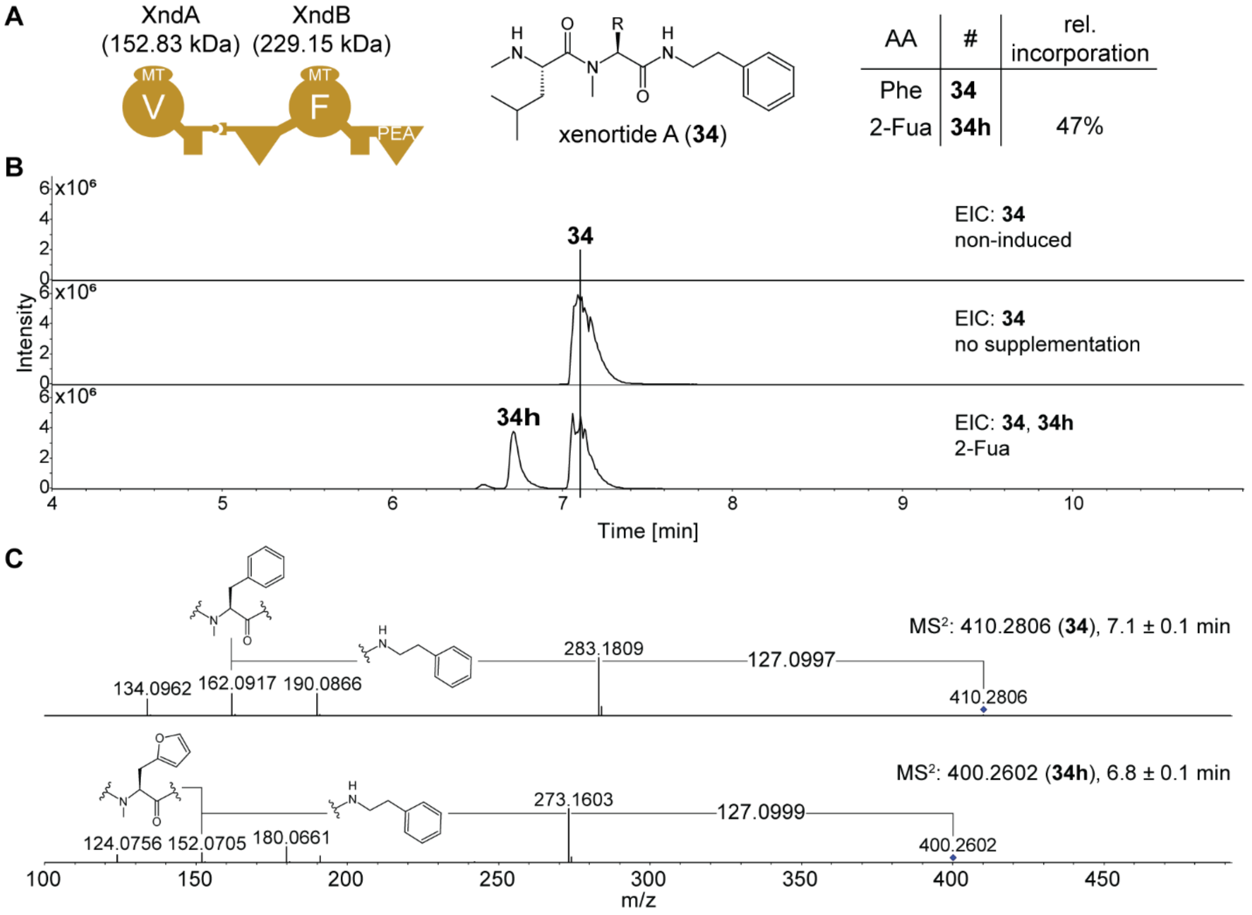

**Figure S22.** Precursor-directed biosynthesis of xenortide derivatives. **A.** Schematic depiction of the NRPS XndAB from *X. nematophila* ATCC 19061 and produced xenortide A (**34**) and its derivative **34h** identified by supplementing the production medium with 2-Fua (**7**). A-domain specificity indicated by the amino acid one-letter code. **B.** Extracted ion chromatograms (EIC) of the xenortide A (**34**) and its derivative **34h**. **C.** MS$^2$ fragmentation spectra of **34** and **34h**.

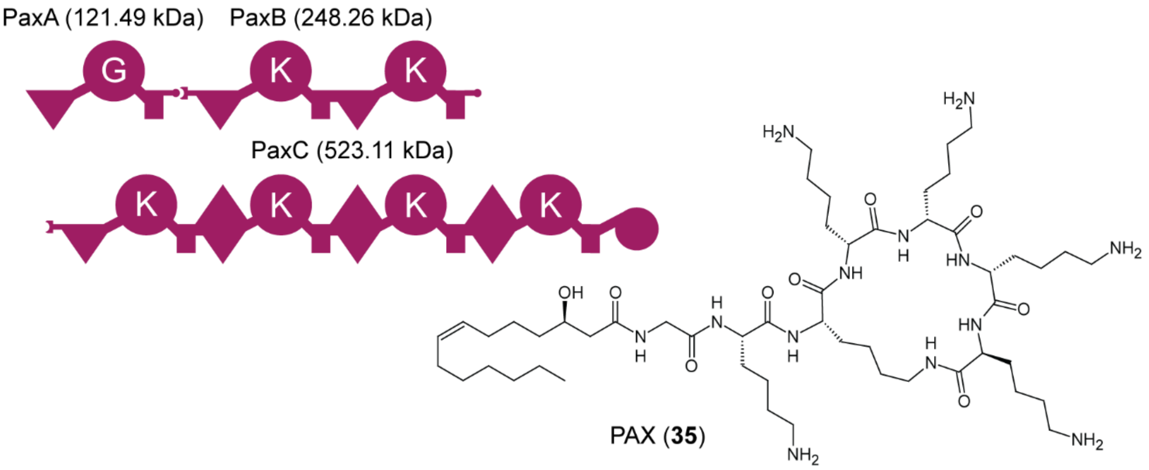

**Figure S23.** Schematic depiction of the NRPS PaxABC *X. doucetiae* DSM 17909 and structure of produced PAX (**33**). A-domain specificity indicated by the amino acid one-letter code.

**A**

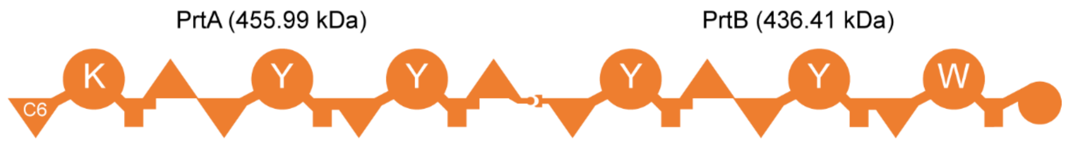

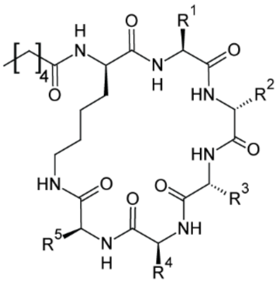

protegomycin (**36**)

| AA | R | # | rel. incorporation | AA | R | # |
|---|---|---|---|---|---|---|
| 4x Tyr, Trp | 1 - 4, 5 | **36** | | | | |
| *O*PrY | 5 | **36b** | 15% | | | |
| *O*PrY | 1 / 2 / 3 / 4 | **36c** | 23% | 2x PAPA | 1 / 2 / 3 / 4 , 5 | **36f** |
| PAPA | 5 | **36d** | 56% | 2x PAPA | 1 / 2 / 3 / 4 | **36g** |
| PAPA | 1 / 2 / 3 / 4 | **36e** | 328% | 3x PAPA | 1 / 2 / 3 / 4 | **36h** |
| DOPA | 1 / 2 / 3 / 4 | **36i** | 40% | 2x DOPA | 1 / 2 / 3 / 4 | **36j** |

**B**

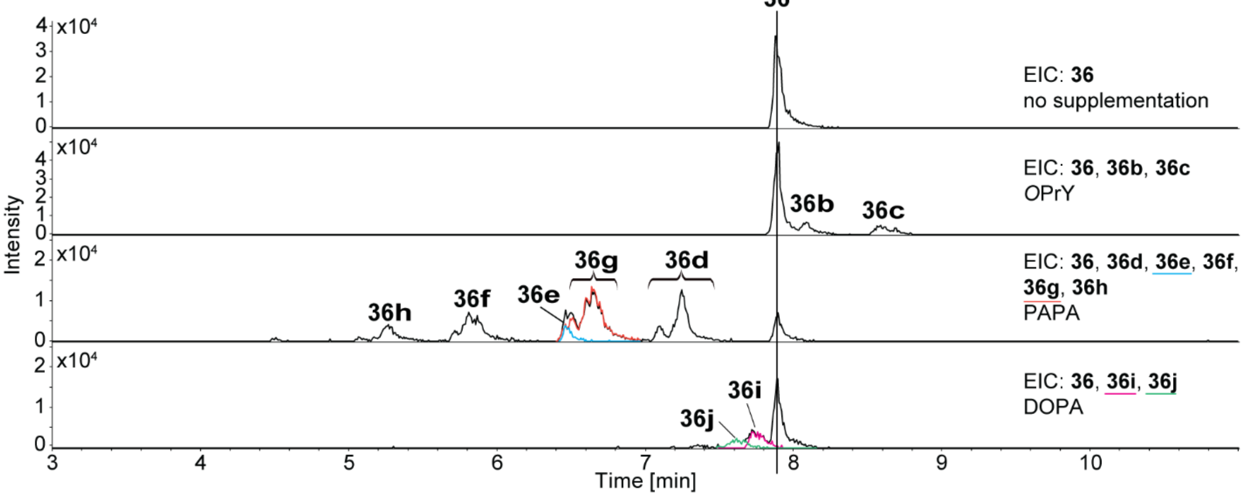

**Figure S24.** Precursor-directed biosynthesis of protegomycin derivatives. **A.** Schematic depiction of the NRPS PrtAB from *X. doucetiae* DSM 17909 and produced protegomycin derivatives **36**-**36j** identified by supplementing the production medium with indicated non-cognate amino acids. A-domain specificity indicated by the amino acid one-letter code. For structures of **36c** and **36e**-**36j** no clear exchange position could be assigned. **B.** Extracted ion chromatograms (EIC) of the protegomycin derivatives **36**-**36j**.

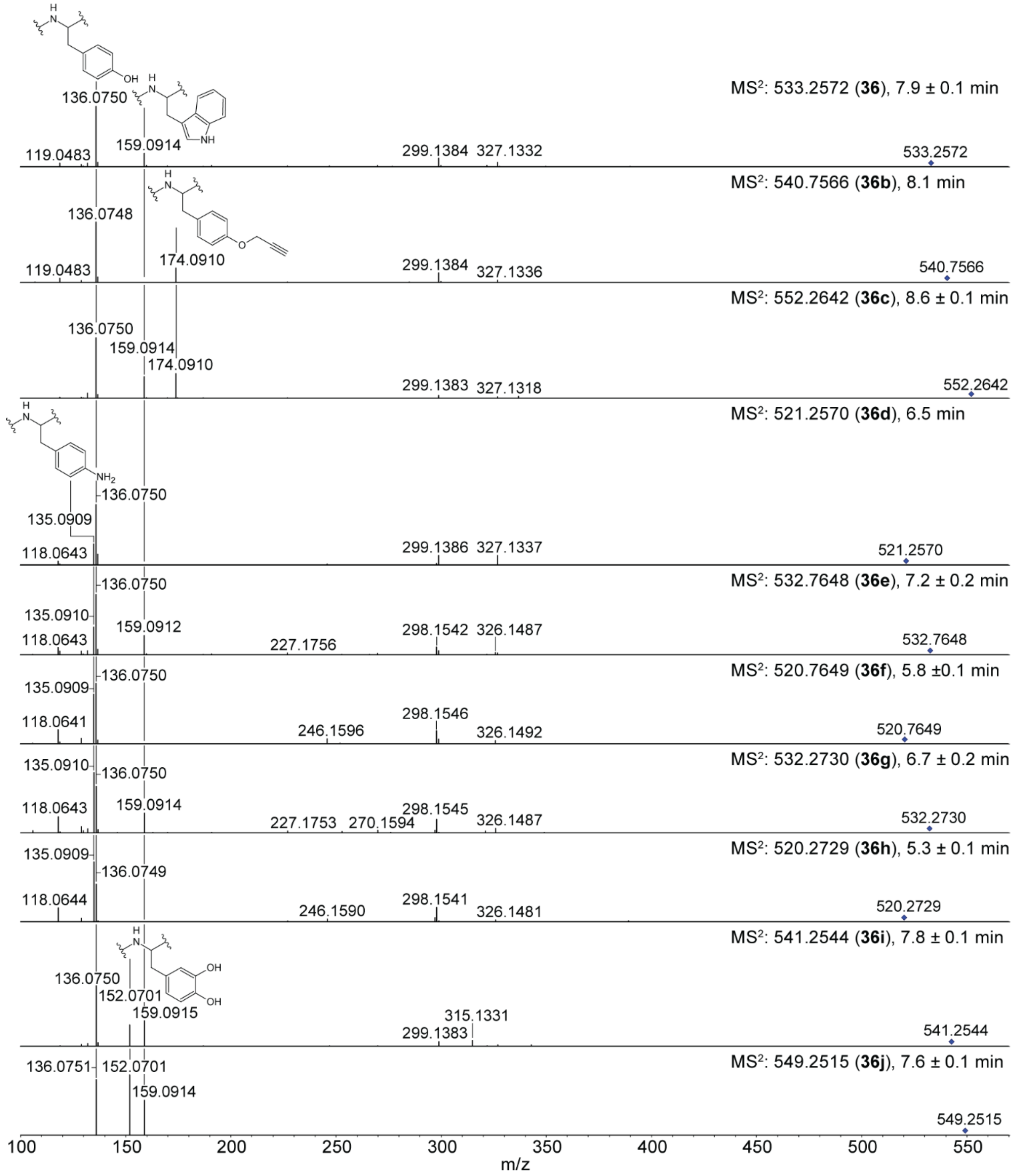

**Figure S25.** $MS^2$ fragmentation spectra of protegomycin derivatives **36b-36j** depicted in Figure S24 produced by precursor-directed biosynthesis compared to natural protegomycin (**36**).

#### NRPS Engineering with Promiscuous A-Domains

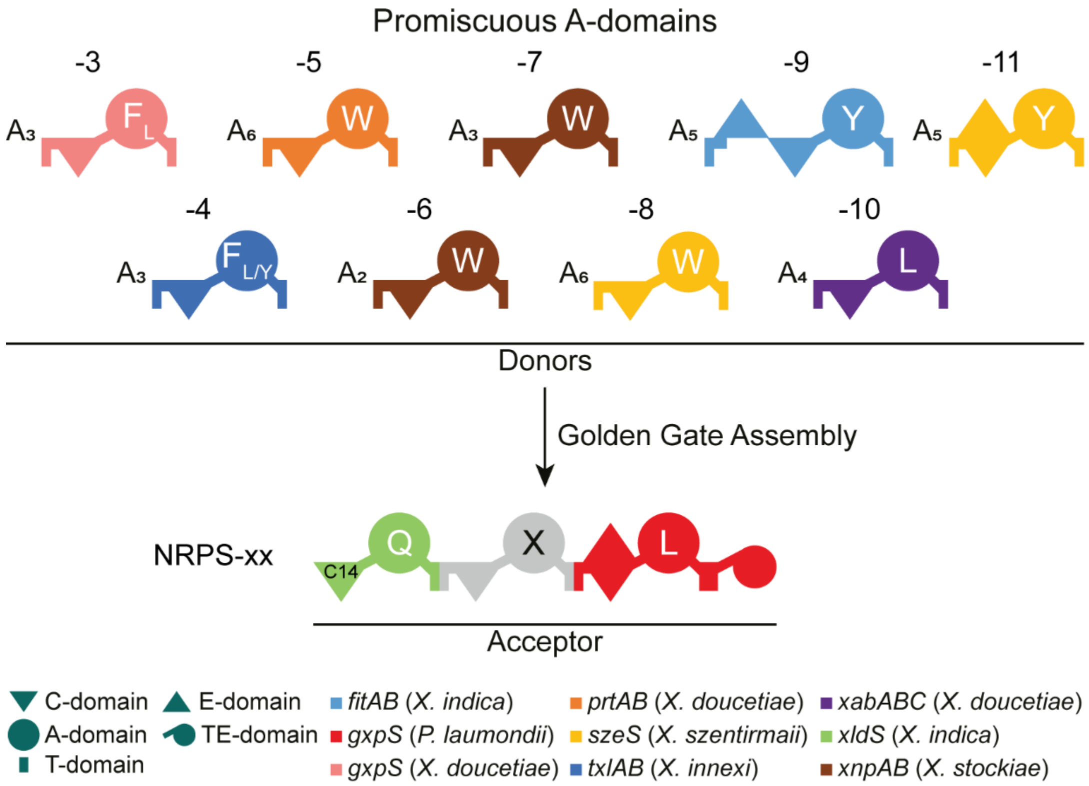

**Figure S26.** Overview of the selected promiscuous A-domains for Golden Gate-based NRPS engineering[57,58]. Natural A-domain specificity indicated by the amino acid one-letter code. Ten different XUT$^{IV}$ modules[32] from eight different NRPS are used as donor library. Golden Gate assembly of the donors with the acceptor results in the synthetic hybrids NRPS-3 to NRPS-11.

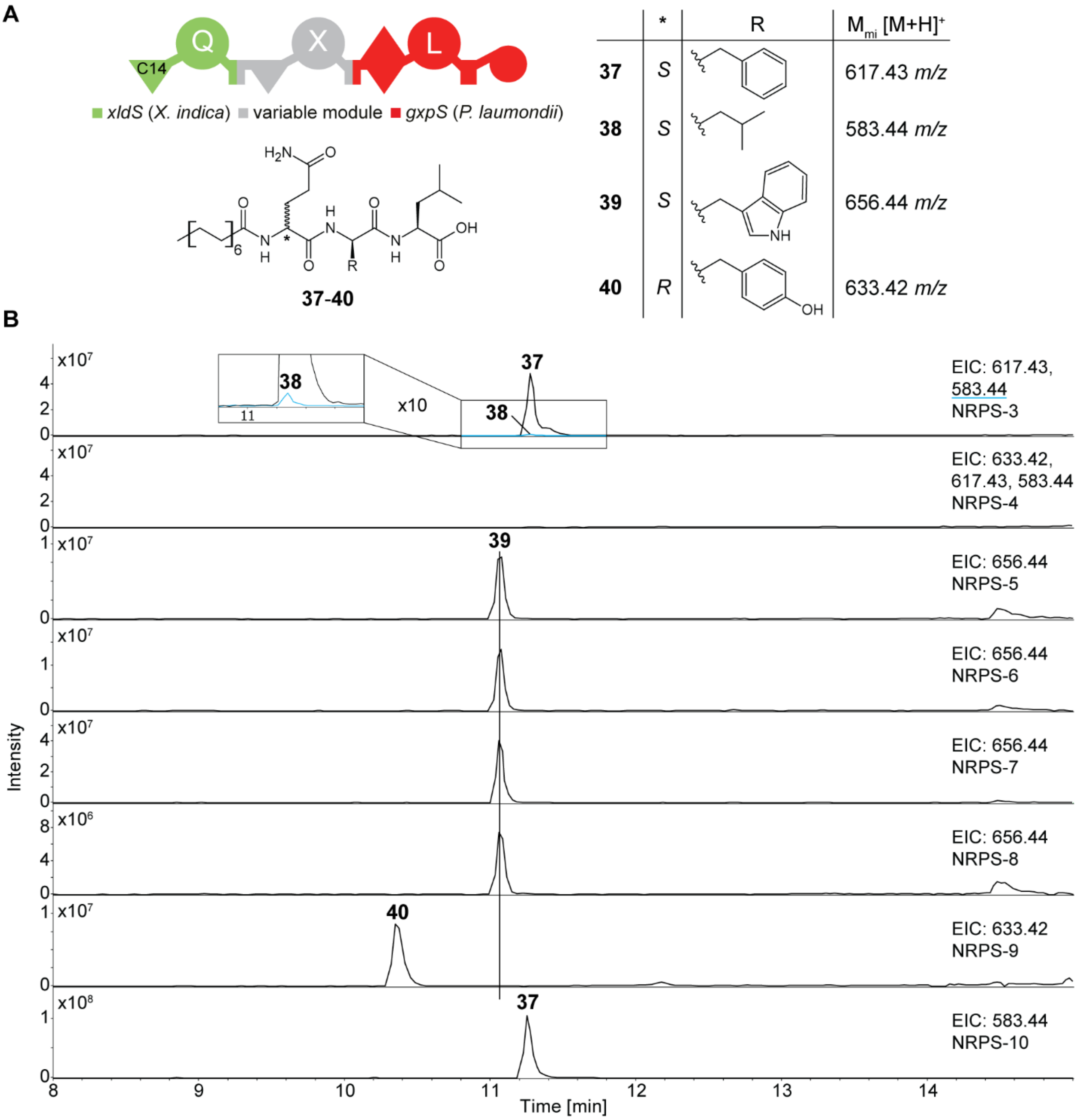

**Figure S27.** Production of synthetic hybrids *NRPS-3* to *NRPS-10* heterologously expressed in *E. coli* DH10B::*mtaA*. **A.** Schematic depiction of hybrid NRPS and the structures of produced lipopeptides **37-40**. Grey module highlights variable position within the NRPS corresponding to Figure S26. A-domain specificity indicated by the amino acid one-letter code. **B.** Extracted ion chromatograms (EIC) of the lipopeptides **37-40**.

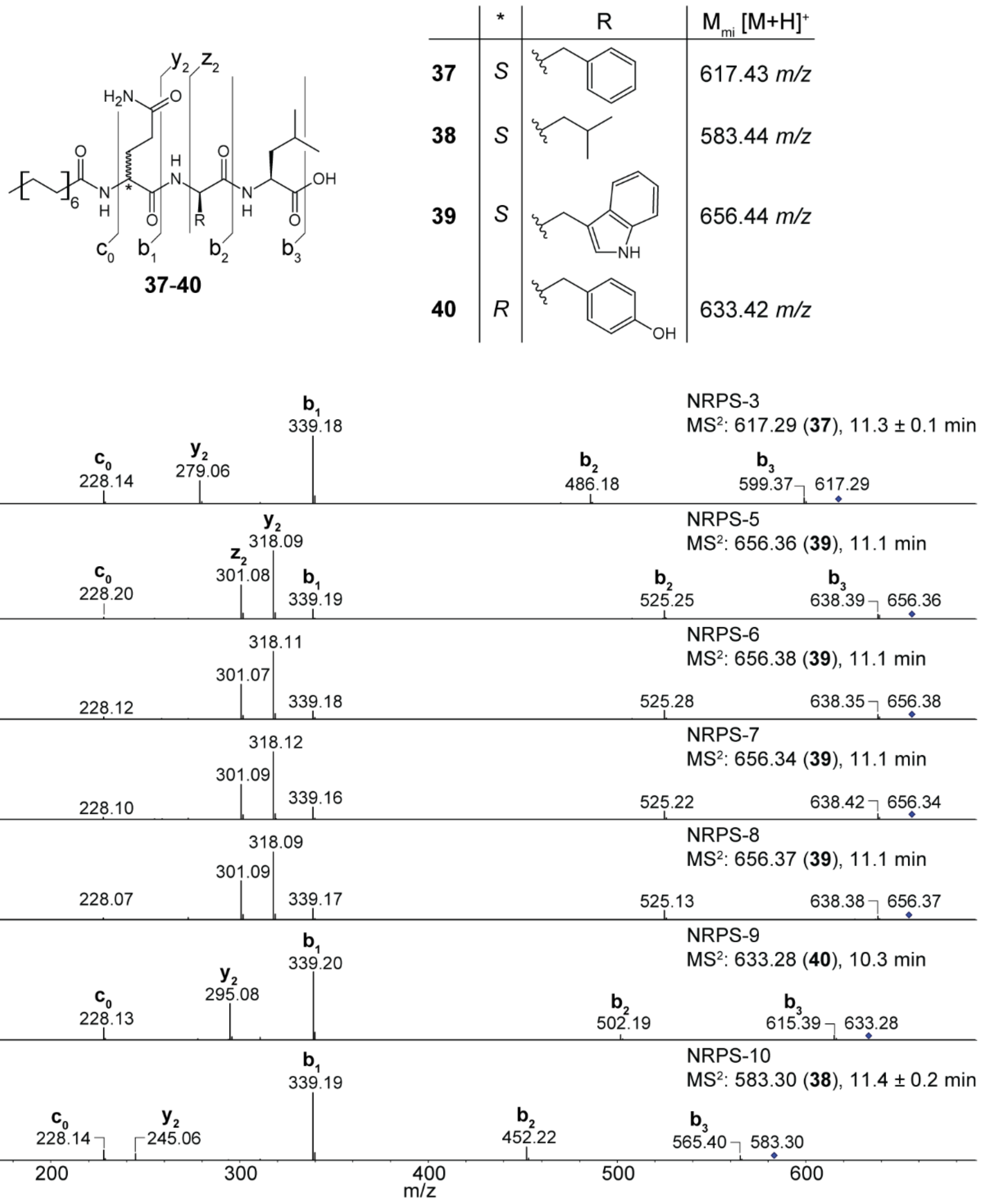

**Figure S28.** $MS^2$ fragmentation spectra corresponding to Figure S27 of the lipopeptides **37**-**40** produced by synthetic hybrid NRPS-3 and NRPS-5 to NRPS-10.

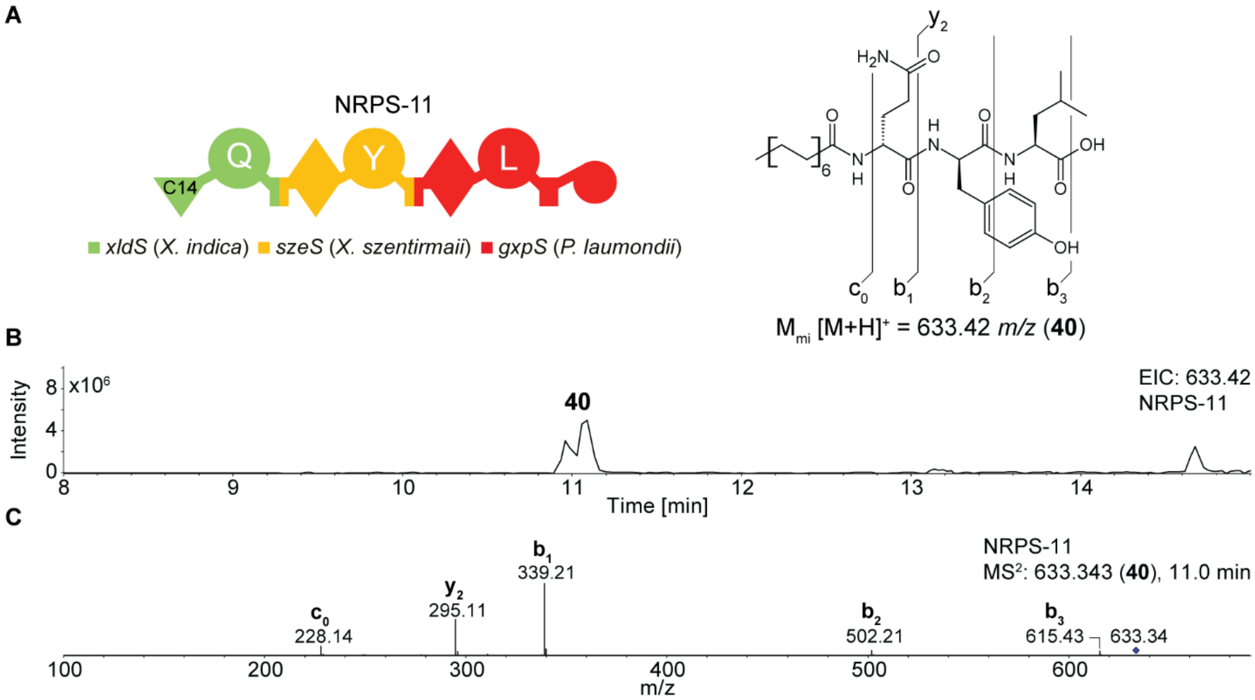

**Figure S29.** Function of NRPS-11. **A.** Schematic depiction of synthetic hybrid NRPS-11 and structure of produced lipopeptide **40**. A-domain specificity indicated by the amino acid one-letter code. **B.** Extracted ion chromatogram (EIC) of the lipopeptide **40**. **C.** $MS^2$ fragmentation spectrum of **40**. Data and NRPS-11 taken from Podolski *et al.* **2025**[58].

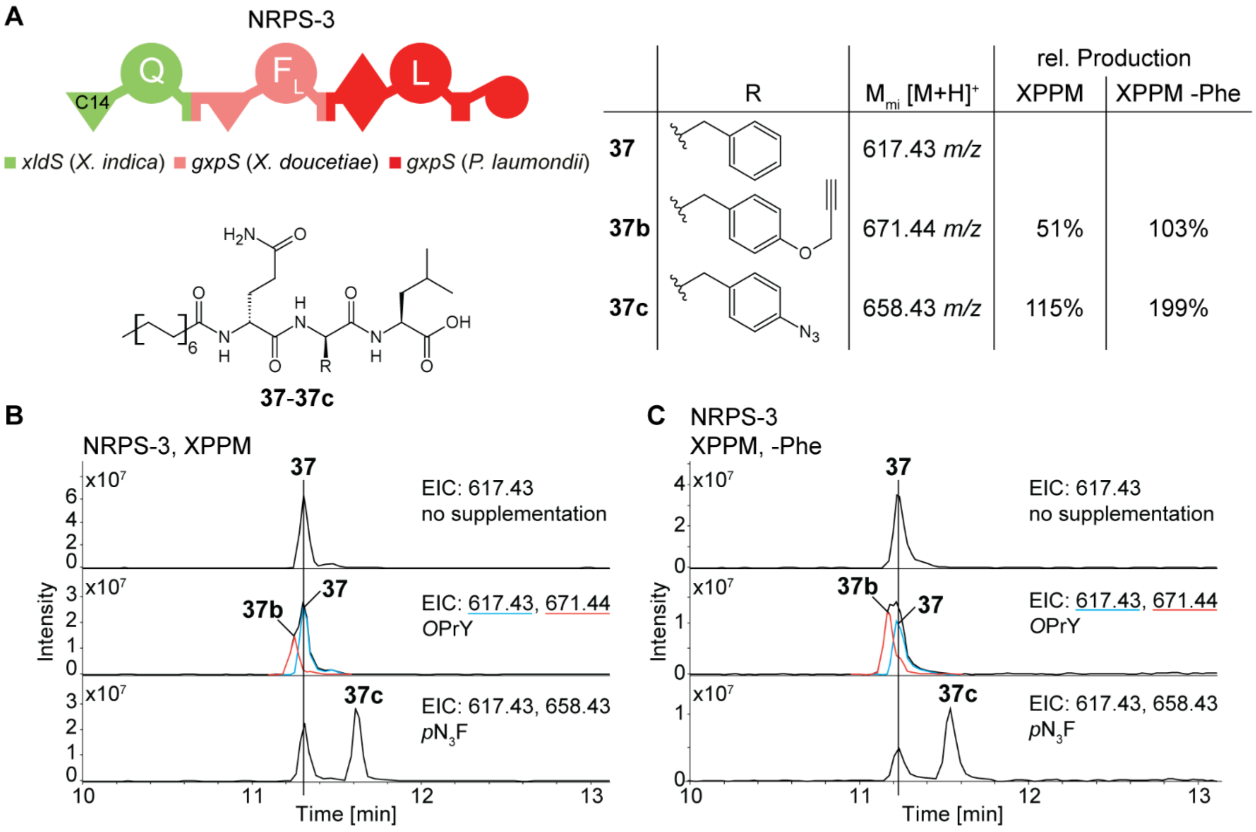

**Figure S30.** Precursor-directed biosynthesis of lipopeptides produced by heterologous expression of *NRPS-11* in *E. coli* DH10B::*mtaA*. **A.** Schematic depiction of synthetic hybrid NRPS-11 and structures of produced lipopeptides **37**, **37b**, and **37c** together with their relative production compared to **37** in different media. A-domain specificity indicated by the amino acid one-letter code. **B.** Extracted ion chromatograms (EIC) of the lipopeptides **37-37c** produced in XPPM supplemented with **12** ($pN_3F$) or **13** (*O*PrY). **C.** EIC of **37-37c** produced in XPPM without phenylalanine (-Phe) supplemented with **12** or **13**.

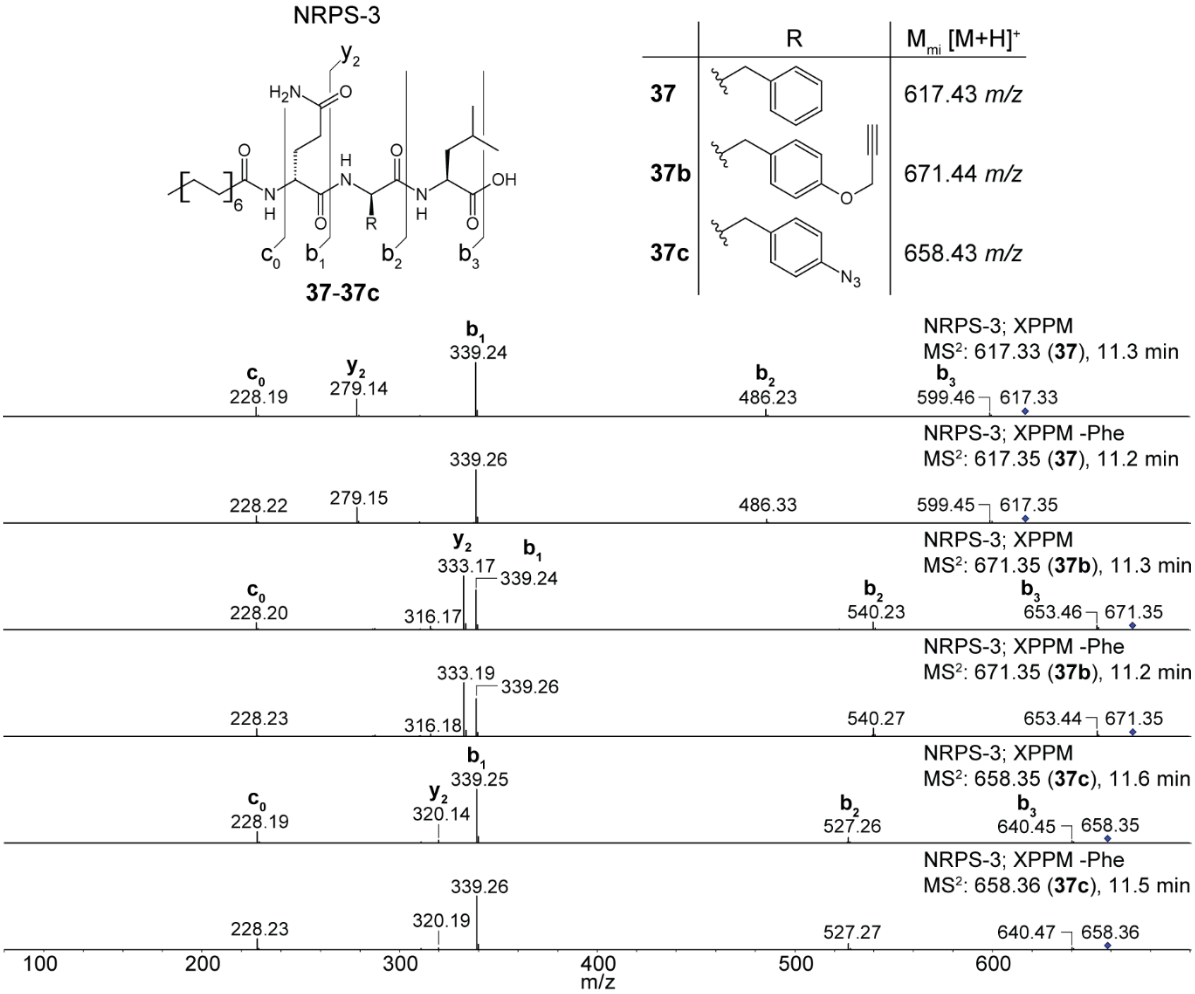

**Figure S31.** $MS^2$ fragmentation spectra corresponding to Figure S30 of lipopeptides **37b** and **37c** produced by precursor-directed biosynthesis using NRPS-3 compared to **37**.

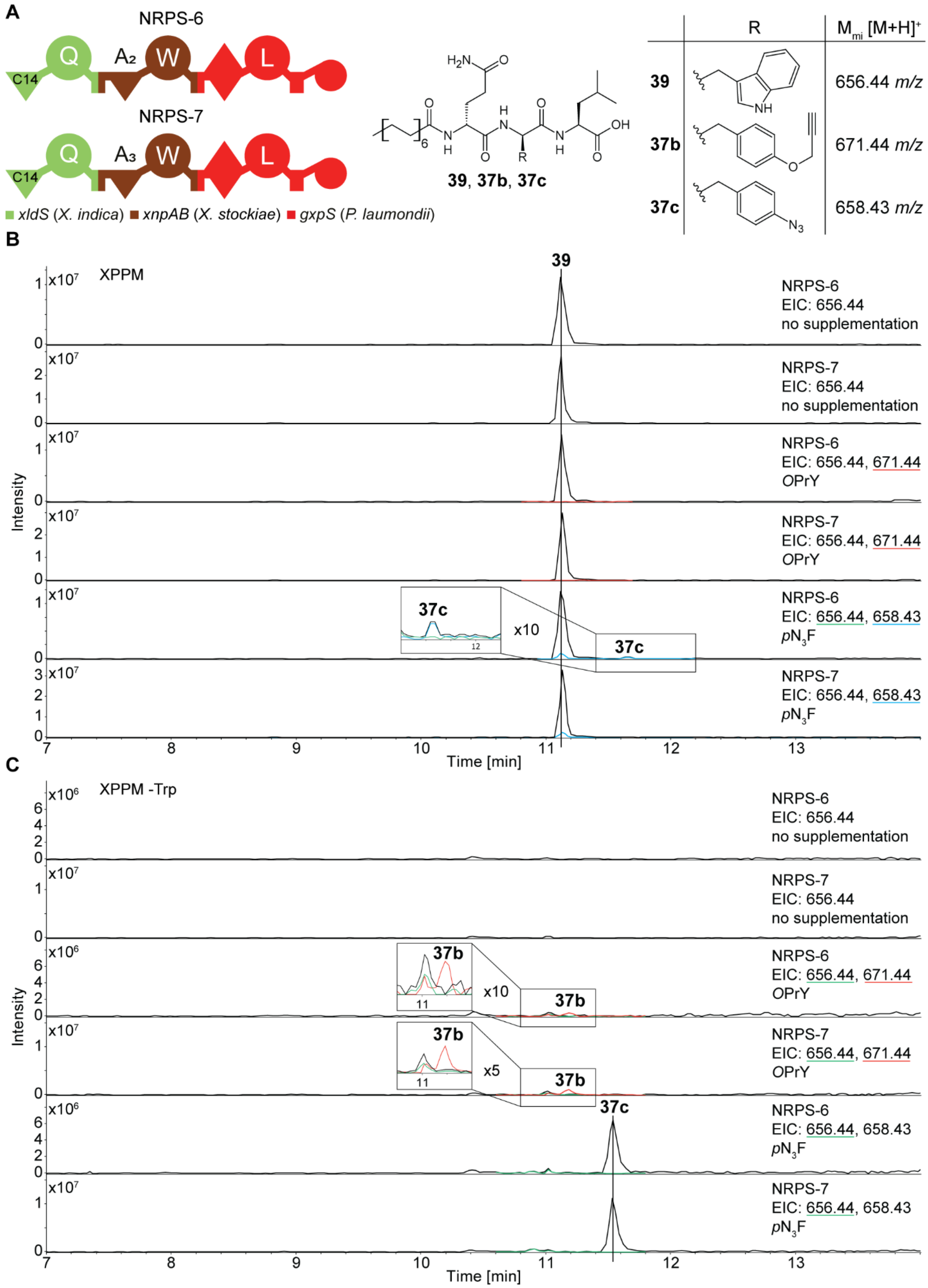

**Figure S32.** Precursor-directed biosynthesis of lipopeptides produced by heterologous expression of *NRPS-6* and *NRPS-7* in *E. coli* DH10B::*mtaA*. **A.** Schematic depiction of synthetic hybrids NRPS-6 and NRPS-7 together with the structures of produced lipopeptides **39**, **37b**, and

**37c**. A-domain specificity indicated by the amino acid one-letter code. **B.** Extracted ion chromatograms (EIC) of the lipopeptides **39**, **37b**, and **37c** produced in XPPM supplemented with **12** ($pN_3F$) or **13** (*O*PrY). **C.** EIC of **39**, **37b**, and **37c** produced in XPPM without tryptophan (-Trp) supplemented with **12** or **13**.

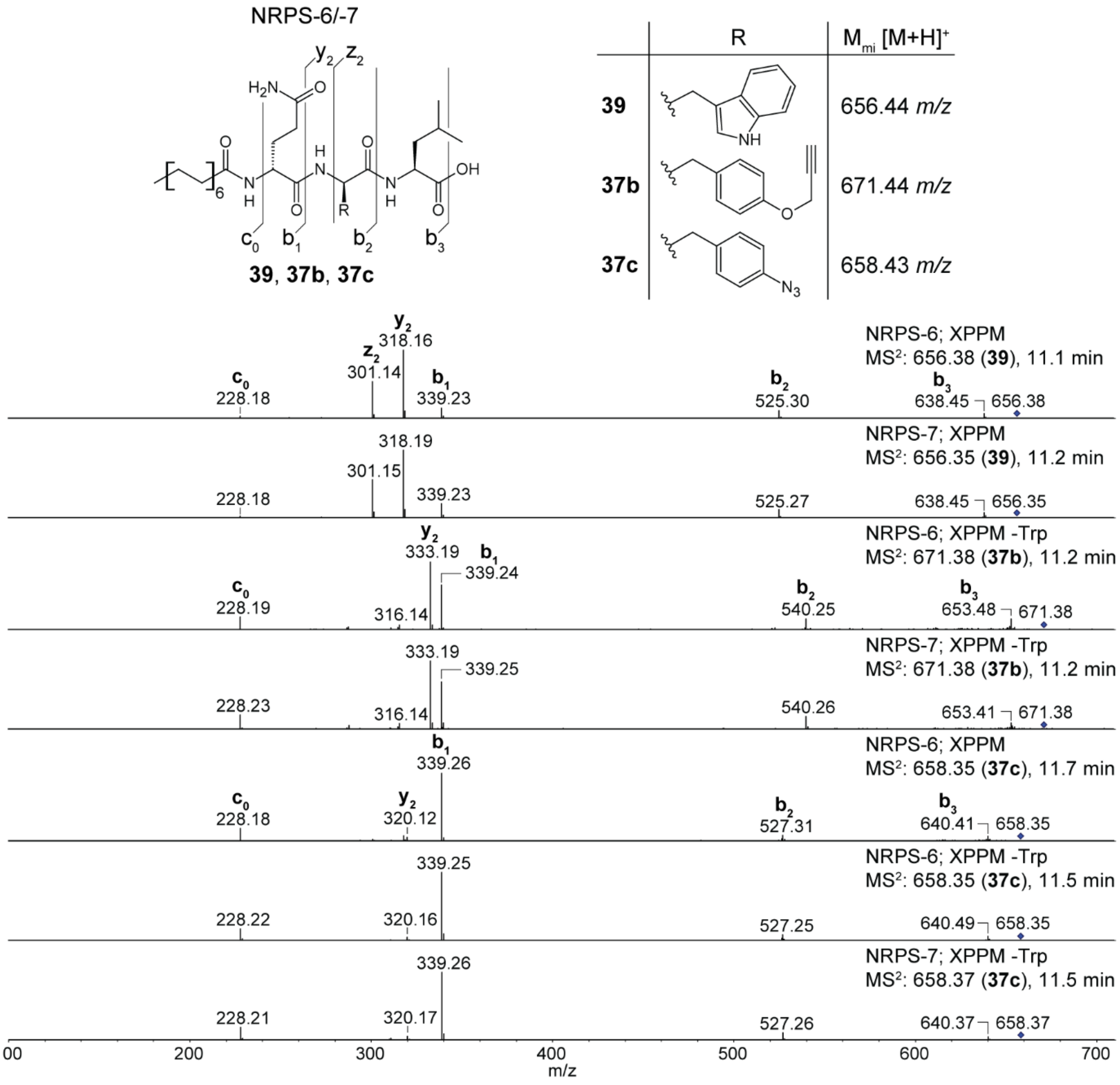

**Figure S33.** $MS^2$ fragmentation spectra corresponding to Figure S32 of lipopeptides **35b** and **35c** produced by precursor-directed biosynthesis using NRPS-6 and NRPS-7 compared to **37**.

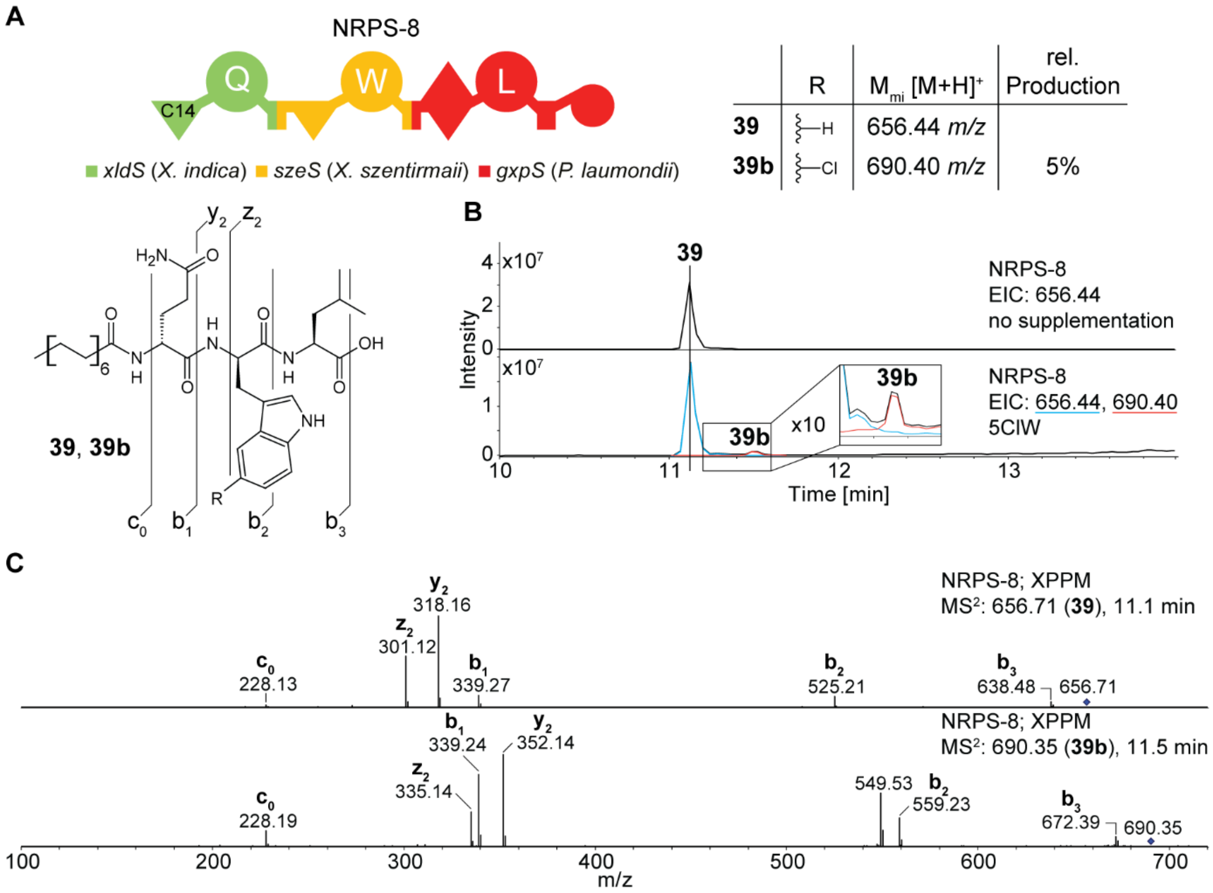

**Figure S34.** Precursor-directed biosynthesis of lipopeptides produced by heterologous expression of *NRPS-8* in *E. coli* DH10B::*mtaA*. **A.** Schematic depiction of synthetic hybrid NRPS-8 and structures of produced lipopeptides **39** and **39b** together with their relative production. A-domain specificity indicated by the amino acid one-letter code. **B.** Extracted ion chromatograms (EIC) of the lipopeptides **39** and **379b** produced in XPPM supplemented with **2** (5ClW). **C.** $MS^2$ fragmentation spectra of **39b** produced by NRPS-16 compared to **39**.

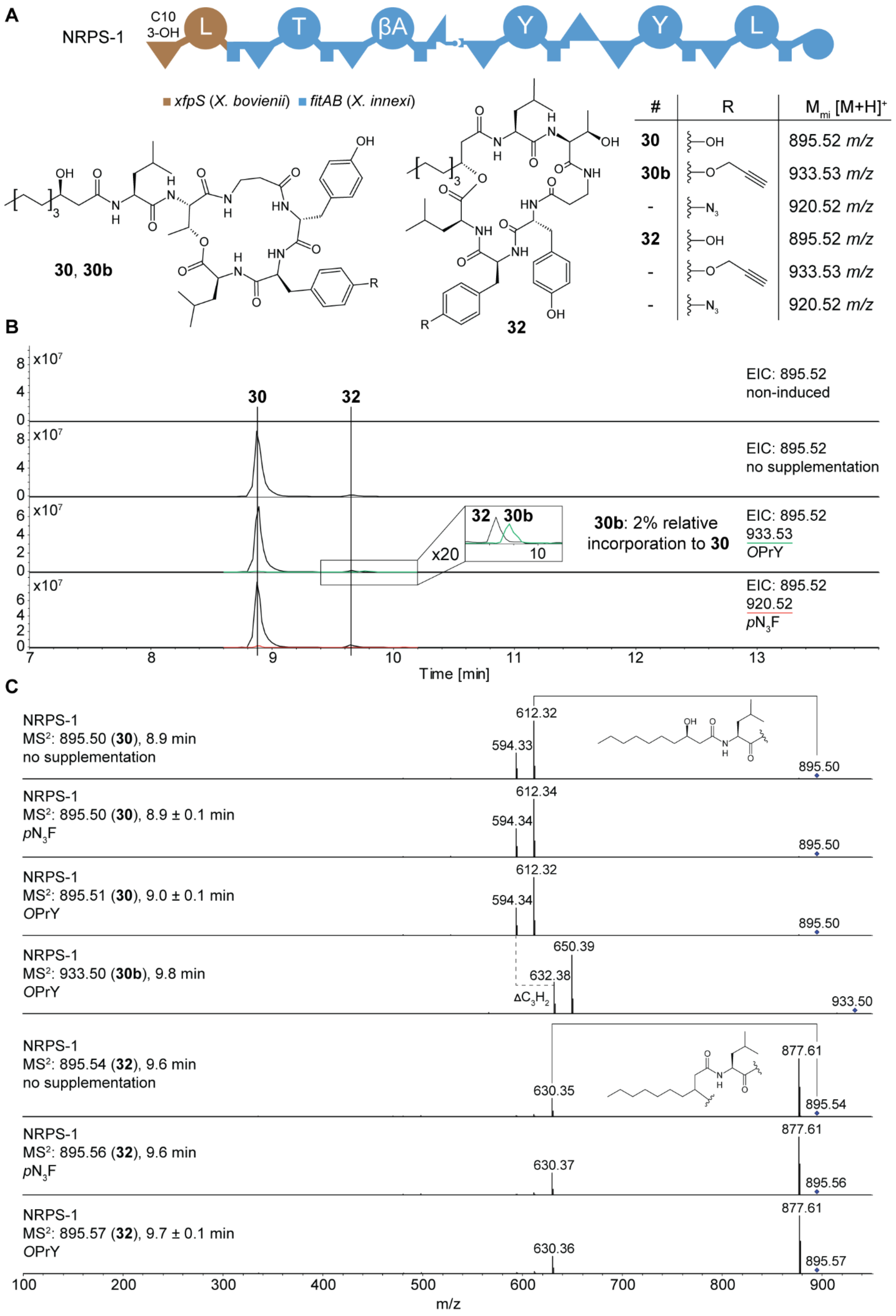
A
NRPS-1
C10
3-OH
L
T
βA
Y
Y
L
xfpS (X. bovienii) fitAB (X. innexi)
30, 30b
32
#
R
M<sub>mi</sub> [M+H]+
30 OH 895.52 m/z
30b 933.53 m/z
- 920.52 m/z
32 OH 895.52 m/z
- 933.53 m/z
- 920.52 m/z
B
EIC: 895.52
non-induced
EIC: 895.52
no supplementation
30b: 2% relative
incorporation to 30
EIC: 895.52
933.53
OPrY
EIC: 895.52
920.52
pN3F
x20
Time [min]
C
NRPS-1
MS2: 895.50 (30), 8.9 min
no supplementation
NRPS-1
MS2: 895.50 (30), 8.9 ± 0.1 min
pN3F
NRPS-1
MS2: 895.51 (30), 9.0 ± 0.1 min
OPrY
NRPS-1
MS2: 933.50 (30b), 9.8 min
OPrY
ΔC3H2
NRPS-1
MS2: 895.54 (32), 9.6 min
no supplementation
NRPS-1
MS2: 895.56 (32), 9.6 min
pN3F
NRPS-1
MS2: 895.57 (32), 9.7 ± 0.1 min
OPrY
m/z

**Figure S35.** Plasmid-based heterologous expression of *NRPS-1* in *E. coli* DH10B::*mtaA*. **A.** Schematic depiction of NRPS-1 and structures of produced peptides **30**, **30b**, and **32**. A-domain specificity indicated by the amino acid one-letter code. **B.** Extracted ion chromatograms (EIC) of precursor-directed biosynthesis with NRPS-1 supplementing with 1 mM *p*$N_3$F (**12**) or *O*PrY (**13**). C. $MS^2$ fragmentation spectra of **30**, **30b**, and **32** produced by NRPS-1.

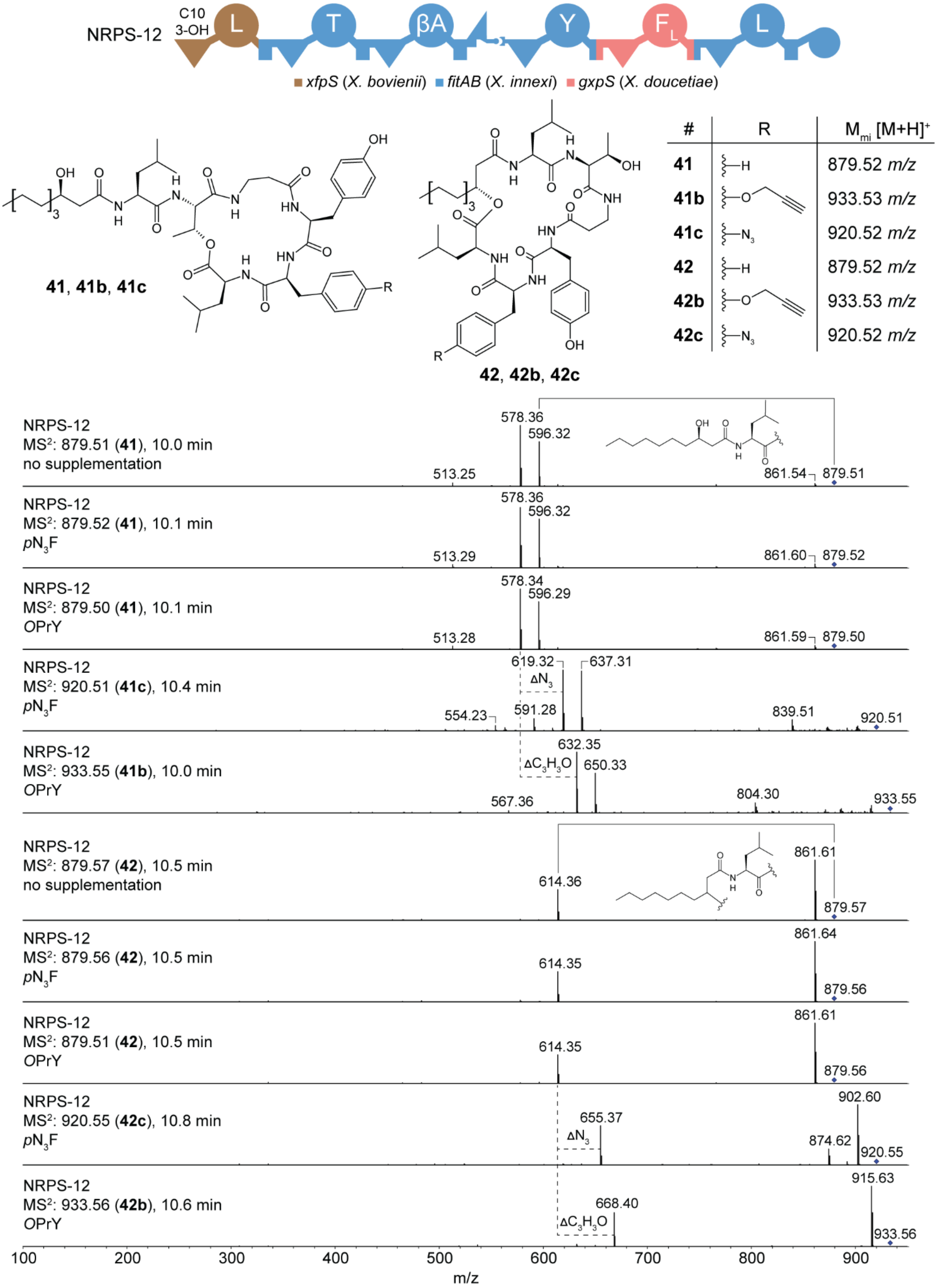

**Figure S36.** MS$^2$ fragmentation spectra corresponding to Figure 4 of compounds **41-41c** and **42-42c** generated through precursor-directed biosynthesis by heterologous expression of

*NRPS-12* in *E. coli* DH10B::*mtaA*. For production, 1 mM *p*N3F (**7**) or 1 mM *O*PrY (**14**) were added to the production media XPPM.

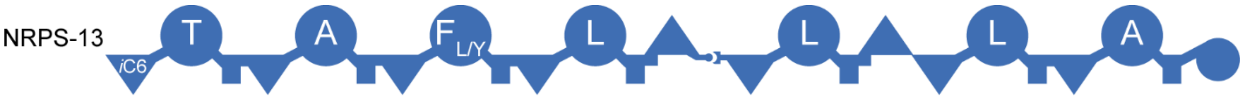

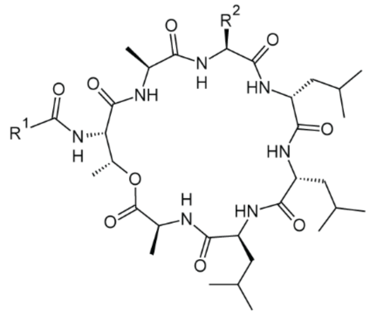

**43-49**

| | $R^1$ | $R^2$ | $M_{mi}$ $[M+H]^+$ |
|---|---|---|---|
| **43** | C5 | Bn | 828.52 *m/z* |
| **44** | C5 | *i*Bu | 794.54 *m/z* |
| **45** | C3 | Bn | 766.51 *m/z* |
| **46** | C3 | *i*Bu | 800.49 *m/z* |
| **47** | C5 | Bn(4-OH) | 844.52 *m/z* |
| **48** | C1 | Bn | 772.46 *m/z* |
| **49** | C1 | *i*Bu | 738.48 *m/z* |

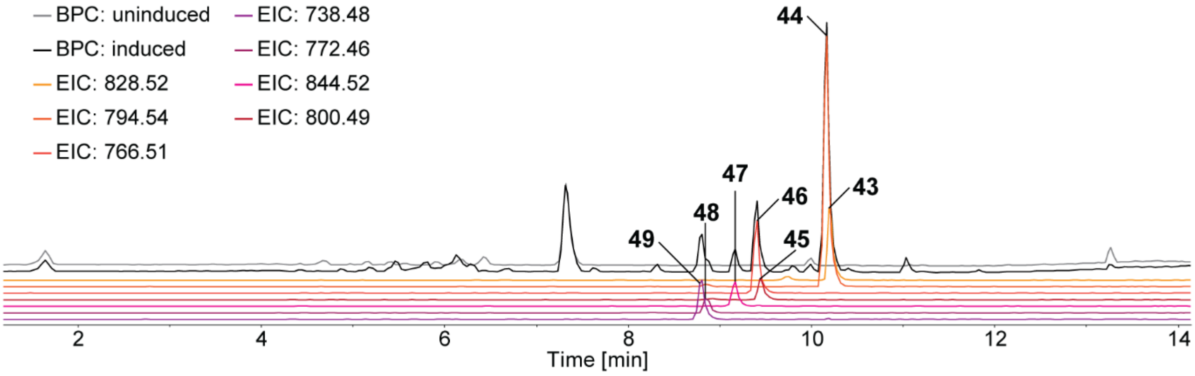

**Figure S37.** Plasmid-based heterologous expression of *txlAB* in *E. coli* DH10B::*mtaA*. Schematic depiction of NRPS-13 and structures of produced taxlllaid derivatives **43-49**. Retention times are marked in the respective base peak (BPC) and extracted ion chromatograms (EIC). A-domain specificity indicated by the amino acid one-letter code.

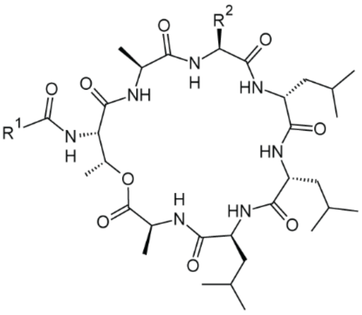

**43-49**

| | R¹ | R² | $M_{mi}$ [M+H]⁺ |
|---|---|---|---|
| **43** | C5 | Bn | 828.52 *m/z* |
| **44** | C5 | *i*Bu | 794.54 *m/z* |
| **45** | C3 | Bn | 766.51 *m/z* |
| **46** | C3 | *i*Bu | 800.49 *m/z* |
| **47** | C5 | Bn(4-OH) | 844.52 *m/z* |
| **48** | C1 | Bn | 772.46 *m/z* |
| **49** | C1 | *i*Bu | 738.48 *m/z* |

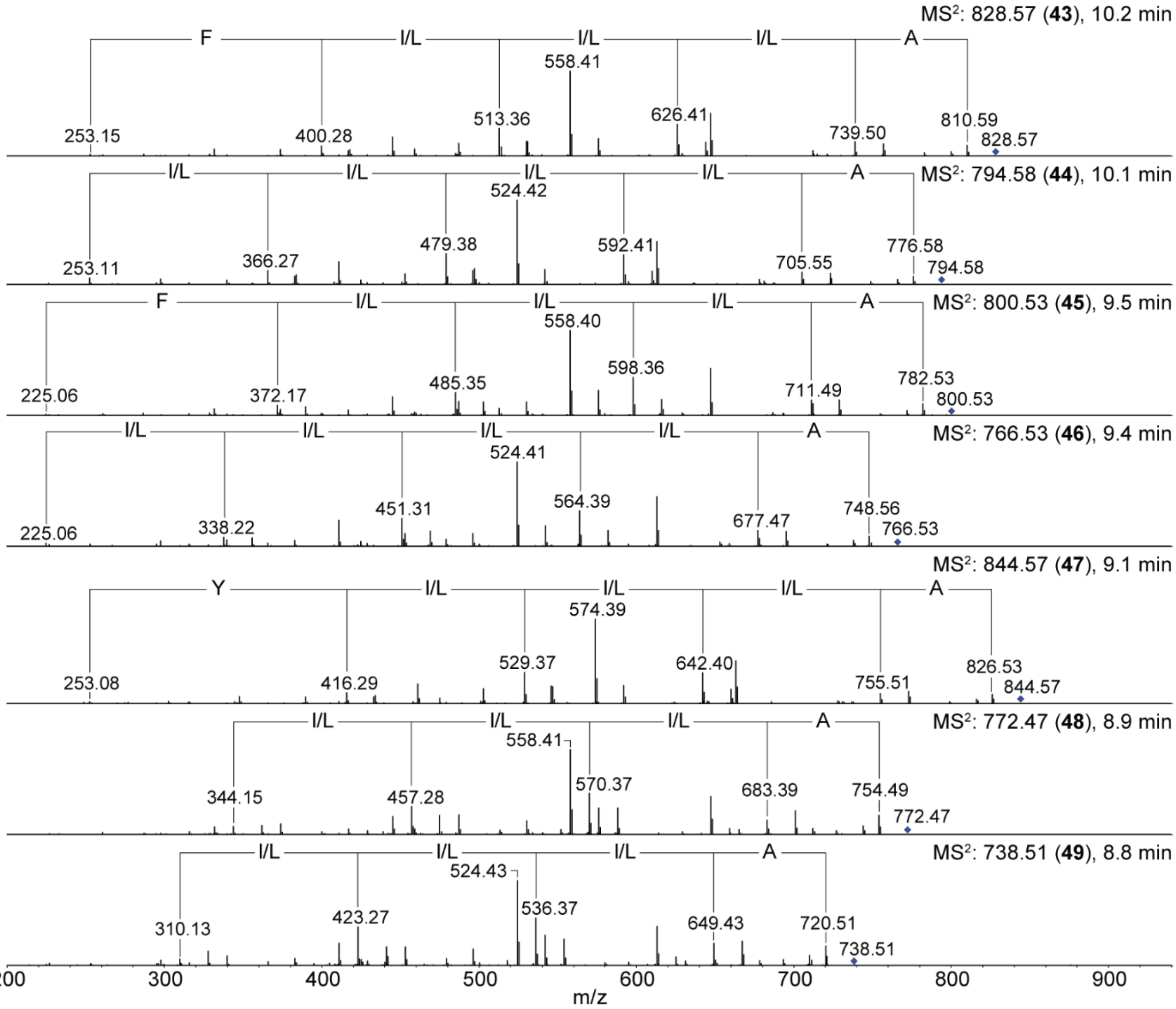

**Figure S38.** $MS^2$ fragmentation spectra corresponding to Figure S37 of taxIIIaid derivatives **43-49** produced by plasmid-based heterologous expression of *NRPS-13*.

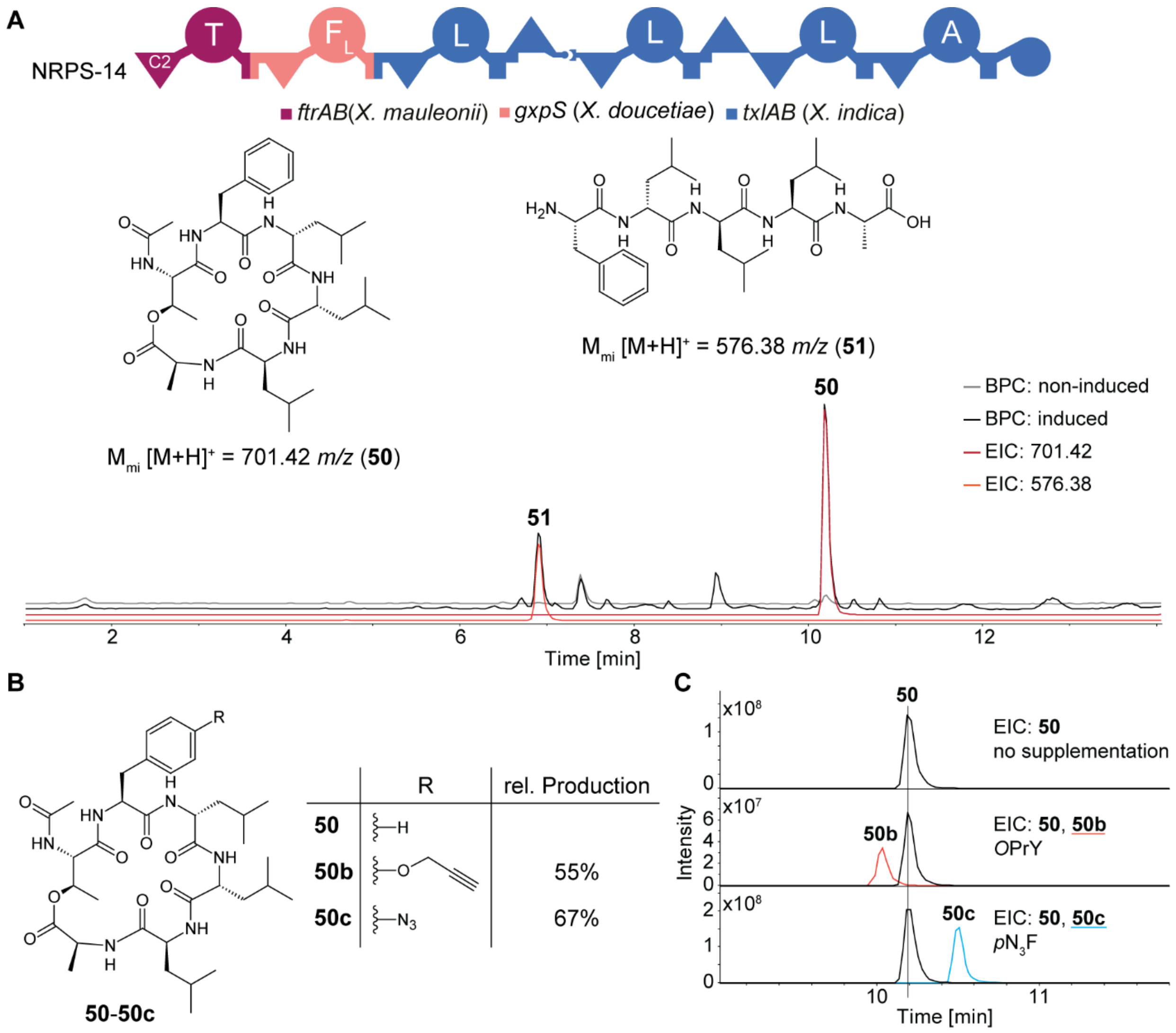

**Figure S39.** Production of *E. coli* DH10B::*mtaA* heterologously expressing *NRPS-14*. **A.** Schematic depiction of NRPS-14 and the structures of produced compounds **50** and **51**. Retention times are marked in the respective base peak (BPC) and extracted ion chromatograms (EIC). A-domain specificity indicated by the amino acid one-letter code. **B.** Structures of compounds **50-50c** generated by precursor-directed biosynthesis with NRPS-14. **C.** EIC of the production of **50-50c** in XPPM with and without the supplementation of 1 mM *p*$N_3$F (**12**) or 1 mM *O*PrY (**13**).

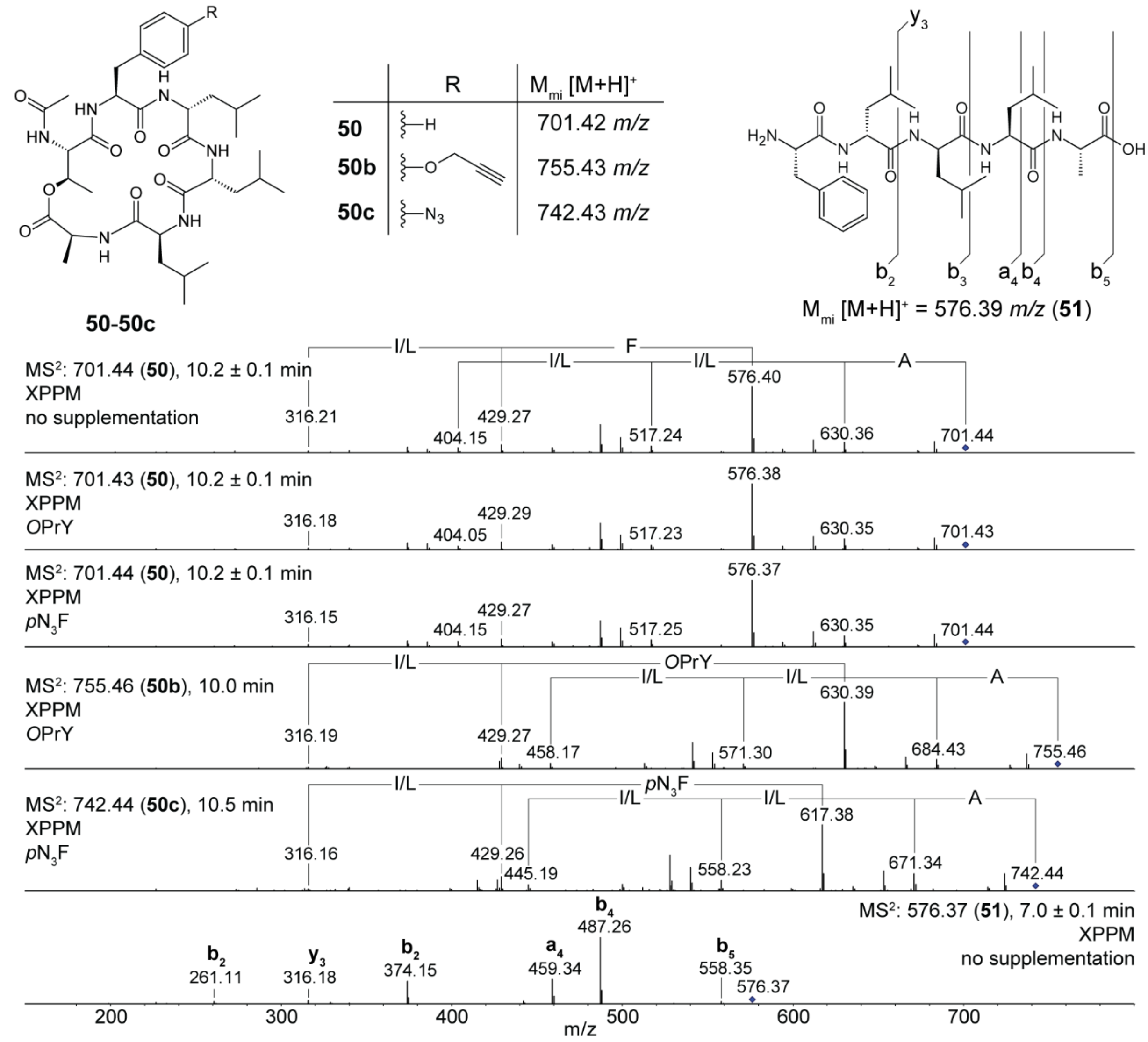

**Figure S40.** $MS^2$ fragmentation spectra of compounds corresponding to Figure S39 produced by plasmid-based heterologous expression of *NRPS-14* in *E. coli* DH10B::*mtaA*. Fragmentation spectra of compounds **50-50c** generated through precursor-directed biosynthesis with addition of 1 mM *p*N3F (**12**) or 1 mM *O*PrY (**13**) to the production medium XPPM and the fragmentation spectrum of **51**.

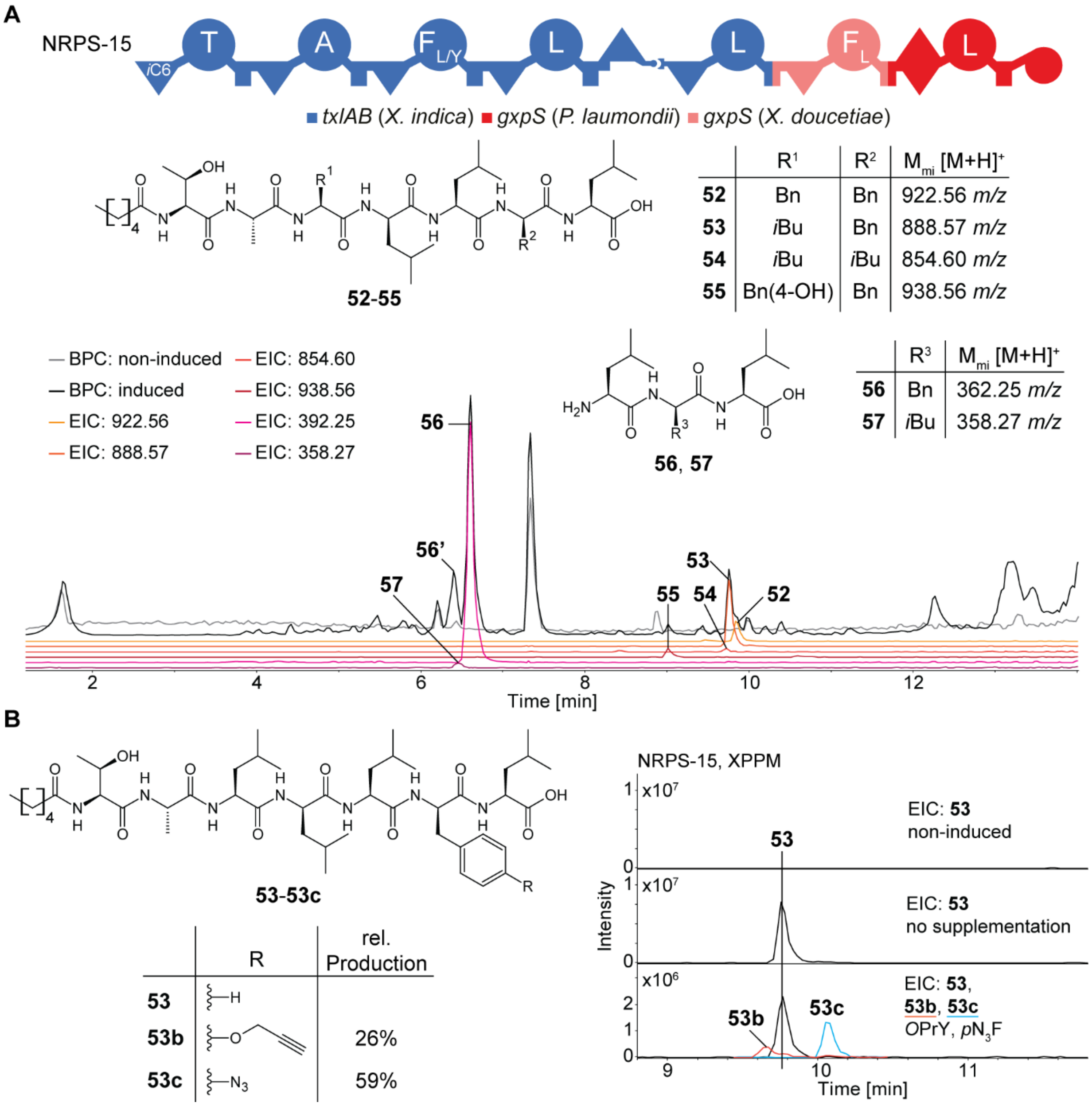

**Figure S41.** Production of *E. coli* DH10B::*mtaA* heterologously expressing *NRPS-15*. **A.** Schematic depiction of NRPS-15 and the structures of produced compounds **52-57**. Retention times are marked in the respective base peak (BPC) and extracted ion chromatograms (EIC). A-domain specificity indicated by the amino acid one-letter code. **B.** Structures of compounds **53-53c** generated by precursor-directed biosynthesis with NRPS-15. EIC of their production in XPPM with and without simultaneous supplementation of $pN_3F$ (**12**) and *O*PrY (**13**).

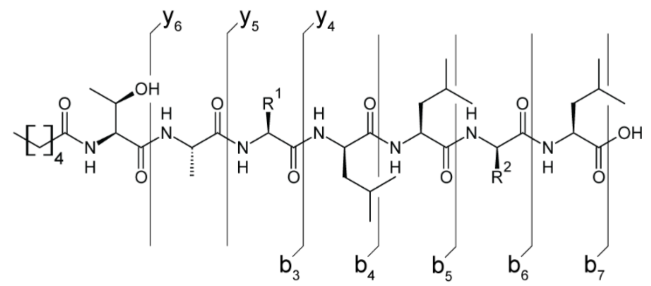

**52-55**

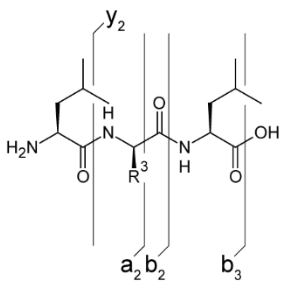

**56**, **57**

| | R[1] | R[2] | $M_{mi}$ [M+H]+ |
|---|---|---|---|
| **52** | Bn | Bn | 922.56 *m/z* |
| **53** | *i*Bu | Bn | 888.57 *m/z* |
| **54** | *i*Bu | *i*Bu | 854.60 *m/z* |
| **55** | Bn(4-OH) | Bn | 938.56 *m/z* |

| | R[3] | $M_{mi}$ [M+H]+ |
|---|---|---|
| **56** | Bn | 362.25 *m/z* |
| **57** | *i*Bu | 358.27 *m/z* |

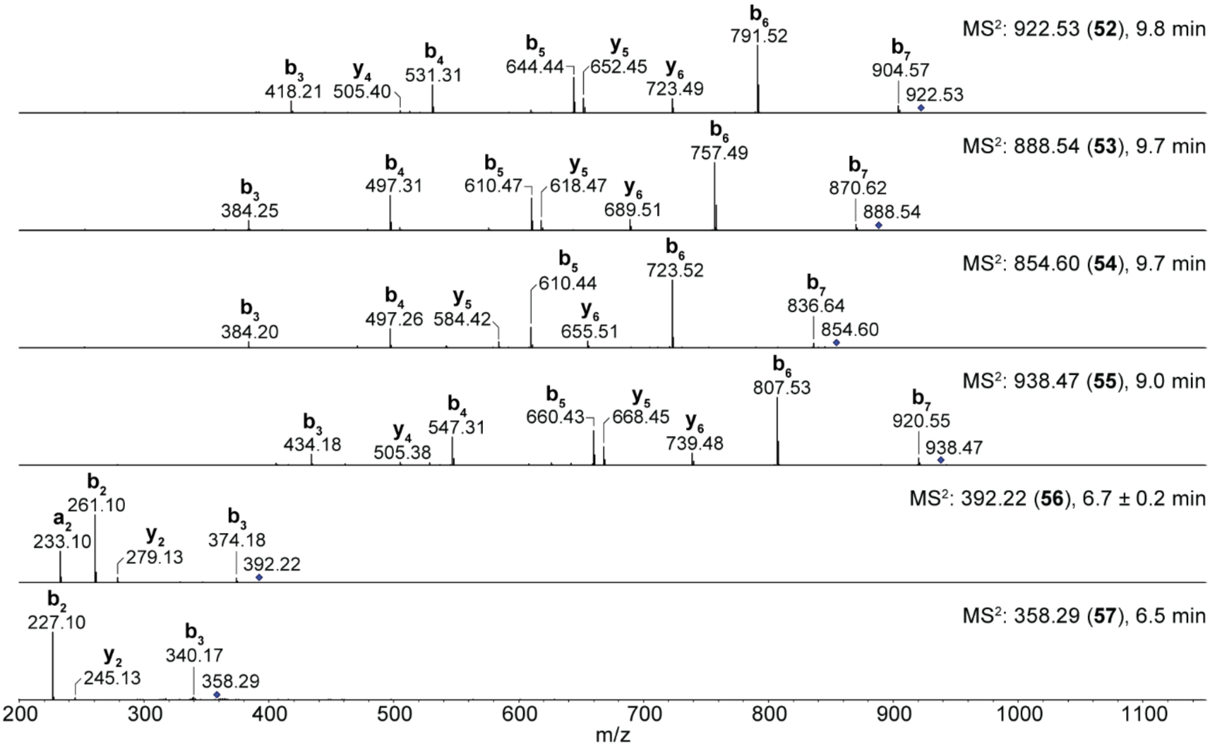

**Figure S42.** $MS^2$ fragmentation spectra corresponding to Figure S41A of compounds **52-57** produced by plasmid-based heterologous expression of *NRPS-15* in *E. coli* DH10B::*mtaA*.

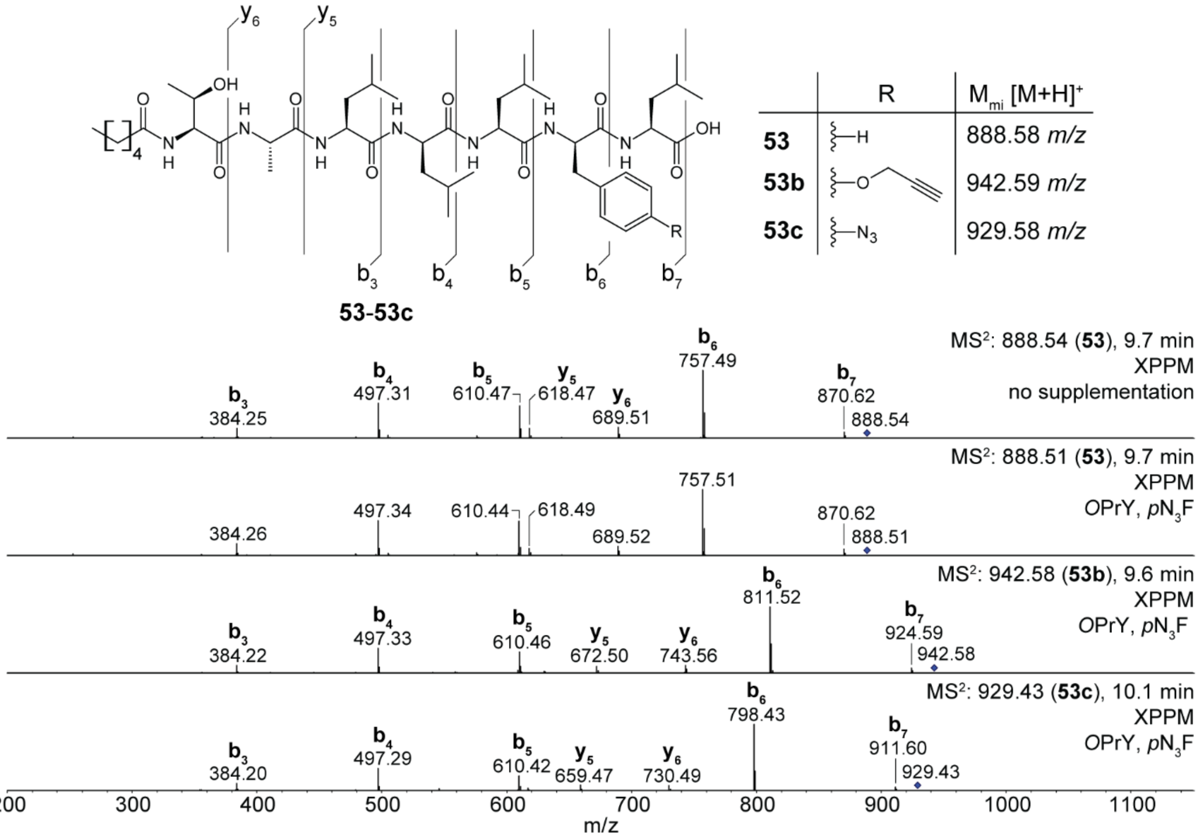

**Figure S43.** $MS^2$ fragmentation spectra corresponding to Figure S41B of compounds **53-53c** generated through precursor-directed biosynthesis by heterologous expression of *NRPS-15* in *E. coli* DH10B::*mtaA*. For production, 1 mM *p*$N_3$F (**12**) and 1 mM *O*PrY (**13**) were simultaneously added to the production medium XPPM.

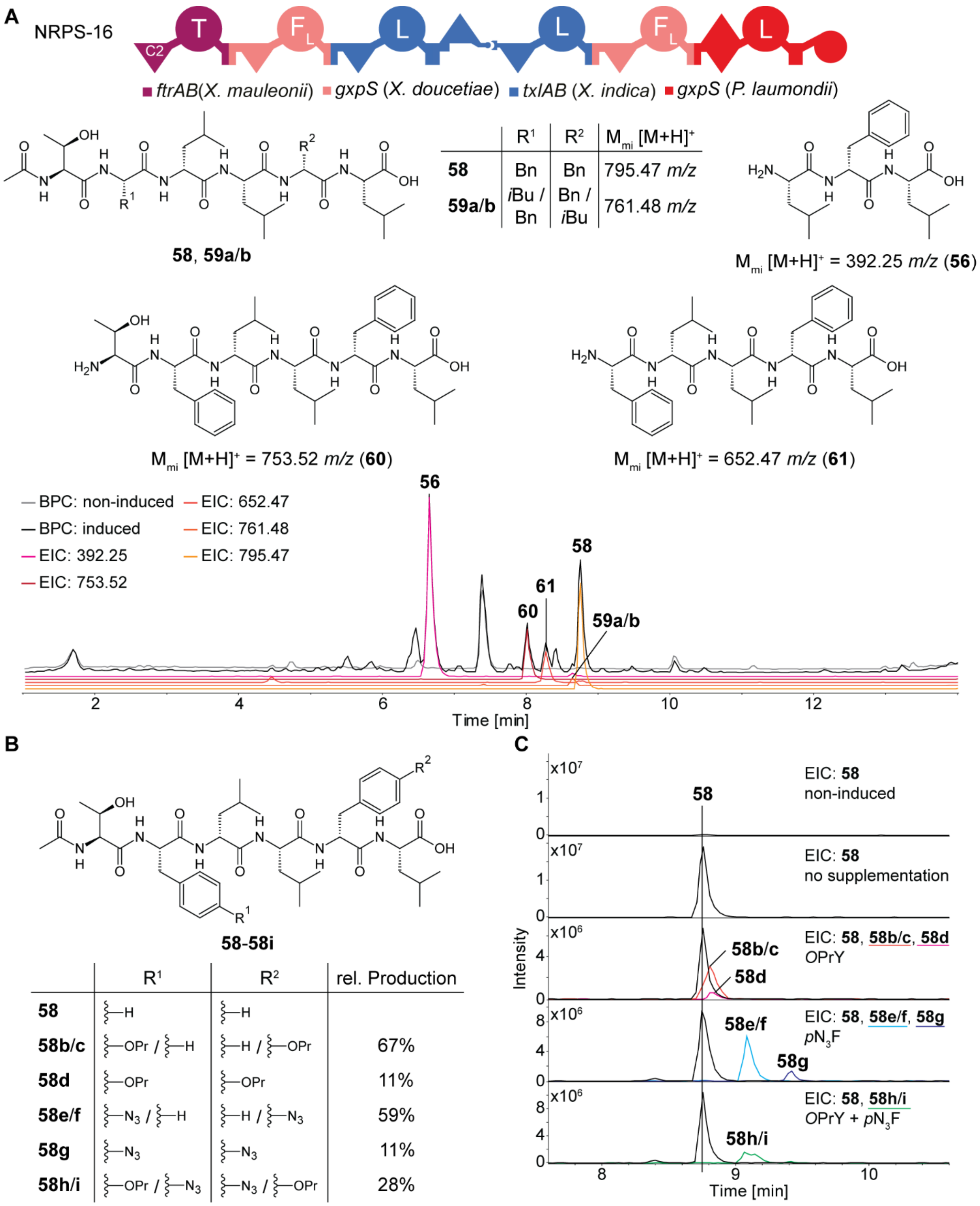

**Figure S44.** Production of *E. coli* DH10B::*mtaA* heterologously expressing *NRPS-16*. **A.** Schematic depiction of NRPS-16 and the structures of produced compounds **56** and **58-61**. Retention times are marked in the respective base peak (BPC) and extracted ion chromatograms (EIC). A-domain specificity indicated by the amino acid one-letter code. **B.** Structures of compounds **58** and **58b-i** generated by precursor-directed biosynthesis with NRPS-16. **C.** EIC of the production of **58** and **58b-i** in XPPM with and without the supplementation of 1 mM *p*$N_3$F (**12**) or/and 1 mM *O*PrY (**13**).

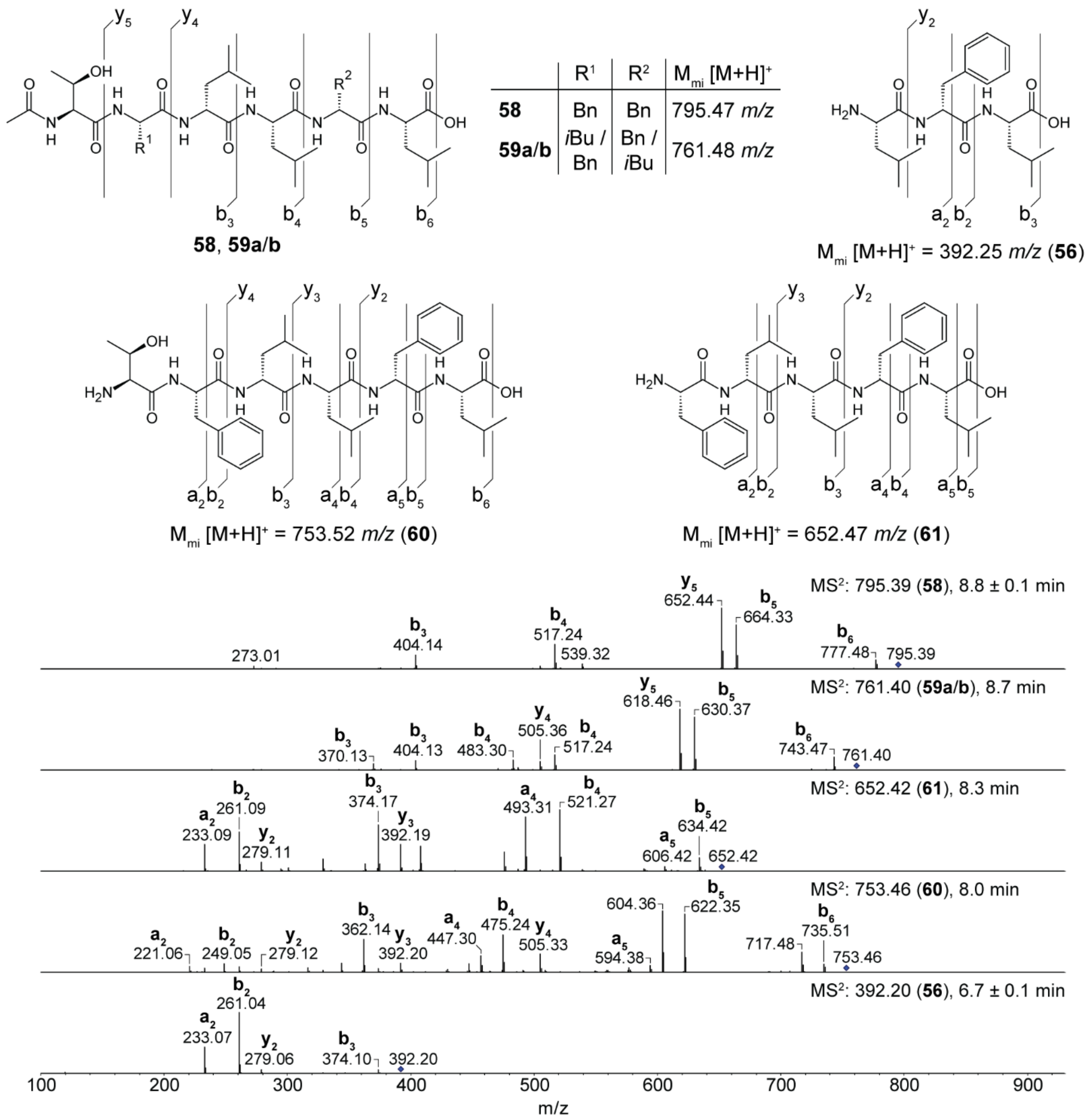

**Figure S45.** $MS^2$ fragmentation spectra corresponding to Figure S44A of compounds **56** and **58-61** produced by plasmid-based heterologous expression of *NRPS-16* in *E. coli* DH10B::*mtaA*.

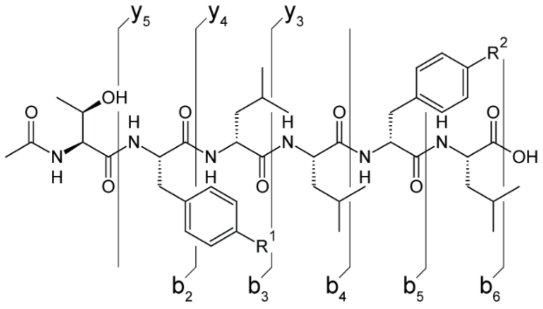

**58**-**58i**

| | $R^1$ | $R^2$ | $M_{mi}$ $[M+H]^+$ |
|---|---|---|---|
| **58** | –H | –H | 795.47 *m/z* |
| **58b/c** | –OPr / –H | –H / –OPr | 849.48 *m/z* |
| **58d** | –OPr | –OPr | 903.49 *m/z* |
| **58e/f** | $–N_3$ / –H | –H / $–N_3$ | 836.47 *m/z* |
| **58g** | $–N_3$ | $–N_3$ | 877.47 *m/z* |
| **58h/i** | –OPr / $–N_3$ | $–N_3$ / –OPr | 890.48 *m/z* |

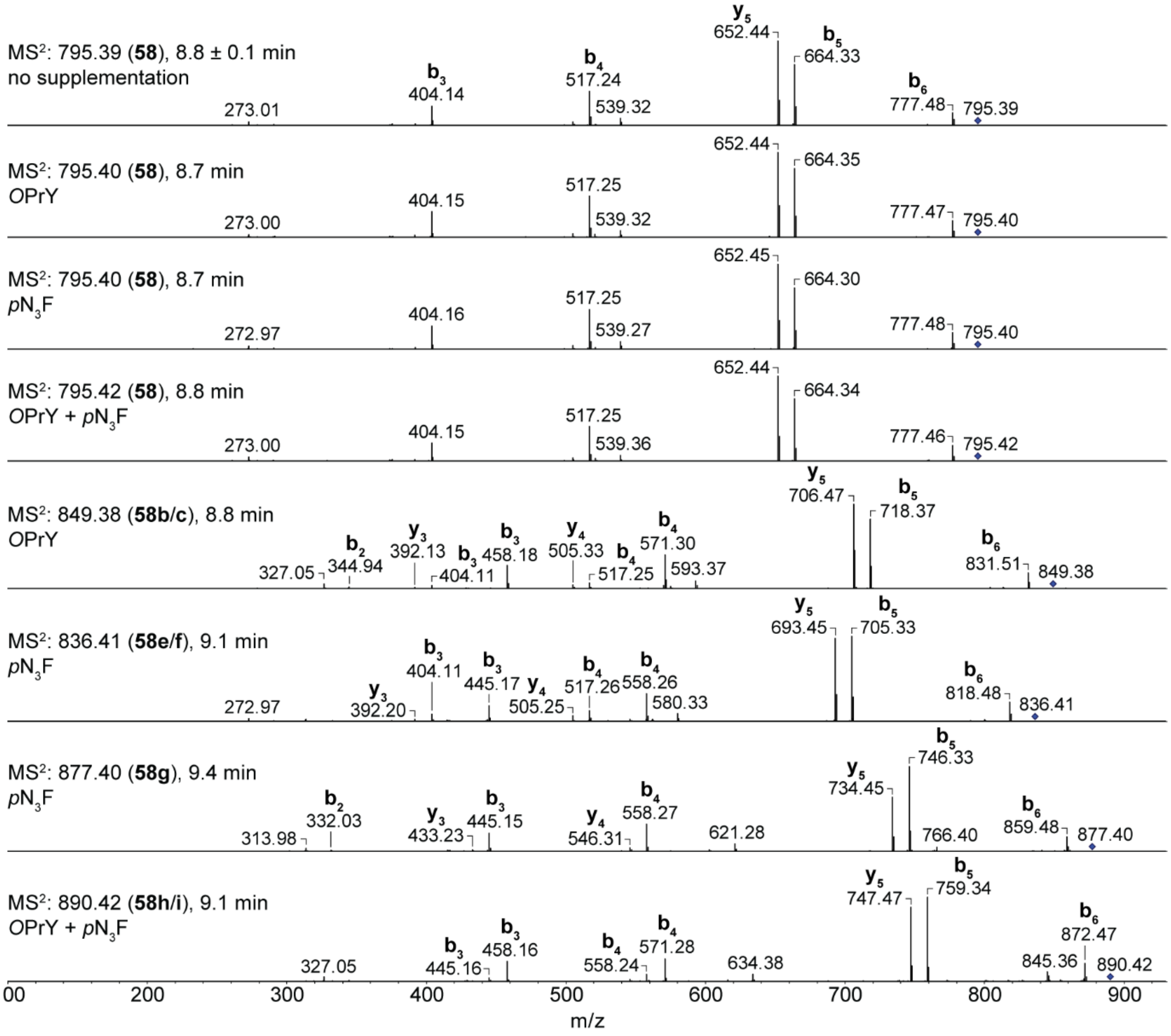

**Figure S46.** $MS^2$ fragmentation spectra corresponding to Figure S44B/C of compounds **58** and **58b-i** generated through precursor-directed biosynthesis by heterologous expression of *NRPS-16* in *E. coli* DH10B::*mtaA*. For production, 1 mM *p*N3F (**12**) and/or 1 mM *O*PrY (**13**) were simultaneously added to the production media. For **58d**, no $MS^2$ fragmentation spectra could be obtained.

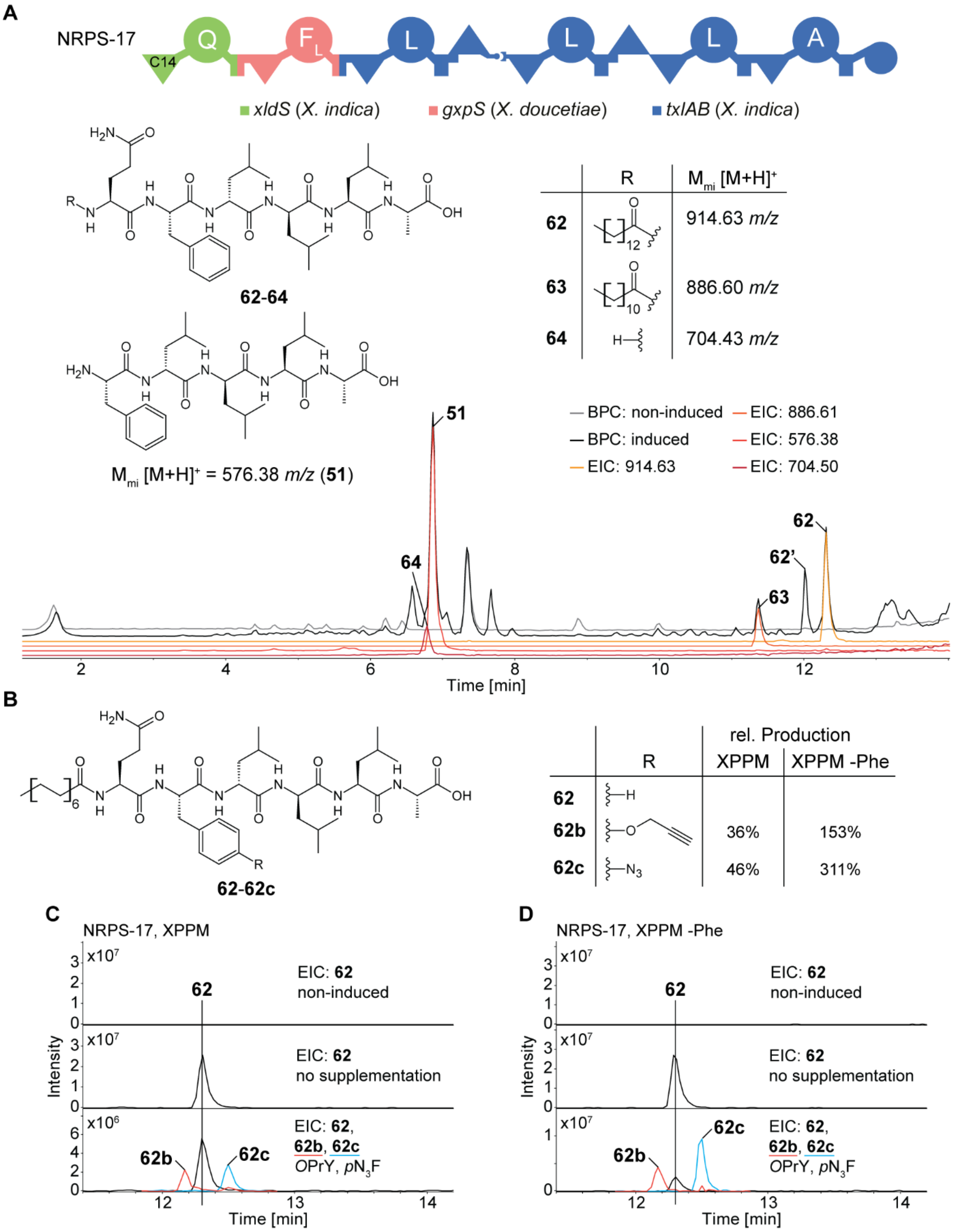

**Figure S47.** Production of *E. coli* DH10B::*mtaA* heterologously expressing *NRPS-17*. **A.** Schematic depiction of NRPS-17 and the structures of produced compounds **51** and **62-64**. Retention times are marked in the respective base peak (BPC) and extracted ion chromatograms (EIC). **62'** is a *C*-terminal glycerol ester of **62**. A-domain specificity indicated by the amino acid one-letter code. **B.** Structures of compounds **62-62c** generated by precursor-directed biosynthesis with NRPS-17. EIC of the production of **62-62c** in XPPM (**C**) and XPPM without phenylalanine (-Phe, **D**) with and without simultaneous supplementation of 1 mM *p*$N_3$F (**12**) and 1 mM *O*PrY (**13**).

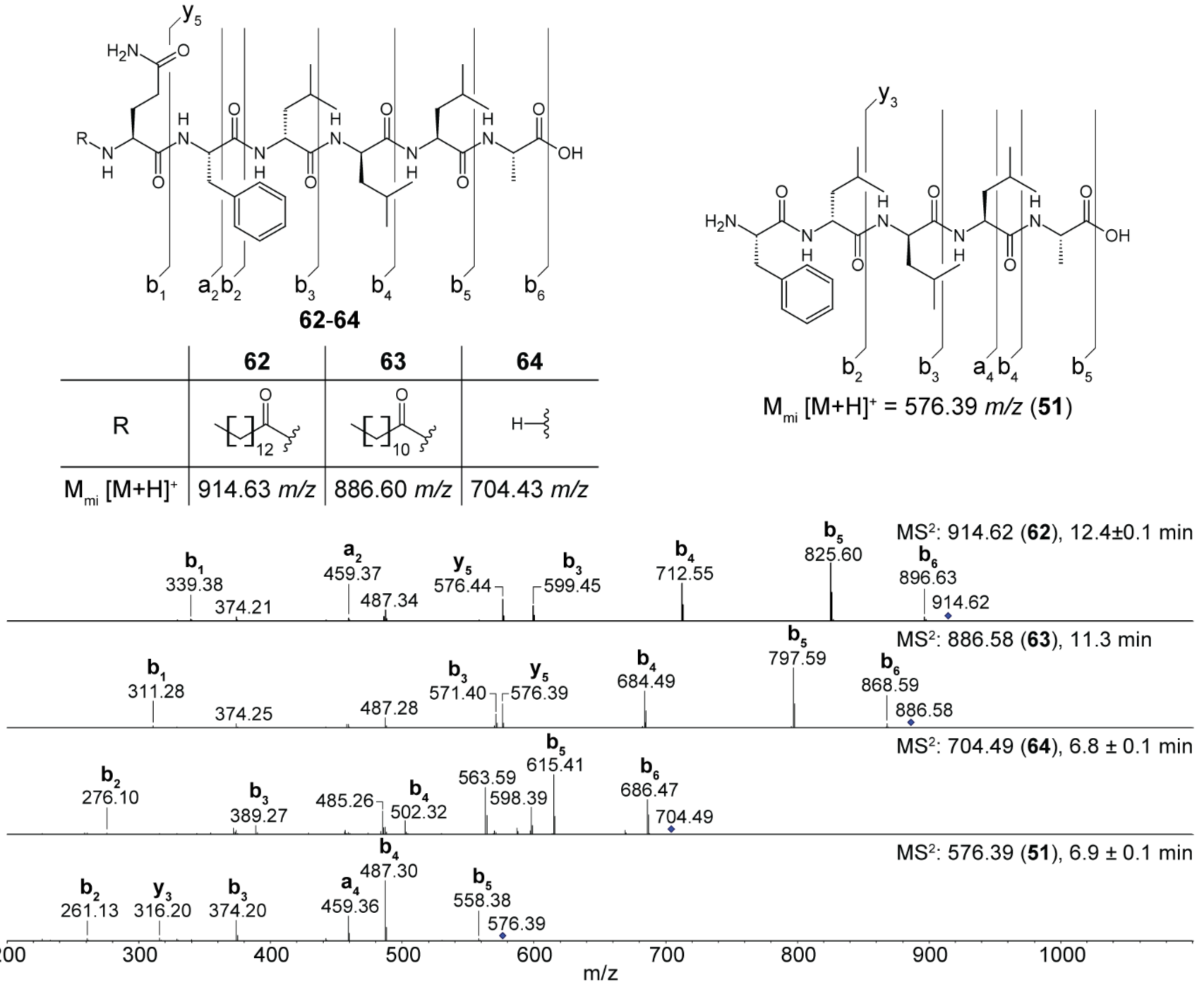

**Figure S48.** MS$^2$ fragmentation spectra corresponding to Figure S47A of compounds **60-62** and **49** produced by plasmid-based heterologous expression of *NRPS-17* in *E. coli* DH10B::*mtaA*.

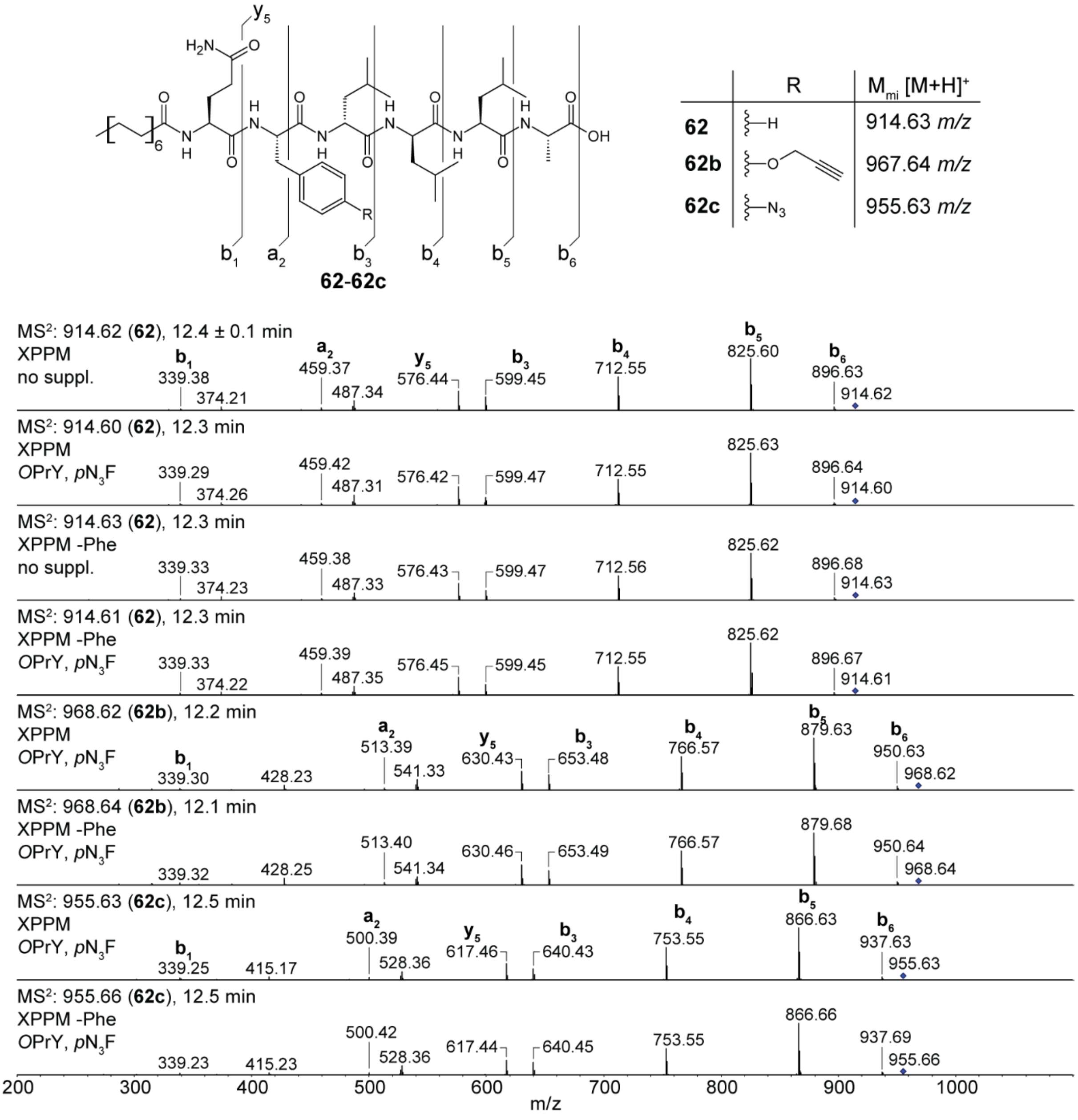

**Figure S49.** MS$^2$ fragmentation spectra corresponding to Figure S47B-D of compounds **62**-**62c** generated through precursor-directed biosynthesis by heterologous expression of *NRPS-17* in *E. coli* DH10B::*mtaA*. For production, 1 mM *p*N3F (**12**) and 1 mM *O*PrY (**13**) were simultaneously added to the production media, XPPM and XPPM without phenylalanine (XPPM -Phe).

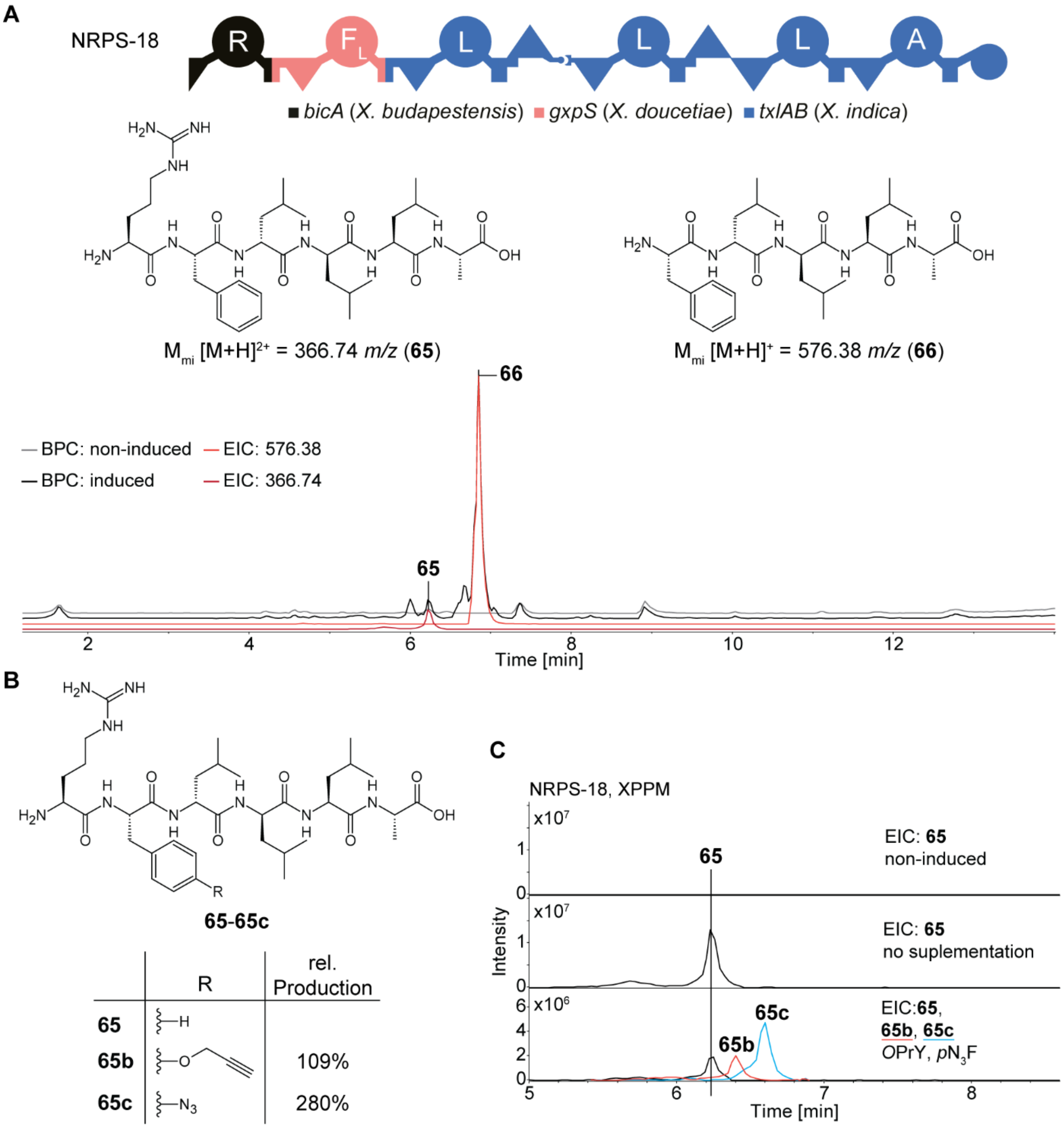

**Figure S50.** Production of *E. coli* DH10B::*mtaA* heterologously expressing *NRPS-18*. **A.** Schematic depiction of NRPS-18 and the structures of produced compounds **65** and **66**. Retention times are marked in the respective base peak (BPC) and extracted ion chromatograms (EIC). A-domain specificity indicated by the amino acid one-letter code. **B.** Structures of compounds **65-65c** generated by precursor-directed biosynthesis with NRPS-29. **C.** EIC of the production of **65-65c** in XPPM with and without simultaneous supplementation of 1 mM *p*$N_3$F (**12**) and 1 mM *O*PrY (**13**).

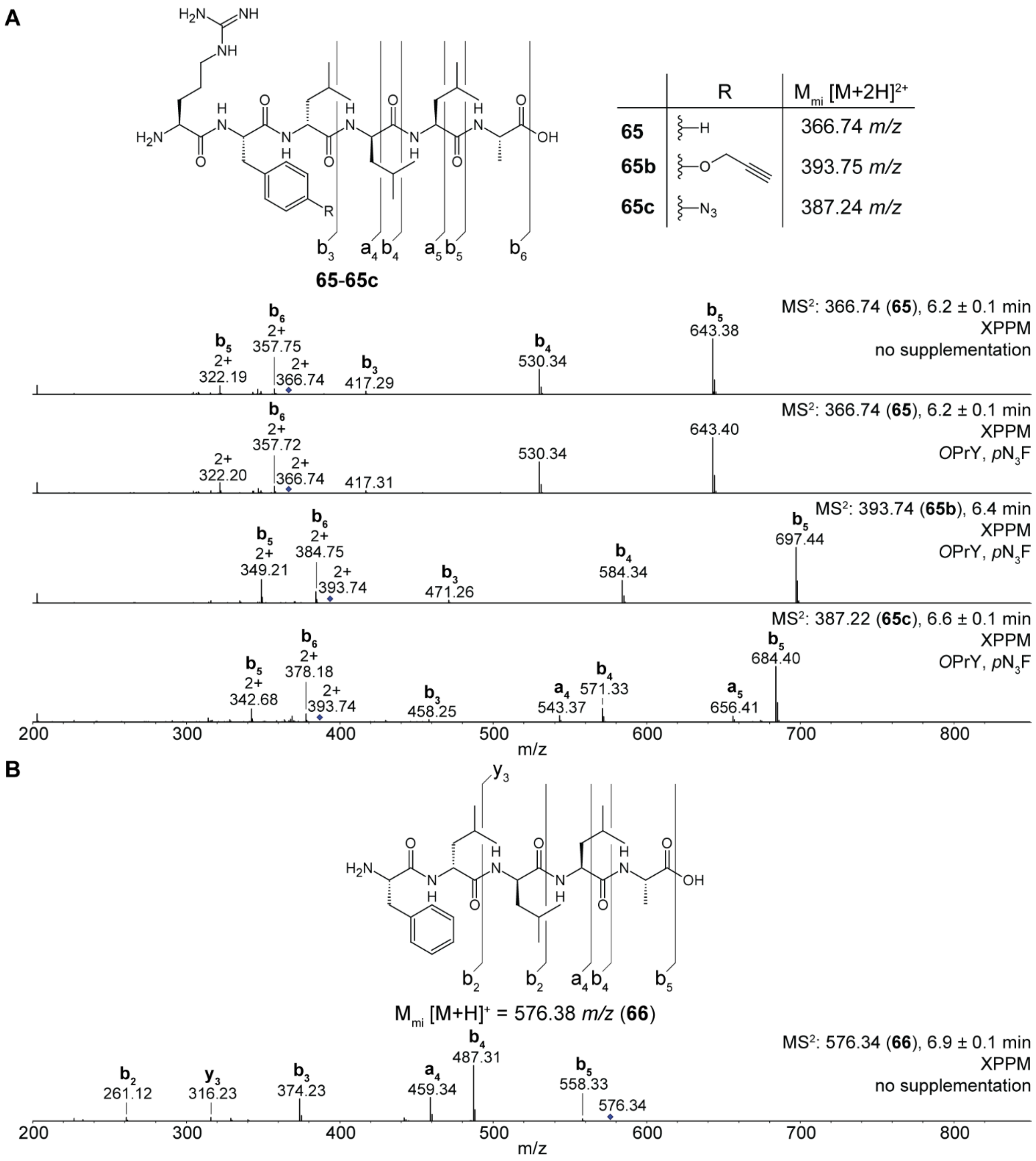

**Figure S51.** $MS^2$ fragmentation spectra of compounds corresponding to Figure S50 produced by plasmid-based heterologous expression of *NRPS-18* in *E. coli* DH10B::*mtaA*. **A.** Fragmentation spectra of compounds **65**-**65c** generated through precursor-directed biosynthesis with simultaneous addition of 1 mM *p*$N_3$F (**12**) and 1 mM *O*PrY (**13**) to the production medium XPPM. **B.** Fragmentation spectrum of **66**.

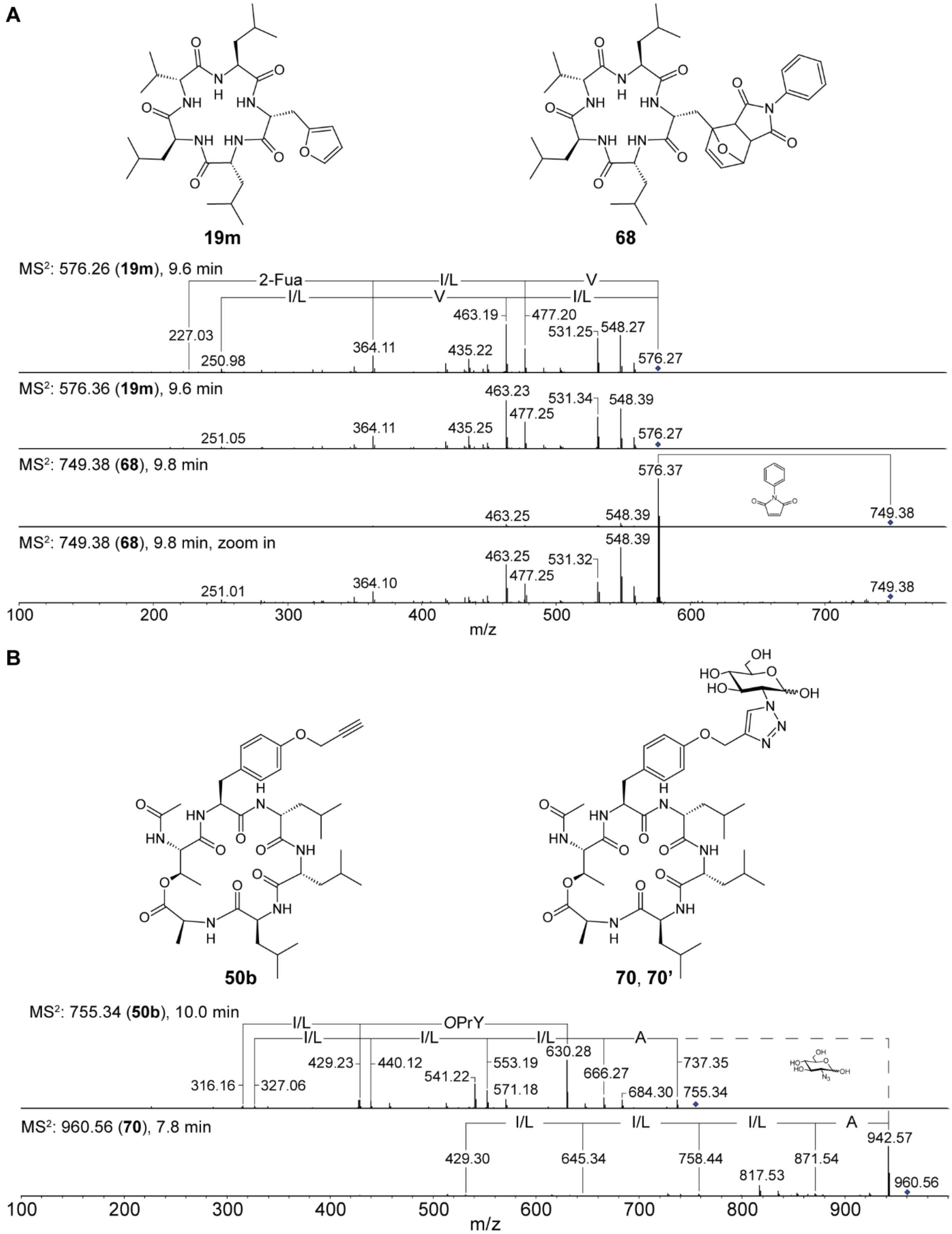

**Figure S52.** $MS^2$ fragmentation spectra corresponding to Figure 5 of compounds **19m**, **50b**, **68**, and **70** generated through precursor-directed biosynthesis and chemical modification, respectively. **A. 19m** was generated expressing *GxpS* in *X. doucetiae* supplementing with 2-Fua (**7**). **68** was generated by Diels-Alder reaction with *N*-phenylmaleimide (**67**) in an extract containing **19m**. **B. 50b** was generated by heterologous expression of *NRPS-14* in *E. coli* DH10B::*mtaA* supplementing *O*PrY (**13**). For glycosylation with $N_3$Gluc (**67**) in an extract containing **50b**, copper-catalyzed azide-alkyne cycloaddition was performed. **70** and **70'** resemble α- and β-anomers of the glucopyranose. No MS$^2$ spectrum was obtained for **70'**.

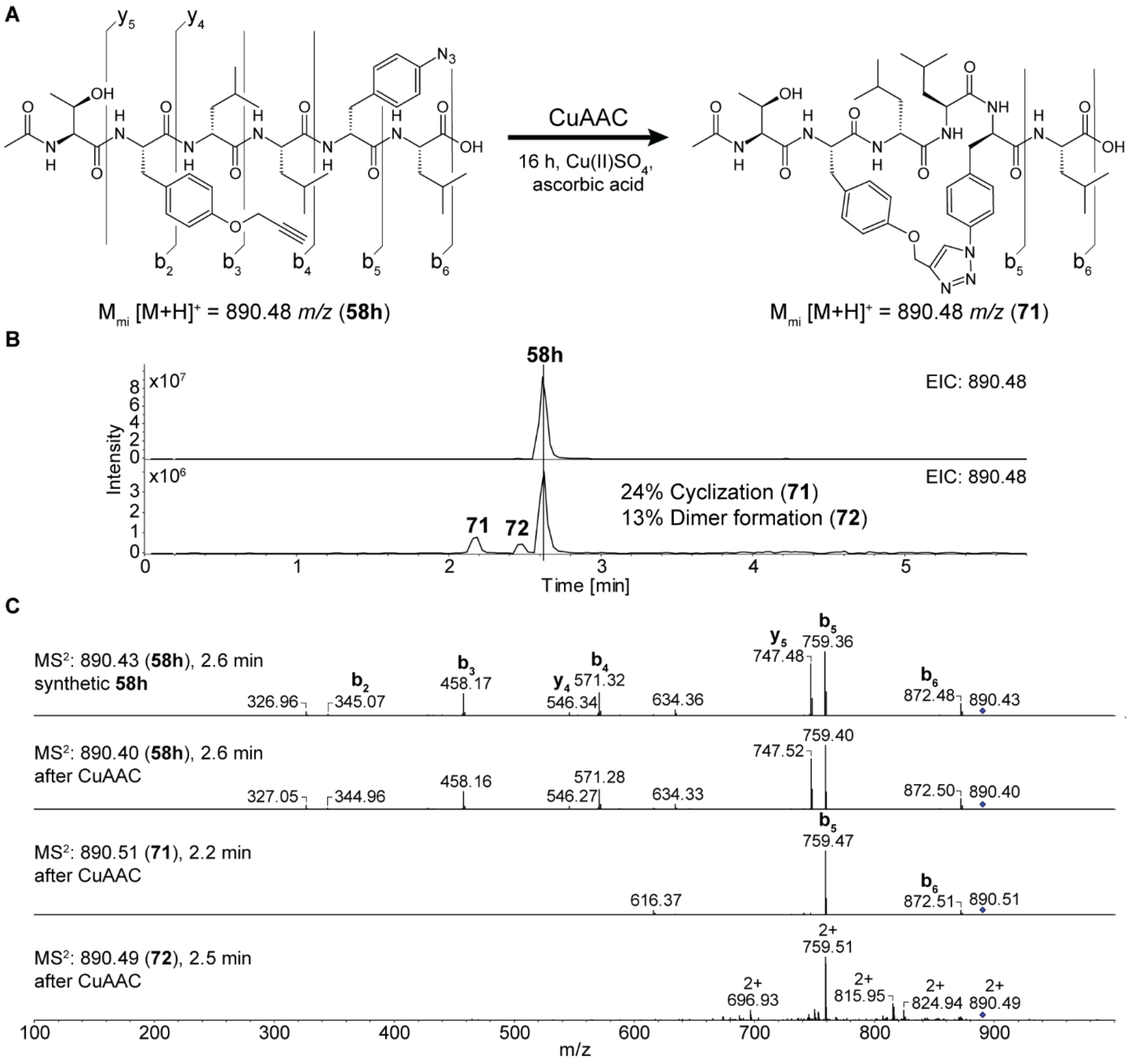

**Figure S53.** Intra-molecular copper-catalyzed azide-alkyne cycloaddition. **A.** Structure of synthetic peptide **58h** and cyclized **71**. **58h** can be biosynthetically produced by precursor-directed biosynthesis using NRPS-16 (Figure S44B, Figure S46). **B.** Extracted ion chromatograms (EIC) of **58h**, **71**, and **72** (dimer of **58h**). **C.** MS$^2$ fragmentation spectra of **58h**, **71**, and **72**.
